# Targeted Photoconvertible BODIPYs Based on Directed Photooxidation Induced Conversion for Applications in Photoconversion and Live Super Resolution Imaging

**DOI:** 10.1101/2023.07.28.550940

**Authors:** Lazare Saladin, Victor Breton, Thibault Lequeu, Pascal Didier, Lydia Danglot, Mayeul Collot

## Abstract

Photomodulable fluorescent probes are drawing an increasing attention due to their applications in advanced bioimaging. Whereas photoconvertible probes can be advantageously used in tracking, photoswitchable probes constitute key tools for single molecule localization microscopy to perform super resolution imaging. Herein we shed light on a red and far-red BODIPY, namely BDP-576 and BDP-650 possessing both properties of conversion and switching. Our study demonstrates that theses pyrrolyl-BODIPYs respectively convert towards typical green- and red-emitting BODIPYs that are perfectly adapted to microscopy. We also showed that these pyrrolyl-BODIPYs undergo Directed Photooxidation Induced Conversion, a photoconversion mechanism that we recently introduced and where the pyrrole moiety plays a central role. These unique features were used to develop targeted photoconvertible probes towards different organelles or subcellular units (plasma membrane, mitochondria, nucleus, actin, Golgi apparatus, *etc*.) using chemical targeting moieties and Halo tag. We notably showed that BDP-650 could be used to track intracellular vesicles over more than 20 minutes in two color imaging with laser scanning confocal microscopy demonstrating its robustness. The switching properties of these photoconverters were studied at the single molecule level and were then successfully used in live Single Molecule Localization Microscopy in epithelial cells and neurons. Both membrane and mitochondria targeted probes could be used to decipher membrane 3D architecture and mitochondria dynamics at the nanoscale. This study builds a bridge between the photoconversion and photoswitching properties of probes undergoing directed photooxidation and shows the versatility and efficacy of this mechanism in live advanced imaging.

## Introduction

Photomodulable fluorescent probes able to modify their properties upon light irradiation are gaining an increasing attention due to their appealing properties. Whereas those which undergo a cleavage of a chemical bond enable to activate a biomolecules (also called photocages) or release a cargo,^1^ probes that change their photophysical properties (*e*.*g*. fluorescence intensity, blinking, maximum emission wavelength) are gaining interest due to their applications in advanced microscopic bioimaging.^2 3 4^ Among this latter type of probes, those which photobleach and rapidly turn off their fluorescence upon irradiation can be advantageously used in fluorescence recovery after photobleaching experiment (FRAP) to study the diffusion of molecules.^5 6^ In contrast, photoactivatable probes that turn on their fluorescence emission,^7 10^ mainly find applications in single molecule tracking,^9 10^ and super resolution microscopy based on PhotoActivated Localization Microscopy (PALM). ^2 3 11 12 13 14^ Photoswitching probes are dyes that reversibly switch from a non-emissive or emissive specie towards another emissive one.^7 15 16 17^ They are particularly adapted to single molecule localization microscopy (SMLM) on live samples as the localization of their multiple spontaneous blinking events enables the reconstruction of images with enhanced resolution.^18 19 20 21 22^ Although several OFF-ON photoswitchers have been described, ^2 15 23^ those able to switch from an emissive form to another are less frequent.^18 19 24 25^

Finally, photoconvertible probes (herein restricted to the visible range 400 nm ≤ λ_Em max_ ≤ 700 nm) are emissive dyes that upon light irradiation convert towards another emissive form of a different colour through an irreversible mechanism. These probes are suitable to unambiguously track labeled object and study dynamic phenomenon in living samples with large spatiotemporal scales.^26 27 28^ Despite their limitations, photoconvertible fluorescent proteins (PCFP),^29 30^ including Kande,^31^ Dronpa,^32^ Dendra,^33^ and those of the EosFP family^34^ dominate in bioimaging applications. Indeed, genetically encoded PCFP can only label proteins, require a transfection step and lead to heterogeneously labeled cells. Moreover, UV irradiation is often required to trigger the conversion, which is detrimental for cells. Conversely, small molecular probes provide an homogeneous staining and benefit from accessible tuning and improvement of their properties through chemical modifications.^35 36^ For these reasons, chemists recently focused their efforts on the development of Dual-Colour Photoconvertible Fluorophores (DCPF). Inspired by photoconvertible fluorescent proteins, DCPF providing a bathochromic shift upon conversion were developed. Chenoweth and co-workers proposed an approach based photo-cyclodehydrogenation of diazaxanthilidene,^37 38^ and showed their application in bioimaging using a targeted version.^39^ Similarly, Tang’s group made photoconverters based on a photooxidative dehydrogenation mechanism.^40^ In a different approach, Raymo’s group developed several BODIPYs photoconverters based on photoinduced disconnection of an spiro-oxazine.^12 28 41^ Recently photoextrusion of sulfur monoxide (SO) has been proposed as an elegant approach to photoconvert a sulfoxide-bridged dimeric BODIPY towards a red-shifted conjugated dimeric BODIPY.^42^ Inversely, DCPFs undergoing a hypsochromic shift upon conversion, a phenomenon also called “photoblueing”,^43^ were also proposed. Although empirical approaches consisting in studying the potential emitting photoproducts after photobleaching have been investigated,^44 45 46^ more rational approaches have been reported. Cyanine dyes have drawn a special attention as they were shown to undergo a phototruncation under irradiation, leading to the generation of blue-shifted cyanine analogues.^26 27 47 48 43^ Hell’s group also reported that rhodamines could undergo photooxidative dealkylation leading to hyposchromic shifts.^49^

Although these works brought new insights in the photoconversion of small fluorophores, they do not offer a general approach to obtain DCPFs. Moreover, it is important to keep on developing new mechanism of conversion to span DCPFs properties and their field of applications. In this regard, several features remain to be optimized. An ideal DCPF should 1) photoconvert with a high chemical yield, 2) display sufficient shift in emission wavelength and 3) possess a quantum yield of photoconversion, which depicts the rate of conversion, adapted to bioimaging (too fast conversion is unusable, a too slow one would lead to photodamage of living samples). In addition, both initial and converted forms should 1) be efficiently excited by usual microscopes laser lines, 2) be excitable in the visible range to avoid UV irradiation, 3) display a high brightness and 4) be efficiently targeted to the desired protein or organelle.

We recently introduced a new mechanism of photoconversion called Directed Photooxidation Induced Conversion (DPIC).^50^ This approach allowed to develop DCPFs based on the conjugation of a fluorophore to an Aromatic Singlet Oxygen Reactive Moieties (ASORM).^50^ Upon irradiation, the obtained red-shifted dye produces singlet oxygen (^1^O_2_) that reacts on the ASORM in a directed manner to disrupt the conjugation between the ASORM and the fluorophore leading to a blue-shift in emission. In this pioneer study,^50^ pyrrole was shown to be a suitable ASORM to obtain efficient photoconverters as it provided an high shift upon conversion (68 nm) with high efficiency (Quantum yield of photoconversion ϕ_Pc_ = 6.0 10^−2^ %) and decent yield of conversion that was further improved up to 63% by *N*-substitution of the pyrrole.^51^ Although our first coumarin-based photoconverters could be used to photoconvert lipid droplets^50^ and whole cells,^51^ their emission spectra were quite broad and their brightness was limited. On the contrary, BODIPYs is a family of bright fluorophores displaying narrow emission spectra.^52 53 54^ Their appealing properties led researchers to use them as a bright scaffold to develop photomodulable probes including photocages,^55 56^ photoactivatable,^8 12^ and photoconvertible probes.^42^ Consequently, BODIPYs found applications in SMLM. For instance, BAA Styryl-BODIPYs were used in combination with imaging buffers on fixed cells,^57^ whereas live SMLM was performed using red-shifted BODIPY,^58^ but also based on photoinduced disconnection of an oxazine heterocycle,^12^ transient formation of red-shifted emitting aggregates of green BODIPY,^59^ or through points accumulation for imaging in nanoscale topography (PAINT) in multidimensional super-resolution microscopy.^60^ Despite these examples, BODIPY dyes remain rarely used in SMLM applications compared to cyanines and rhodamines.

Herein we intended to apply the DPIC mechanism to BODIPY fluorophores to obtain bright dual-color photoconvertible fluorophores (DCPF) that can be further targeted to various subcellular domains in live cell imaging. In addition, we assessed the use of these photoconverters as photoswitchers to perform live SMLM.

## Results and discussion

To apply the DPIC mechanism to BODIPY, a pyrrole can be conjugated to the latter. Pyrrolyl-BODIPYs, displaying a pyrrole moiety in α position have been described as red-shifted BODIPY,^61 62^ but they have never been officially reported as photoconvertible fluorophores. However, in 2007, Freundt *et al*. published a letter to the editor to warn against the use of the widely used pyrrolyl-BODIPY: Lysotracker™ red (Figure 1A).^63^ The authors showed that Lysotracker™ red has tendencies to photoconvert toward a green-emitting form, spectrally close to Lysotracker™ green (Figure 1A), provoking false positive colocalization studies with green fluorescent proteins.^63 64^ In this letter, the authors suggested that this conversion might be the “result of a double bond shift or a reduction” at the level of the pyrrole ring leading to a disruption of conjugation thus provoking the change of emission colour. Since then, no rational explanation was provided.

**Figure 1.**
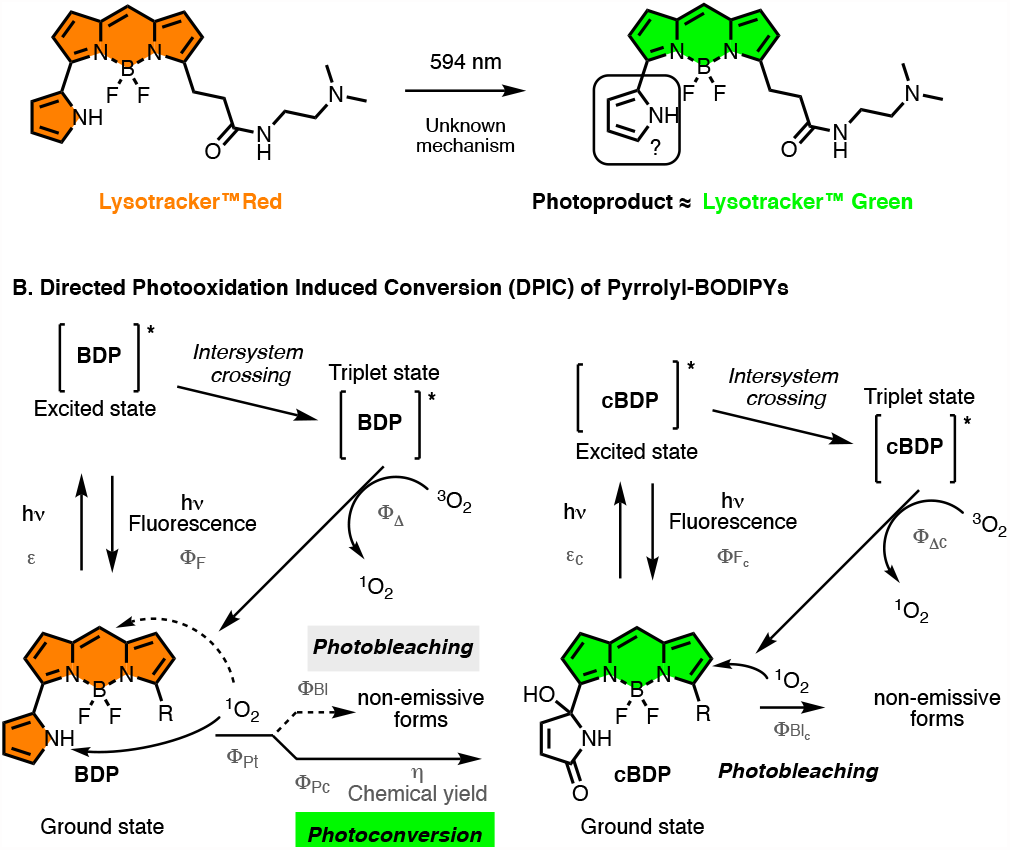
Structures of pyrrolyl-BODIPY Lysotracker™ Red and its unknown photoproduct spectrally close to Lysotracker™ Green as reported by Feundt *et al*. in 2007.^63^ Directed Photooxidation Induced Conversion (DPIC) mechanism applied to pyrrolyl-BODIPYs. cBDP stands for the converted forms of BDP. The constants in grey characterize the DPIC mechanism.

Interestingly, lysotracker red was used in live SMLM to image lysosomes,^58^ and a red-shifted pyrrolyl-BODIPY conjugated to phalloidin was also used in STORM without any specific buffer,^65^ suggesting that pyrrolyl-BODIPY are not only potential photoconverters but could also be efficient photoswitchers. We thus hypothesized that the mechanism that drives the photoconversion of pyrrolyl-BODIPYs could be our recently reported Directed Photooxidation Induced Conversion (DPIC) (Figure 1B). Upon excitation, pyrrolyl-BODIPYs (**BDP**) reach an excited state, which through intersystem crossing can reach an excited triplet state. The latter state can quench triplet oxygen (^3^O_2_) to generate reactive singlet oxygen (^1^O_2_). Once generated, ^1^O_2_ can react on the core BODIPY to generate non-emissive forms leading to photobleaching, or can react in a directed manner on the pyrrole moiety to generate converted blue-shifted emissive forms, **cBDP**, leading to photoconversion (Figure 1B). After conversion, **cBDP** follow the same pathway, except that its reaction with ^1^O_2_ will only lead to photobleaching. To prove our assumption, we used this pyrrolyl-BODIPY-scaffold to build targeted probes and to study their photoconversion and photoswitching properties.

### Synthesis of phosphonium BODIPYs

To show the unique photoconversion properties of pyrrolyl-BODIPYs through the versatility of the DPIC mechanism, three probes were synthesized (Figure 2A). Reactive *N*-hydroxysuccinimide ester BODIPYs namely: BDP-R6G, BDP-576 and BDP-650 were coupled to an amino-triphenyl phosphonium salt to 1) enhance their solubility in methanol and thus avoiding aggregation phenomenon during spectroscopic studies and 2) to target mitochondria in cellular photoconversion studies.^66^ **BDP-576-Mito** and **BDP-576-Mito** are both pyrrolyl-BODIPYs that possess a pyrrole moiety in α position of the BODIPY, which was shown to be an efficient aromatic singlet oxygen reactive moieties (ASORM) for the DPIC mechanism.^50^ **BDP-R6G-Mito** was synthesized as a control where the aromatic moiety in α position of the BODIPY, a phenyl group, is not known to be reactive towards singlet oxygen.

**Figure 2.**
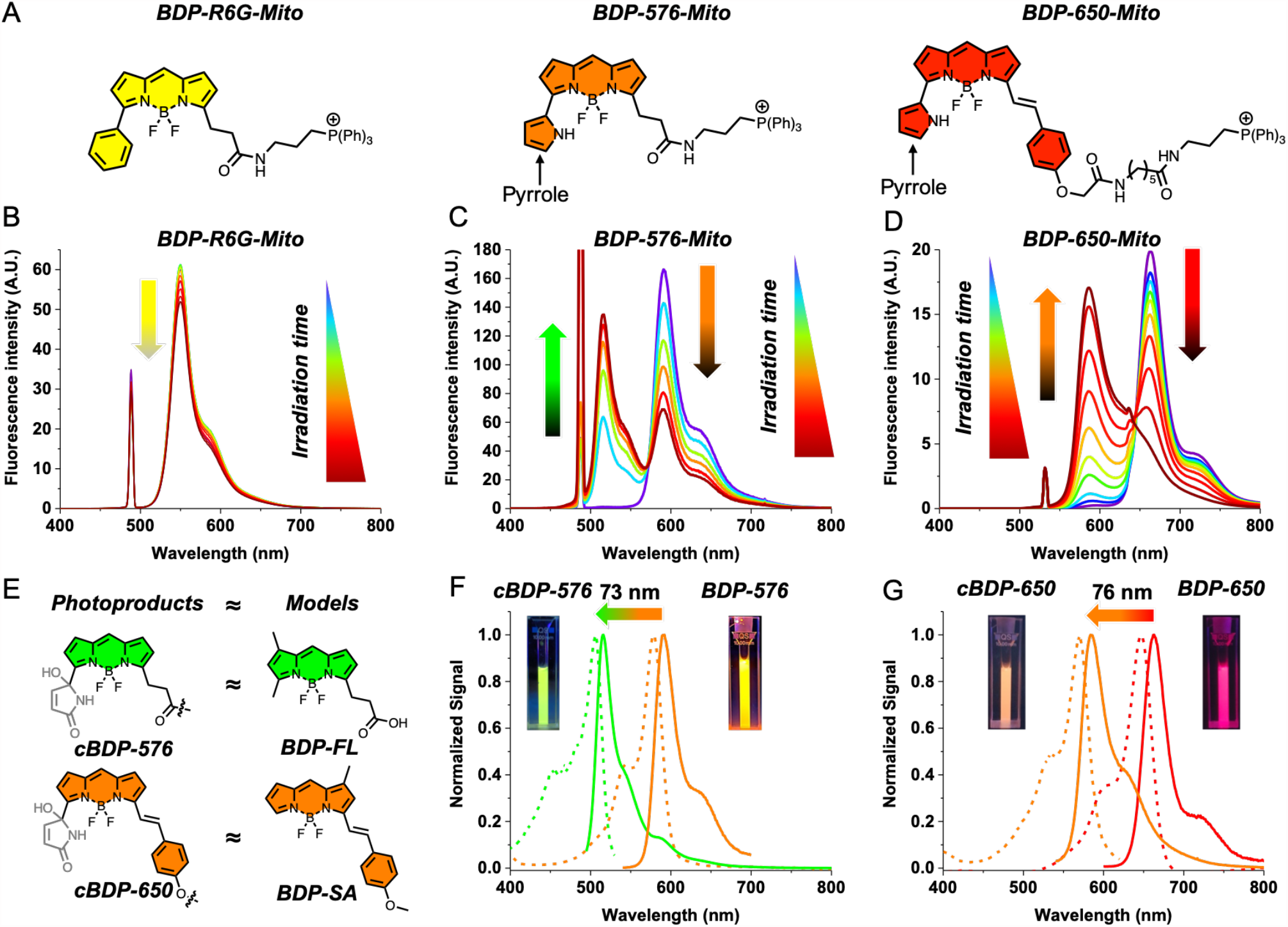
Photoirradiation of BDP-Mito showing the unique photoconversion features of pyrrolyl-BODIPYs. Structures of BDP-Mito: BDP-R6G-Mito and pyrrolyl-BODIPYs: BDP-576-Mito and BDP-650-Mito (A). Emission spectra of BDP-Mito (1 μM in MeOH) upon photoirradiation (B, C, D). BDP-R6G-Mito and BDP-576-Mito were irradiated at 532 nm for 5 h (61 mW.cm^-2^) and the emission spectra were recorded under 488 nm excitation. BDP-650-Mito was irradiated at 638 nm for 45 min (142 mW.cm^-2^) and the emission spectra were recorded under 532 nm excitation. For our study, BDP-FL and BDP-SA were taken as models of photoproducts for BDP-576-Mito and BDP-650-Mito respectively due to their close structure and properties (E). Emission (solid lines) and excitation (dashed lines) spectra of BDP-576-mito (F) and BDP-650-mito (G) and their photoproducts after conversion. Insets in F and G are pictures of the probes under UV light (365 nm) before and after conversion.

### Spectral properties

The photophysical properties of the three BODIPY-based mitochondrial probes were measured and confirmed that the addition of the phosphonium salt did not impair with their brightness as they displayed high quantum yields of fluorescence (Table 1). Although **BDP-R6G-Mito** is a yellow emitting dye (λ_abs max_ = 530 nm), **BDP-576-Mito** (λ_abs max_ = 576 nm) and **BDP-650-Mito** (λ_abs max_ = 650 nm) are respectively perfectly adapted to the red channel with typical excitation at 560 nm and the far-red channel with typical excitation at 640 nm laser line.

**Table 1.** Photophysical properties of BDP-Mito and their photoproducts cBDP-Mito (1 μM in MeOH).

| | Dye | $\lambda_{Abs}$<br>(nm) | $\epsilon$<br>($M^{-1}.cm^{-1}$) | $\lambda_{Em}$<br>(nm) | $\Phi_f$ (%)<sup>a</sup> | $\Phi_{Pt}$ (%)<sup>b</sup> | $\Phi_{Pc}$ (%)<sup>c</sup> | $\Phi_{Bl}$ (%)<sup>d</sup> | $\eta$ (%)<sup>e</sup> | $\Phi_{\Delta}$ (%)<sup>f</sup> |
| --- | --- | --- | --- | --- | --- | --- | --- | --- | --- | --- |
| BDP-Mito | BDP-R6G | 530 | 70,000 | 548 | 96 | $1.1 \pm 0.2 \cdot 10^{-4}$ | N/A | $1.1 \pm 0.2 \cdot 10^{-4}$ | N/A | 0.8 |
| | BDP-576 | 575 | 83,000 | 588 | 70 | $3.1 \pm 0.5 \cdot 10^{-4}$ | $2.4 \pm 0.4 \cdot 10^{-5}$ | $2.9 \pm 0.5 \cdot 10^{-4}$ | $7.7 \pm 0.4$ | 1.4 |
| | BDP-650 | 646 | 102,000 | 660 | 52 | $3.6 \pm 0.1 \cdot 10^{-4}$ | $1.03 \pm 0.04 \cdot 10^{-4}$ | $2.5 \pm 0.1 \cdot 10^{-4}$ | $28.9 \pm 1.3$ | 1.3 |
| cBDP-Mito | cBDP-576 | 506<sup>g</sup> | N/A | 515 | N/A | $2.8 \pm 0.8 \cdot 10^{-5}$ | N/A | $2.8 \pm 0.8 \cdot 10^{-5}$ | N/A | N/A |
|  | cBDP-650 | 570<sup>g</sup> | N/A | 584 | N/A | N/A<sup>h</sup> | N/A<sup>h</sup> | N/A<sup>h</sup> | N/A | N/A |

<sup>a</sup> Fluorescence quantum yield<sup>b</sup> Quantum yield of phototransformation<sup>c</sup> Quantum yield of photoconversion<sup>d</sup> Quantum yield of photobleaching<sup>e</sup> Chemical yield of photoconversion<sup>f</sup> Quantum yield of generation of $^1O_2$ <sup>g</sup> Based on the excitation spectrum<sup>h</sup> Stable in our conditions

### Photoconversion properties

To assess their photoconversion properties, the three probes were continuously irradiated by a continuous wave laser and their emission spectra were recorded live over the time (Figure 2B-D). Unlike **BDP-R6G-Mito** that displayed a low photoreactivity leading to slow photobleaching (Figure 2B), pyrrolyl-BODIPYs **BDP-576-Mito** and **BDP-650-Mito** rapidly photoconverted towards new emissive photoproducts with an hypsochromic shift in emission (Figure 2C and D, Table 1). As reported by Freundt *et al*.,^63^ the converted form of **BDP-576-Mito**, namely **cBDP-576-Mito** possessed similar photophysical properties than a typical green emitting BODIPY like **BDP-FL** (Figure 2E and S1) displaying sharp spectra and low Stokes shift with λ_ex_ = 506 nm and λ_em_= 515 nm (Figure 2F, table 1). Similarly, **cBDP-650-Mito**, the converted form of **BDP-650-Mito**, displayed sharp excitation and emission spectra with λ_ex_ = 570 nm and λ_em_ = 584 nm (Figure 2, table 1), spectrally close to mono styryl BODIPYs,^67 68^ and **BDP-SA** (Figure 2E and S2), a Styryl-Anisol BODIPY that we synthesized as a model of photoproduct of **cBDP-650**. Hence, these results suggested that upon irradiation a disruption of conjugation between the BODIPY and the pyrrole in α position occurred.

The features of these photoconverters are advantageous in microscopy as both the probes and their photoproducts are perfectly adapted to typical green, red and far-red channels, with important hypsochromic shifts of 73 nm and 76 nm after conversion thus limiting crosstalk of channels in imaging (Figure 2F-G).

Encouraged by these results, the properties of conversion were deeply studied by the determination of key photophysical constants (Figure 1, grey constants) that are reported in table 1. As depicted in figure 1, upon irradiation pyrrolyl-BDP can undergo a phototransformation (characterized by its quantum yield of phototransformation: ϕ_Pt_), that can be either photobleaching (characterized by its quantum yield of photobleaching: ϕ_Bl_), photoconversion (characterized by its quantum yield of photoconversion: ϕ_Pc_) or both, in such a way that ϕ_Pt_ = ϕ_Bl_ + ϕ_Pc_. The fit of fluorescence decay of the initial form upon irradiation with exponential model (Figure S2) enables to determine the quantum yield of phototransformation (ϕ_Pt_). The chemical yields of conversion (η) were determined using models of emissive photoproduct **BDP-FL** and **BDP-SA** (for synthesis and spectra, see SI, figure S1), where the pyrrole has been removed (Figure 2E) and was used to determine the quantum yield of photoconversion (ϕ_Pc_) and thus the quantum yield of photobleaching (see materials and methods section).

As expected, **BDP-R6G-Mito** possesses a lower quantum yield of phototransformation (ϕ_Pt_) compared to the reactive pyrrolyl-BODIPYs. Interestingly, whereas **BDP-576-Mito** and **BDP-650-Mito** have similar ϕ_Pt_ (∼3 10^−4^ %), the latter displays a 2.3-fold higher quantum yield of photoconversion (ϕ_Pc_), depicting a faster conversion rate. Consequently, and based on our calculations (see experimental section), **BDP-650-Mito** converts with a better yield (η = 28.9 ± 1.3%) compared to **BDP-576-Mito (**η = 7.7 ± 0.4%). Overall it appeared that **BDP-650-Mito** displayed better photoconversion properties compared to **BDP-576-Mito**.

Although pyrrolyl-BODIPYs possess much lower quantum yields of phototransformation (∼3×10^−4^ %) compared to photoconvertible proteins ranging from 10^−1^ to 10^−2^ % like mKikGR,^69^ Kaede,^31^ and others,^70 71^ they might actually be more suitable for photoconversion in cellular imaging. Indeed, as far as our experience in the field, photomodulatable probes with high quantum yield of phototransformation are prone to undesired photoconversion and are thus difficult to handle in bioimaging.

As shown in Figure 1, the efficiency of a converter based on the DPIC mechanism, depends on its ability to generate singlet oxygen as well as its reactivity towards it. Consequently, the quantum yields of generation of ^1^O_2_ (ϕ_Δ_) of the BDP-Mito have been determined. Once again, the results showed that **BDP-576-Mito** and **BDP-650-Mito** have similar ϕ_Δ_ (Table 1), thus suggesting that the better photoconversion properties of **BDP-650-Mito** arose from its higher reactivity towards singlet oxygen and not its ability to generate some. Importantly, **BDP-576** and **BDP-650** with ϕ_Δ_ of 1.4% and 1.3% respectively did not significantly produce more ^1^O_2_ than non-convertible **BDP-R6G-Mito**, suggesting that the pyrrole moiety does not enhance ϕ_Δ_. This result is important, as it shows that whereas the photoconverters based on DPIC require the production of ^1^O_2_ to convert, they do not produce more singlet oxygen than other fluorophores and that they are just more reactive towards it. Hence, pyrrolyl-BODIPYs are not necessarily more phototoxic than other dyes. Indeed these ϕ_Δ_ of ≈ 1% (in methanol) is even lower than other common fluorophores like fluorescein (3% in methanol,^72^ 13% in EtOH,^73^ according to the literature).

Importantly, the converted probes must be sufficiently photostable to avoid photobleaching upon conversion of their non-converted cognates as well as during their imaging after conversion. Consequently, the photostability of the converted forms after conversion was assessed. In our conditions (irradiance at 532 nm, 66.6 mW.cm^-1^) **cBDP-650** was too stable to determine its ϕ_Bl_, which depicted a high photostability. The green-emitting **cBDP-576** with a ϕ_Bl_ = 2.8 ± 0.8 10^−5^ was found to be much more photostable than the common BODIPY-493 (ϕ_Bl_ = 2.1 ± 0.5 10^−3^, figure S3) and as stable as non-convertible **BDP-R6G-Mito** (Table 1) as well as converted form of photoconvertible proteins.^69^

Overall, these results showed that pyrrolyl-BODIPYs possessed efficient photoconversion properties with relatively good yields of conversion, balanced photoconversion rates along with photostable emissive photoproducts. Combined to adapted spectral properties and potentially high brightness these photoconverters appeared adapted to bioimaging applications.

### Mechanism of conversion

Although pyrrolyl-BODIPY displayed efficient photoconversion properties, several experiments were performed to show that the DPIC mechanism was involved. First of all, HPLC-mass analysis after irradiation by visible light showed that the main photoproducts were the result of oxidation [M+O_2_] (Figure S4). To prove the implication of photooxidation in the conversion process several experiments were conducted. First, the pyrrolyl-BODIPYs were exposed to several Reactive Oxygen Species (ROS). Singlet oxygen (^1^O_2_), hydroxyl radical (OH•) were both found to trigger the conversion at the ground state of the dyes (Figure S5-S6). Then, irradiation of pyrrolyl-BODIPYs in the presence of DPBF (a singlet oxygen probe) and HPF (a hydroxyl radical probe) proved that both **BDP-576** and **BDP-650** generate ^1^O_2_ and OH• upon irradiation (Figure S5-S6). In addition, conversion was performed in deuterated methanol where the lifetime of singlet oxygen is increased,^74^ and resulted in an acceleration of the transformation with increased quantum yields of photo-transformation (Figure S7). These experiments showed that upon irradiation pyrrolyl-BODIPYs generate ROS (^1^O_2_ and OH•) that are directly responsive for their conversion, which is in line with the DPIC mechanism.^50^

Overall, these results reaffirm that DPIC was responsible for the observed photoconversion of pyrrolyl-BODIPYs as they are perfectly in line with our previous study on coumarin-pyrrole based photoconverters.^50^

### Cytotoxicity and phototoxicity

Despite their low quantum yield of generation of singlet oxygen (ϕ_Δ_, table 1), the cytotoxicity and the phototoxicity of both **BDP-576-Mito** and **BDP-650-Mito** were assessed by MTT viability test prior to microscopy studies. The results indicated that they were not cytotoxic at the concentration used in imaging (200 nM) and were not phototoxic upon the irradiation time required for their photoconversion *in cellulo* (Figure S8). Interestingly, in the same conditions, widely used MitoTracker™ deep red, showed slightly higher (though statistically non-significant) phototoxicity, thus strengthening the point that these photoconverters do not exhibit enhanced phototoxicity compared to other fluorescent probes that are commonly used in live cell imaging.

### Photoconversion in cells

To extend the applications of the pyrrolyl-BODIPY as photoconverters in cellular imaging, BDP-576 and BDP-650 were targeted to various sub-cellular structures and organelles using selective targeting moieties (Figure 3).^75 76^ Consequently, in addition to **BDP-576-Mito** and **BDP-650-Mito** that bear a triphenyl phosphonium moiety to targets the inner membrane of mitochondria, BDP-576 and BDP-650 were also converted into plasma membrane (PM) probes: **BDP-576-PM** and **BDP-650-PM**, using our targeting moiety, whose efficiency has already been proven (Figure 3).^77 78 79 80^ Owning to its better performance compared to BDP-576, *i*.*e*. higher ϕ_Pc_ and conversion yield (η), BDP-650 was chosen to target other subcellular components. Phalloidin was thus coupled to BDP-650 to obtain **BDP-650-Actin** that targets the actin filaments, and a PEG_12_-Halo tag was used to obtain **BDP-650-Halo** to selectively perform *in cellulo* staining of genetically encoded Halo-tagged proteins (Figure 3).^81^

**Figure 3.**
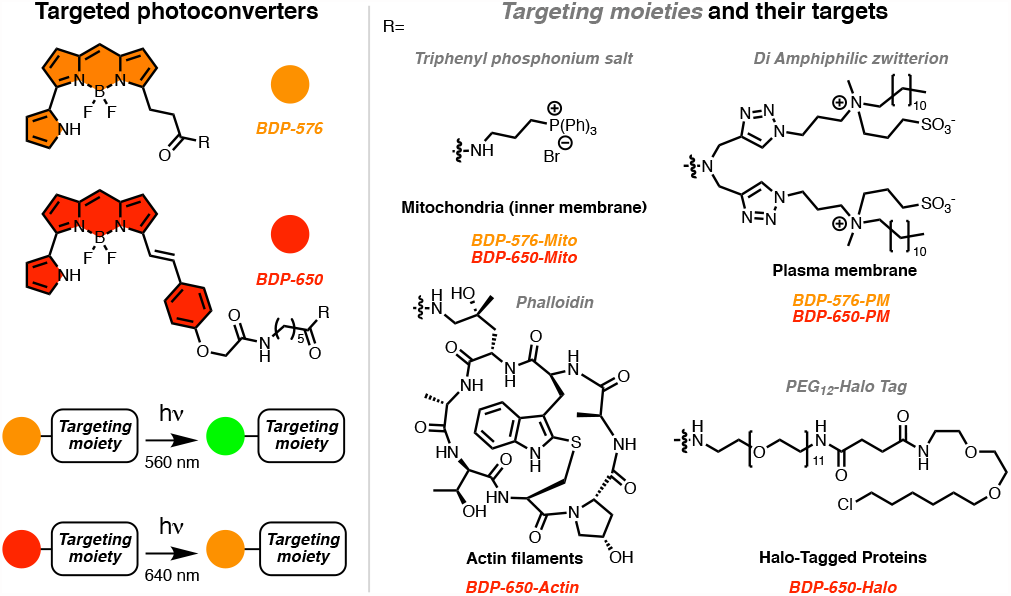
Targeted photoconverters based in Pyrrolyl-BODIPYs, obtained by coupling BDP-576 and BDP-650 with various targeting moieties.

After checking the kinetic of binding of **BDP-576-PM** and **BDP-650-PM** using large unilamellar vesicles as PM models (Figure S9) the selective targeting of the plasma membrane probes and the mitochondrial probes were verified on HeLa cells using MemBright and Mitotrackers probes respectively as reference co-staining probes (Figure S10).

First, the photoconversion of **BDP-576** based-probes in live cells was undertaken. Using a laser scanning confocal microscope, cells labelled with **BDP-576-Mito** could be imaged in the initial form’s channel for several frames without triggering undesired photoconversion of mitochondria. Then the photoconversion of a single cell was readily performed by applying a zoom on a region of interest followed by several scans at higher laser power and by monitoring the signal’s decrease in the initial form’s channel and the signal’s increase in the converted form channel (Figure S11). The photoconversion of **BDP-576-Mito** performed within ≈ 30 s with no sign of phototoxicity. Using **BDP-576-PM**, the plasma membranes of HeLa cells were sequentially converted without triggering the conversion of adjacent cells (Figure S12). In addition, both **BDP-576-Mito** and **BDP-576-PM** displayed an important fluorescence enhancement in the “converted channel” of 6-8-fold (Figure S11, S12, and S17), which showed the efficiency of photoconversion in both mitochondria and plasma membrane.

In a second time, live cell photoconversion of targeted **BDP-650** probes were evaluated. Owning to the high brightness of **BDP-650, BDP-650-Mito** (Figure 4A and B, S13) and **BDP-650-PM** (Figure 4C and D, S14) provided an intense and selective staining and could be converted in tens of seconds with fluorescence enhancement in the “converted” channel ranging from 3- to 7-fold, along with stable converted forms (Figure 4A-D). Although **BDP-650-Actin** required more irradiation time (80 s) to reach the plateau of fluorescence intensity of the converted form (Figure 4 E-F, S15), it provided a clear photoconversion of actin fibbers. This difference might be assigned to the fact that the cells were fixed compared to live cell experiments with **BDP-650-Mito** and **BDP-650-PM** or by the non-lipid environment actin compared to membrane-based mitochondria and PM.

**Figure 4.**
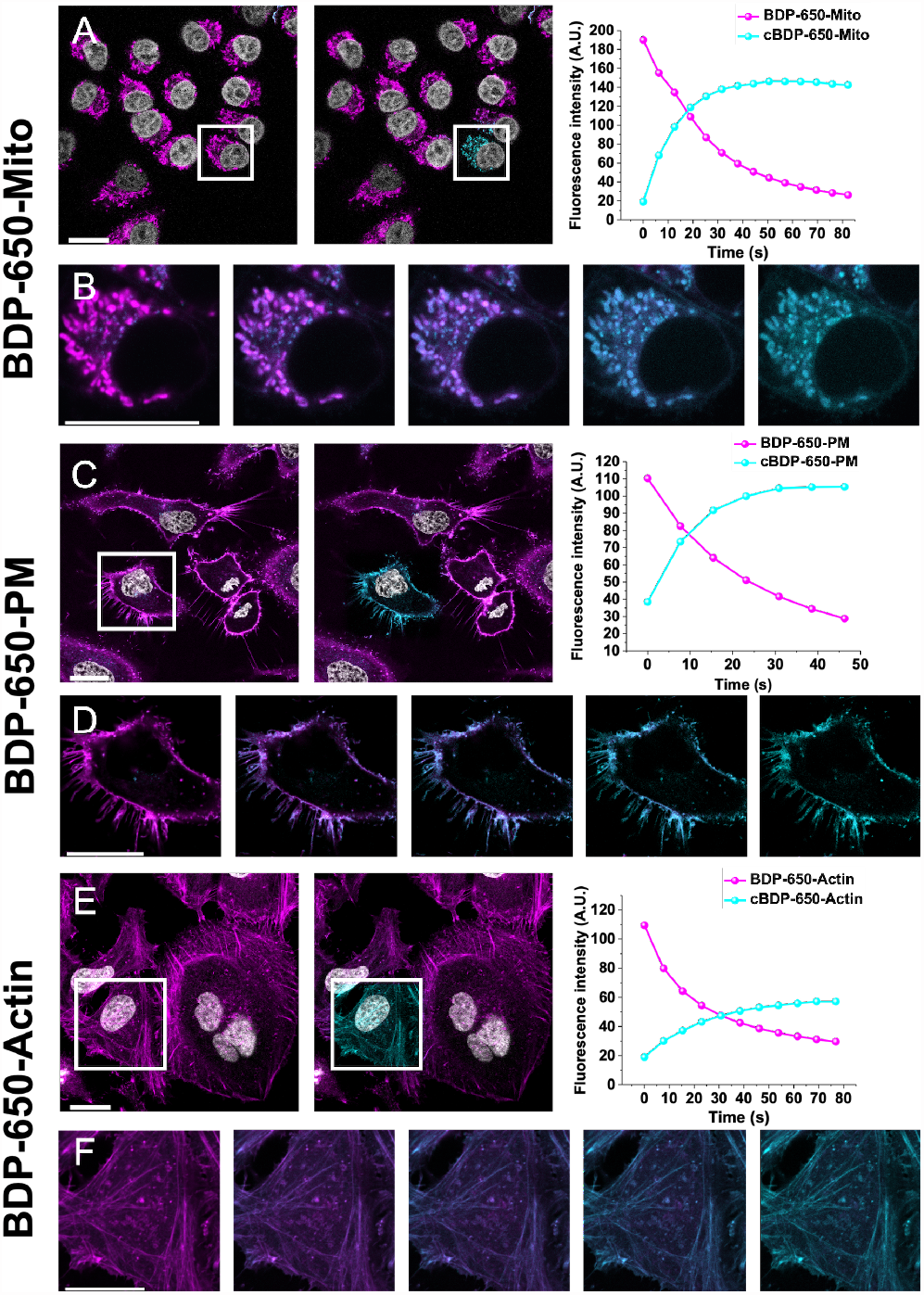
Photoconversion of various cellular structures using BDP-650 targeted probes. (A, C) Laser scanning confocal microscope images of live KB (A) and HeLa (C) cells stained with BDP-650 targeted probes (200 nM) before (left) and after (middle) photoconversion, and fluorescence signal (mean intensity of the region of interest, white frame) in the BDP-650 channel (Magenta) and the cBDP-650 one (cyan) over time (right). (E) Laser scanning confocal microscope images of fixed (4% PFA) HeLa cells stained with BDP-650-Actin (200 nM). The nucleus (grey) was stained with Hoechst 33258 (5 μg.mL^-1^). (B, D, F) Merged channels of the zoomed regions of interest upon photoconversion. Scale bar is 20 μm.

In a next series of experiments, **BDP-650-Halo** was used to stain and photoconvert different organelles by targeting genetically-encoded Halo-tagged proteins (Figure 5, S16A-E). After a washing step, **BDP-650-Halo** provided an intense and selective staining of the organelle-selective Halo-tagged proteins. Upon irradiation of regions of interest, actin fibbers, nuclei, mitochondria, Golgi apparatus and nucleoli have been successfully photoconverted in live cells (Figure 5, S16A-E).

**Figure 5.**
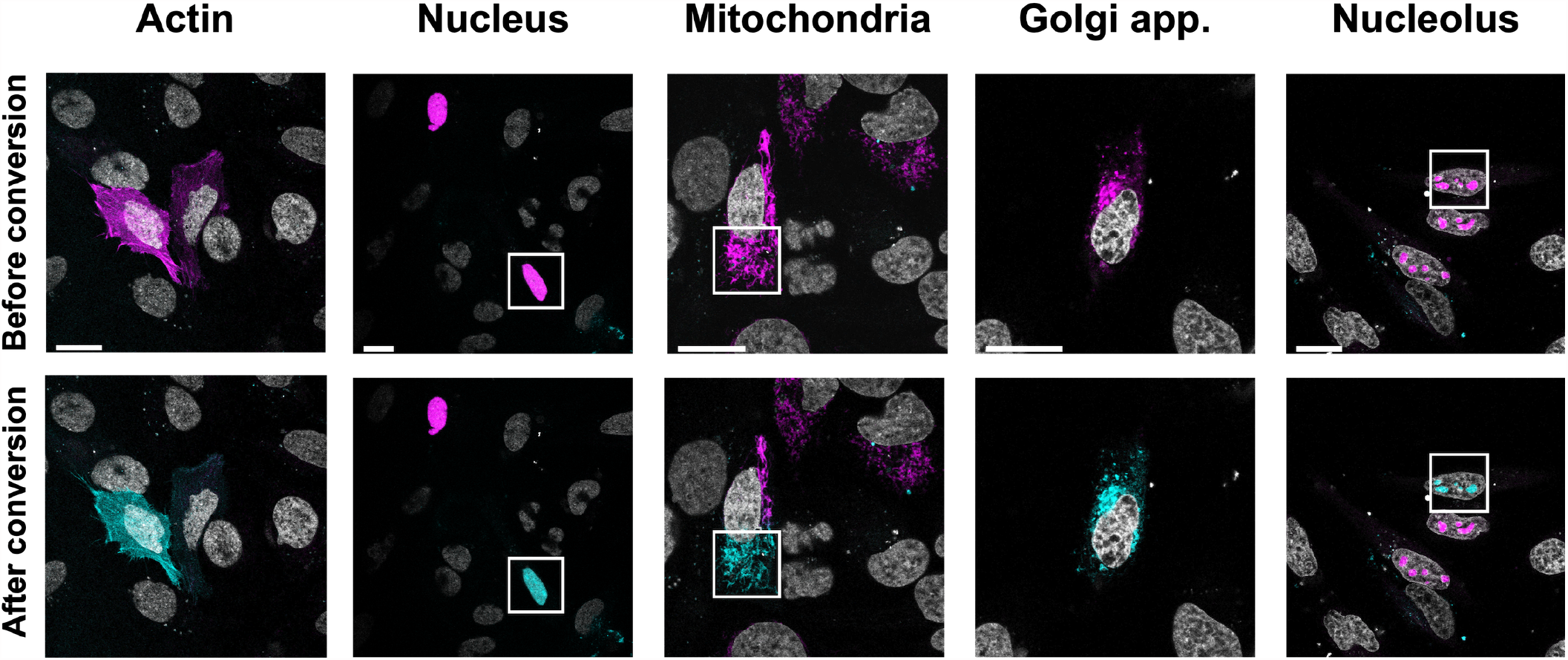
Photoconversion of genetically-encoded Halo-tagged proteins in live cells using BDP-650-Halo. Laser scanning confocal microscope images of live HeLa cells transfected with various plasmids encoding for organelle specifically located-proteins followed by staining with BDP-650-Halo (50 nM). The images correspond to the merge of channel BDP-650 (magenta) and cBDP-650 (cyan) before conversion (top panel) and after conversion (bottom panel). The white frames correspond to the region of interest that has been irradiated. The nucleus (grey) was stained with Hoechst 33258 (5 μg.mL^-1^). Scale bar is 20 μm.

To quantify the efficiency of the photoconversion during microscopy imaging, the signal to noise ratios before and after photoconversion of the different probes have been measured and the results along with the signal to noise ratio enhancement of the converted form after conversion are reported in figure S17. Surprisingly, the plasma membrane and mitochondrial probes showed quite similar signal to noise ratio in both channels (between 20-30) and high signal to noise ratio enhancement of the converted form after conversion (up to 11-fold). Interestingly, the conversion in mitochondria was more efficient using a targeting moiety than through targeting a Halo-tagged protein (S/N enhancement of the converted form: 8 and 9 for **BDP-576-Mito** and **BDP-650-Mito**, 3 for **BDP-650-Halo** in Mitochondria). Conversely, the conversion of actin fibbers using either a drug as a ligand (phalloidin) or by targeting a Halo-tagged protein, gave lower S/N ratios for both the initial and converted form. The best S/N have been obtained with **BDP-650-Halo** especially when targeted to nuclear protein and proteins of the nucleolus (Figure S17). However, the S/N enhancement of the converted form in this condition ranged between 3 to 5-fold. These data suggested that the image quality upon conversion is highly dependent on the localization of the converter. Furthermore, it is noteworthy that, conversely to “chemically” targeted probes, the signal of **BDP-650-Halo** strongly depends on the quantity of expressed proteins by the cells which affects the S/N.

Finally, to prove the usefulness of our converters in tracking applications, HeLa cells were incubated with **BDP-650-PM**. After endocytosis of the PM probe, fluorescently labelled intracellular vesicles (IVs) were obtained (Figure 6A). Several IVs were then successfully converted in a region of interest within 3 s (fast, due to the small area of conversion), to obtain two populations of vesicles that could be clearly discriminated between the non-converted channel (Figure 6B) and the converted one (Figure 6C). Owning to the adapted photoconversion quantum yield of photoconversion of **BDP-650** (ϕ_Pc_ = 1.03 10^−4^ %) and the high photostability of its converted form, **cBDP-650**, both non-converted and converted individual IVs could have been tracked in an unambiguous manner over 21 min (300 frames) (Figure 6D-I, Movie 1).

**Figure 6.**
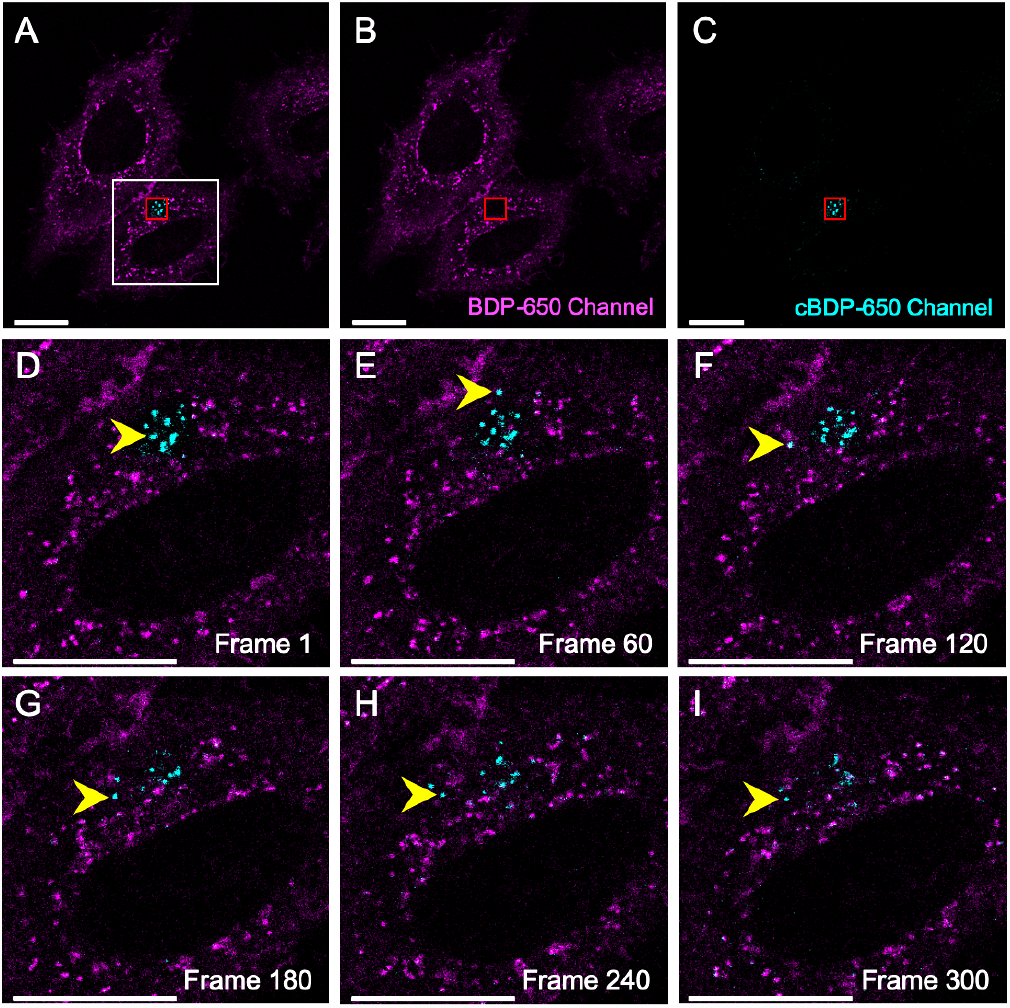
Tracking of intracellular vesicles labelled with BDP-650-PM. (A) HeLa cells were stained with BDP-650-PM and were incubated 3 h to promote endocytosis and to obtain labelled intracellular vesicles. Several vesicles were converted (red box) to obtain two populations of vesicles that could be clearly discriminated between the non-converted channel (B) and the converted one (C). (D-I) Tracking of individual IV (yellow arrow) in a region of interest (white frame in A) over 21 min (300 frames). Scale bar is 20 μm.

Overall, pyrrolyl-BODIPY photoconverters have been targeted to various organelles and subcellular compartments and provided an efficient and fast photoconversion using a laser scanning confocal microscope and proved their efficacy in tracking subcellular compartments over large spatiotemporal scales.

### Photoswitching properties

As discussed in the introduction, photomodulable fluorescent probes generally find application in super resolution imaging. Indeed, due to their ability to modify their emission properties upon irradiation, they can provide blinks enabling Single Molecule Localization Microscopy (SMLM). To evaluate the photoswitching properties of our probes, the pyrrolyl-BODIPYs were coupled to a biotin-ended PEG for immobilization on glass surface (Figures S18). Single molecule analysis was performed in PBS (without any oxygen scavenger) and the switching properties of **BDP-576** and **BDP-650** are reported in Table 2. Since the photoproduct of our converters are well spectrally separated from their initial form and perfectly fits to conventional fluorescent imaging channels (488, 560 and 640 nm excitation laser, typically green, red and far-red channels), their single molecular switching properties were studied in both the initial channel and the converted one. In addition, we also compared the blinking properties in the converted channels before and after irradiation of the initial form (Table 2). In our conditions (Medium power laser:^82^ 561 nm at 0.33 kW.cm^-2^ irradiance), the pyrrolyl-BODIPYs displayed a high number of photons per blink in both initial and converted forms (up to 9,234 for **cBDP-650**) and thus high localization precisions ranging from to 14.8 nm. Due to the absence of additional singlet oxygen scavengers, the pyrrolyl BODIPYs displayed rather low photoswitching times (typically 7-8 s, expect for **cBDP-576**), which are compensated by relatively high number of switching cycle (up to 10.8 for **BDP-650**). Consequently, the converters showed quite high duty cycles, especially **BDP-650** (0.13). Compared to dyes with similar conditions of irradiance and photophysical properties: BODIPY FL as green dye, Cy3B as red dye, AF-647 as far-red dye, and in presence of oxygen scavengers,^82^ pyrrolyl-BODIPYs: respectively **cBDP-576** (green), **BDP-576, cBDP-650** (red) and **BDP-650** (far-red) in physiological conditions displayed higher localization precision, and higher to similar switching cycles. Interestingly, the photoswitching properties of the converted forms appears excellent either upon direct excitation or after their conversion. Consequently, the converted channel constitutes a reservoir of supplementary blinks after the use of the initial channel that could be advantageously used in live SMLM imaging.

**Table 2.** Single molecule analysis and switching properties of BDP-576 and BDP-650. The biotinylated dyes were immobilized on neutravidin functionalized glass and imaging was performed in PBS. Excitation wavelength 488 nm for cBDP-576, 561 nm for BDP-576 and cBDP-650, and 642 nm for BDP-650. The converters were studied in the initial channels, directly in their converted channels (Direct), and in their converted channel after irradiation in the initial channel (After conv.).

| Dye | Photons per blink | Switching Cycles | Total photons | Localization Precision (nm) | Photo-switching Time (s) | Duty Cycle | Molecules analysed | Molecules removed (%) |
| --- | --- | --- | --- | --- | --- | --- | --- | --- |
| **BDP-576** | 2,617 ± 1,316 | 7.9 ± 1.5 | 13,471 ± 9,030 | 11.2 ± 0.3 | 7.5 ± 0.9 | 0.055 ± 0.015 | 246 ± 8 | 57 ± 3 |
| **cBDP-576 (Direct)** | 1,120 ± 916 | 6.4 ± 0.9 | 4,950 ± 3,663 | 14.8 ± 1.9 | 22.8 ± 18.9 | 0.042 ± 0.005 | 125 ± 23 | 47 ± 3 |
| **cBDP-576 (After conv.)** | 771 ± 311 | 6.5 ± 0.5 | 3,829 ± 2,360 | 13.2 ± 0.4 | 39.2 ± 0.3 | 0.037 ± 0.002 | 149 ± 4 | 48 ± 2 |
| **BDP-650** | 5,721 ± 526 | 10.8 ± 1.0 | 35,465 ± 6,163 | 14.2 ± 0.15 | 7.7 ± 0.2 | 0.13 ± 0.01 | 1,362 ± 52 | 34 ± 1 |
| **cBDP-650 (Direct)** | 9,234 ± 1,539 | 6.8 ± 0.1 | 42,641 ± 6,871 | 13.0 ± 0.13 | 7.2 ± 0.2 | 0.071 ± 0.001 | 1,165 ± 9 | 51 ± 2 |
| **cBDP-650 (After conv.)** | 6,560 ± 143 | 6.1 ± 0.6 | 27,436 ± 1,316 | 11.9 ± 0.4 | 8.0 ± 1.1 | 0.067 ± 0.002 | 922 ± 58 | 53 ± 2 |

To sum up, in physiological conditions, pyrrolyl-BODIPYs blink fast and bright for a short time, along with the possibility of using the converted channel as a reservoir of blinks. All these features are appreciable for live SMLM where an efficient switching and a high number of blinking events are required within short acquisition time.

However, these results interrogated us regarding the mechanism of blinking. Although photoconvertible probes based on an irreversible process have been shown to be efficient in SMLM,^42^ one can thus wonder how multiple blinking event could be obtained from pyrrolyl-BODIPYs undergoing the DPIC mechanism. To decipher the involved mechanism, single molecule studies were carried out on **BDP-650** upon 560 nm excitation and where the fluorescence signal has been separated in two channels allowing to simultaneously and independently monitor both the initial form and the converted one. Single molecule traces indicated no obvious correlations between the two signals (Figure S19), suggesting that within our acquisition time (30 ms), complex and fast equilibria occurred between dark states, the initial form and the converted form. Although a dedicated study would be required to decipher the involved mechanism, we hypothesized that a reversible cycloaddition of singlet oxygen on BDP and cBDP through the formation of transient endoperoxide intermediates might be involved.^83 84 85^

Regardless the involved mechanism, pyrrolyl BODIPYs displayed promizing switching properties that could be advantageously used in live SMLM imaging.

### Live SMLM imaging

SMLM can be used to unravel the localization of single molecule either on live or fixed cells. Within SMLM family, one can distinguish PALM that uses photoswitchable or convertible fluorescent proteins (FPs), STORM that uses antibodies or probes coupled to fluorescent photoswitchable organic dyes and DNA PAINT that uses fluorescent oligonucleotides that can bind /unbind their target antibodies within the field of view allowed with TIRF microscopy.

To detect single molecules, STORM and PALM imaging are relying on fluorescent probes able to photoswitch between an excited fluorescent state (“ON” phase) and a dark state (“OFF”). Some fluorophores used in conventional live or confocal imaging can photoswitch but not necessarily spontaneously. That’s the reason why, fluorophores used in STORM microscopy are incubated in “blinking buffers” that favour this alternation between “ON” and “OFF” states. Those blinking buffers contain both oxygen scavengers and reducers (MEA or β-Mercaptoethanol) that are cytotoxic and thus unfriendly for live cell imaging. For these reasons, in the last years, most STORM experiments were done on fixed cells, while live SMLM was done using photoactivable fusion proteins (PALM imaging).

The dark state of fluorophores can be reduced in a “triplet” state, which is a non-emitting form. UV light pulse can bring back fluorophores at the ground state (back-pumping) allowing then a second run of excitation. However, UV light is known to produce singlet oxygen and reactive oxygen species (ROS) which induce cell damage and cytotoxicity.^86^ Consequently, successfully combining all those parameters is a challenging task to achieve live STORM imaging in a physiological manner.

During this last decade, new suitable tools have emerged for super-resolution imaging in living organisms.^87^ A pioneering study in 2010 could elegantly achieve live SMLM of histone at 20°C, using Glutathione as antioxidant.^88^ Later Zhuang *et al*. could image Transferrin labelled with a photoswitchable dye pair, AF405 and AF647 at 34°C for up to 1 min.^89^ They used 0.5% beta-mercaptoethanol to support photoswitching and an oxygen scavenger system (glucose oxidase / catalase) to reduce photobleaching. To achieve fast imaging speeds (0.5 to 6.0 s imaging/frame), they used a 657 nm laser up to 15 kW.cm^−2^ and a 405 nm laser that was two to four orders of magnitude less intense. Thanks to these parameters the authors could follow the internalization of transferrin in 54 s. More recent discoveries have shown that STORM imaging could be done with a resolution around 20 nm on short time scale experiments using UV light.^90 91^

Reducing compound can be deleterious for living cell and may induced irreversible cellular dysfunction. For example, glycans can be reduced and provokes modifications of the transmembrane proteins dynamic and electrical properties of ionic channels. In contrast to beta-mercaptoethanol, glutathione is an endogenous component of cellular metabolism and is thus more closely related to cell physiology. However, it is known to have binding sites in central nervous system and can act as neuromediator.^92^ Up to now, despite all the remarkable efforts that have been made to combine live imaging and STORM, we are still lacking bright organic dyes adapted for STORM imaging on living organisms and capable of self-blinking properties in physiological conditions. We thus assessed the ability of BDP-650 probes to switch in physiological buffer at 37°C and its compatibility with fragile neuronal cells.

### Optimal conditions for live 3D-STORM imaging in physiological buffer

Since higher wavelength are known to be less detrimental to live cells,^86^ we determined if blinking properties of **BDP-650** measured *in vitro* was conserved during *in vivo* live cell imaging. We first evaluated the efficiency of **BDP-650** in live cell imaging condition using the mitochondrial probe **BDP-650-Mito** (Figure 7A). To do so, we realized 3D-STORM imaging on fragile neuronal cells in physiological conditions: in Krebs-Ringer solution at 37°C without any reducing agents. Our first concern was to determinate if **BDP-650** could work at a temporal scale that would fit with cellular dynamic processes. Using the native form (λ_ex_= 640 nm) our results showed that **BDP-650** can be used with live cell STORM imaging during at least 2 minutes (Figure 7B-D). Over this time window we detected more than 90 000 particles with a mean radial precision of 44 nm (Figure 7E). As it can be seen on Figure 7B and C, we were able to reconstruct with high fidelity every detail, even the finest, within the 3D mitochondrial network. Entertainingly we noticed that some mitochondria were still moving during the acquisition since the localization of some of the reconstructed mitochondria (Z rainbow scale) were shifted when considering the first reference widefield image (red in Figure 7B).

**Figure 7.**
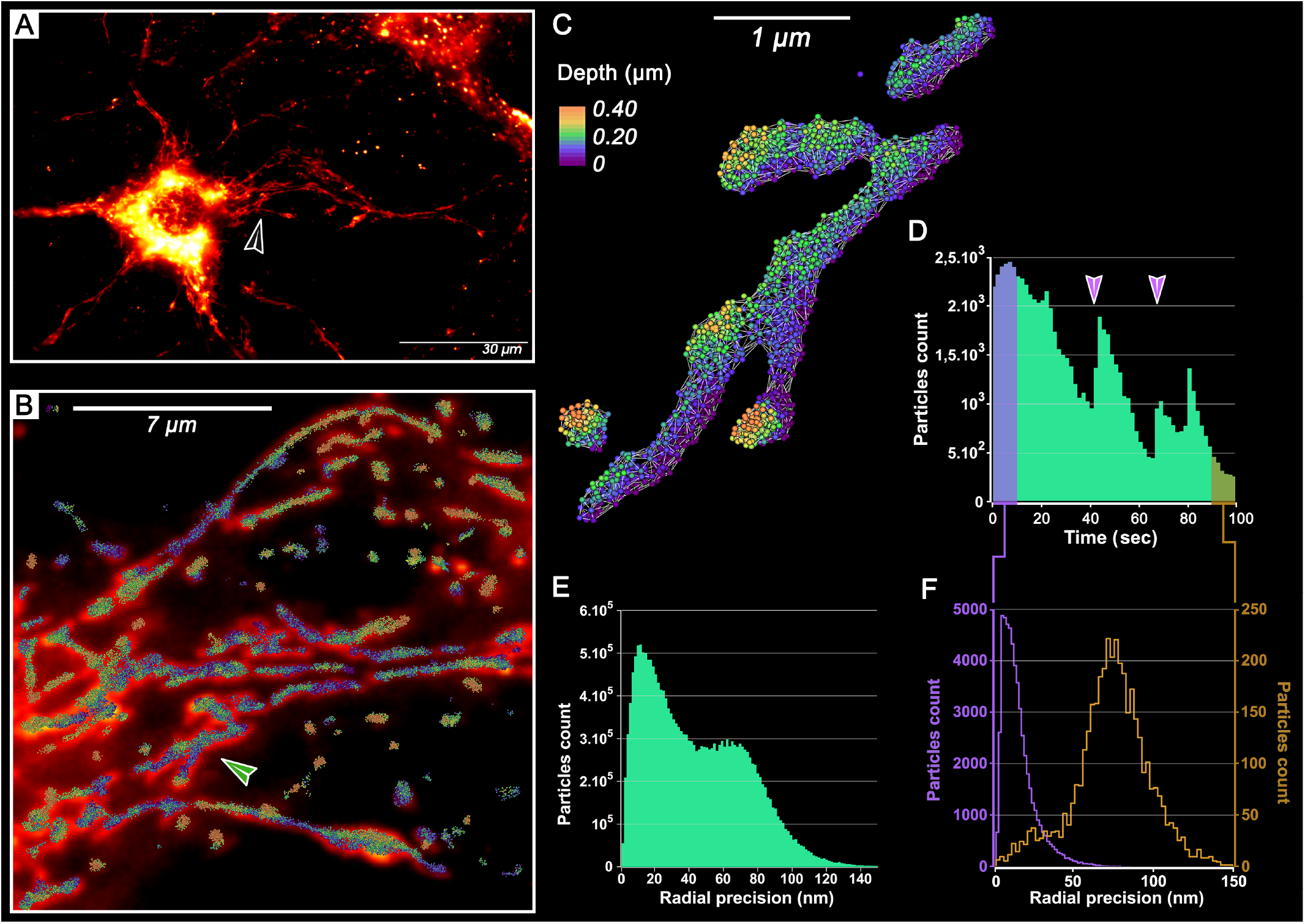
3D-STORM imaging of mitochondria in living neuron using BDP-650-Mito at 37°C in physiological buffer. (A) Widefield fluorescence image of mitochondria in living 21 days in vitro (DIV) neuron, labelled with BDP-650-Mito. Color-code represents fluorescence intensity. Black arrow indicates the region acquired in 3D-STORM and shown in B-C. (B) Merge of widefield fluorescence image and reconstructed 3D-STORM image corresponding to the area indicated by the black arrow in A. 3D-STORM image is reconstructed, using initial form of BDP-650 excited at 640 nm (6.5 kW.cm^-2^) during 100 seconds of illumination at 37°C in physiological buffer (Krebs-Ringer). Green arrow indicates a mitochondrion shown at higher magnification in C. (C) DBScan segmentation of mitochondria (white linear network) and single molecules detected using 3D-STORM (the color code from purple to orange represents the depth), indicated in B. (D) Particles count over 5000 frames (100 seconds) at a frame rate of 20 ms. Graphic bars represent a bin of 100 frames. Purple arrows indicate transient pulse illumination (few sec) with 405 nm wavelength, at low density power (0.243 kW.cm^-2^), followed with an increase of single particles detected. Purple and brown windows in the graphic represent single particles accumulated during the first 500 frames and the last 500 frames of the acquisition. (E) Radial precision of single localizations detected during 3D STORM imaging cession. The mean of radial precision is 44 nm for more than 90 000 localizations. (F) Radial precision of single particles detected during the 500 first (purple) or last (brown) frames of the acquisition, indicated with purple and brown windows in D. During the first 500 frames (purple), we detected 29,935 single particles with a mean radial precision of 15 nm. For the last 500 frames (brown), we detected 1,167 single particles with a mean radial precision of 76 nm. A shift in radial precision can be seen between the first and the last frames.

Interestingly we also noticed that a brief (1-2 sec) addition of UV light at low power (0.243 kW.cm^-2^) induced an increase of detected particles per frame (Figure 7D purple arrows).

However, we could also see that the number of particles detected over time was decreasing, (Figure 7D).

Since 640 nm exposition could promote the switch from native to converted form this could lead inevitably to a diminished pool of available blinking fluorophores. We thus wondered if the 561 nm excitable converted form could be a better candidate for long blinking phase. To evaluate the blinking properties of both native **BDP-650** and converted form **cBDP-650** (Figure S20), we sequentially acquired the native **BDP-650-Mito** (640 nm laser line excitation) and the converted **cBDP-650-Mito** (561 nm) in interleaved mode, over 22,000 frames (10 ms as exposure over 11 cycles of 1,000 frames each). Our results showed that both forms of **BDP-650** are competent for STORM imaging without any reducing buffer. Surprisingly we noticed that low laser power (8.445 kW.cm^-2^ with 561 nm and 6.475 kW.cm^-2^ with 640 nm) for both forms was sufficient to make **BDP-650** blink in STORM imaging. Even more interestingly, **BDP-650** was capable of self-blinking continuously over 22,000 frames without any addition of UV light (405 nm excitation laser line). Finally, we showed that both forms of **BDP-650** labelled the same cellular structures: 561 and 640 nm signal sharing the same morphology and regions (Figure S20 B-C-D). Moreover, our results showed that the native form (λ_ex_= 640 nm) has a mean radial precision of 11 nm (Figure S20E), while the converted form (λ_ex_ = 561 nm) has a lower mean radial precision of 19 nm (Figure S20E), which is in line with the higher brightness of **BDP-650** compared to its converted form.

Since localization precision of native form (magenta in Figure S20E) was presenting two local maxima, we wondered if radial precision was decreasing over time. We thus measured radial precision for both forms in the first 500 frames (Figure S20G) and the later 500 frames (Figure S20H). This showed that the native form harbours a better radial precision which is decreasing along the time. Interestingly, the converted form (cyan in Figure S20FGH) had a lower precision, but could show a more sustainable blinking behaviour over the time (Figure S20F). We conclude that both forms of **BDP-650** can be used for live cell STORM imaging in physiological condition with a precision of localization in the same range than the one usually obtained on fixed samples with a reducing buffer. Altogether, these results demonstrate that the converted form of **BDP-650** can be used for STORM live imaging of cellular structures in physiological conditions, with a suitable resolution and adapted blinking time for live cell imaging.

As converted form **cBDP-650** (λ_ex_ = 561 nm) displayed satisfactory radial precision with a sustainable blinking, we decided to focus our work on this form to test live cell STORM imaging of the plasma membrane. Indeed, reconstructing the whole plasma membrane is challenging and require a huge amount of localizations in 3D. We thus used the membrane converted form of **BDP-650** to unravel the neuronal plasma membrane nanostructure (Figure 8A-D). Our results showed that **cBDP-650-PM** is remarkably effective. As a matter of fact, we were able to sharply reconstruct the entire organization of a complex neuronal network in only 3 min (Figure 8F) (10,750 frames with 20 ms as exposure time). Strikingly we could detect 247,225 particles which have a mean radial precision of 25.7 nm, (Figure 8E). In these conditions we could segment with high fidelity thin details of the plasma membrane (Figure 8CD). Unlike the mitochondrial version, we noticed that the membrane version showed a stable number of particles per frame (Figure 8F), without any shift in radial precision over time (Figure 8G). A transient application of low UV light (0.240 kW.cm^-2^ – few seconds, purple arrow in F) induced a sustainable increase of the number of particles detected for the rest of the acquisition (Figure 8F).

**Figure 8.**
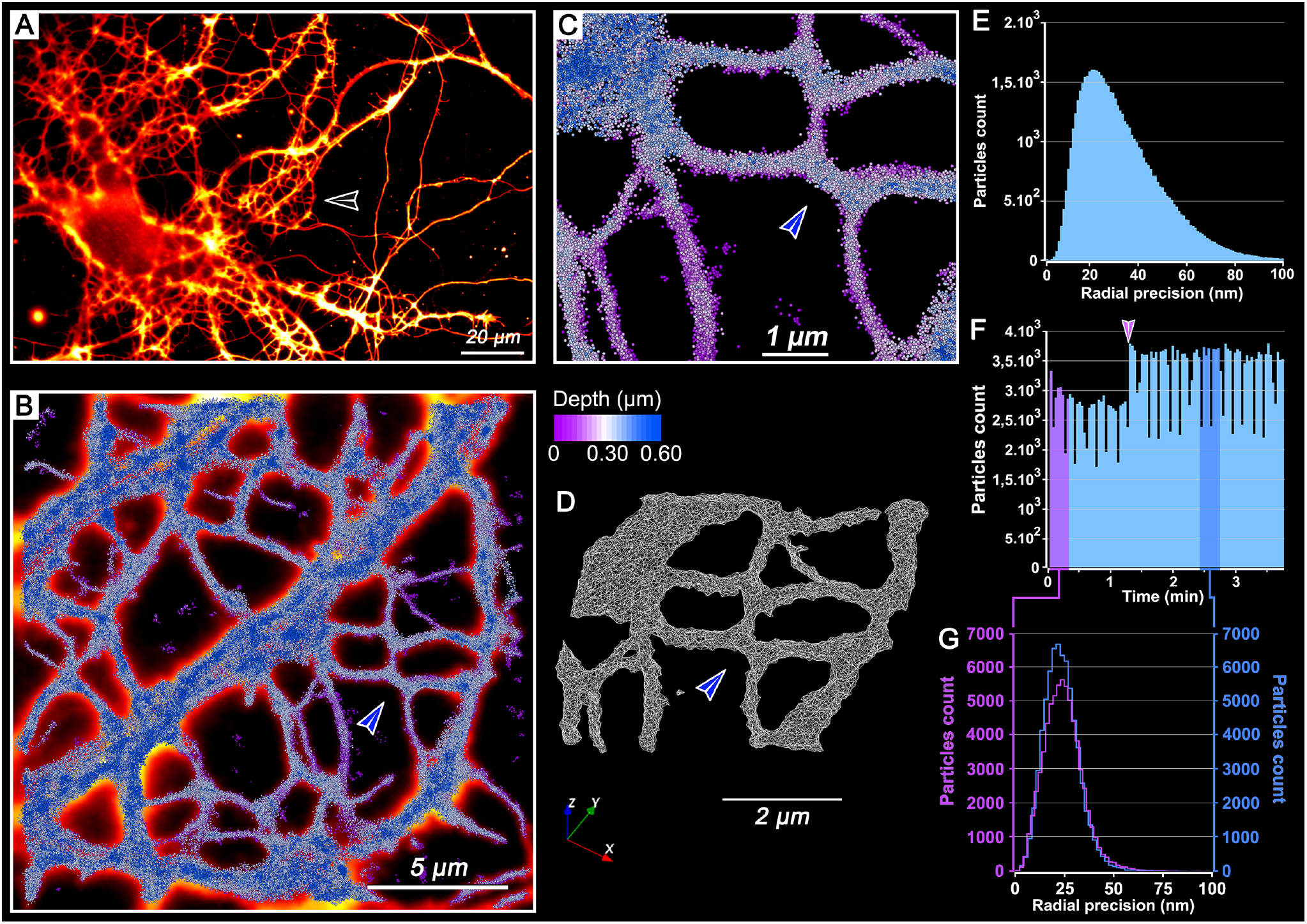
3D-STORM imaging of plasma membrane in living neuron using converted form cBDP-650-PM at 37°C in physiological buffer. (A) Widefield fluorescence image of membrane network in living 14 DiV-neuron labelled with cBDP-650-PM. Color code represents fluorescence intensity. Black arrow shows the area acquired in 3D-STORM represented in B-C-D. (B-C) Live-cell 3D-STORM imaging at 37°C in physiological buffer (Krebs-Ringer). Reconstruct 3D-STORM image, using converted form cBDP-650-PM, is made of 10,750 frames at a frame rate of 20 ms with 561 nm (8.445 kW.cm^-2^). Single particles are represented as spheres of 50 nm. (B) Merge of widefield fluorescence image (fire LUT) and reconstructed 3D STORM (blue gradient) image corresponding to the area indicated by the black arrow in A. (C) Expanded view of membrane network indicated with a blue arrow in B, which shows accurate reconstruction of the neuronal membrane. (D) Nanoscale mitochondrial wireframe obtained using DBScan 3D segmentation of membrane structure represented in C. Blue arrow is indicating the same intersection shown in B and C. (E) Radial precision of single particles detected during 3D-STORM imaging. The mean of radial precision is 26 nm for 247,225 single localizations. (F) Particles count over 10,750 frames (3 min 35 s) before and after short UV irradiation. Graphic bars represent a bin of 100 frames. Purple arrow indicates transient illumination with 405 nm wavelength (few seconds) at 0.240 kW.cm^-2^. Purple and blue windows in the graphic represent single particles accumulated during the first 1000 frames and from frames 7,500 to 8,500. (G) Radical precision of single particles detected during the first 1000 frames (purple): 66,377 single particles with a mean radial precision of 29 nm. Between frames 7,500 to 8,500 (blue), 67,734 single particles with a mean radial precision of 28 nm.

Since **cBDP-650-PM** was good enough to unravel the membrane architecture in a short time scale, we decided to challenge the probe on a thick sample over live Z-stack acquisition. To do so, we realized 3D-STORM imaging on a thick dendritic shaft of mature neuron (Figure 9). Without any UV light, we successfully imaged 1.5 μm thick dendritic shaft during 8 min (Figure 9E). As it can be seen in Figure 9A and B, we reconstructed with fidelity the structure of the dendritic membrane and even small plasma membrane protrusions emerging from the main dendrite (Figure 9C). Our results show that we detected 288,146 particles with a mean radial precision of 21 nm (Figure 9D). This acquisition highlights the efficiency and the robustness of **cBDP-650**. Indeed, we successfully made two Z-stacks in a row (Figure 9E where the second stack is indicated by the symbol ΣZ2) without bleaching **cBDP-650** during the first Z stack.

**Figure 9.**
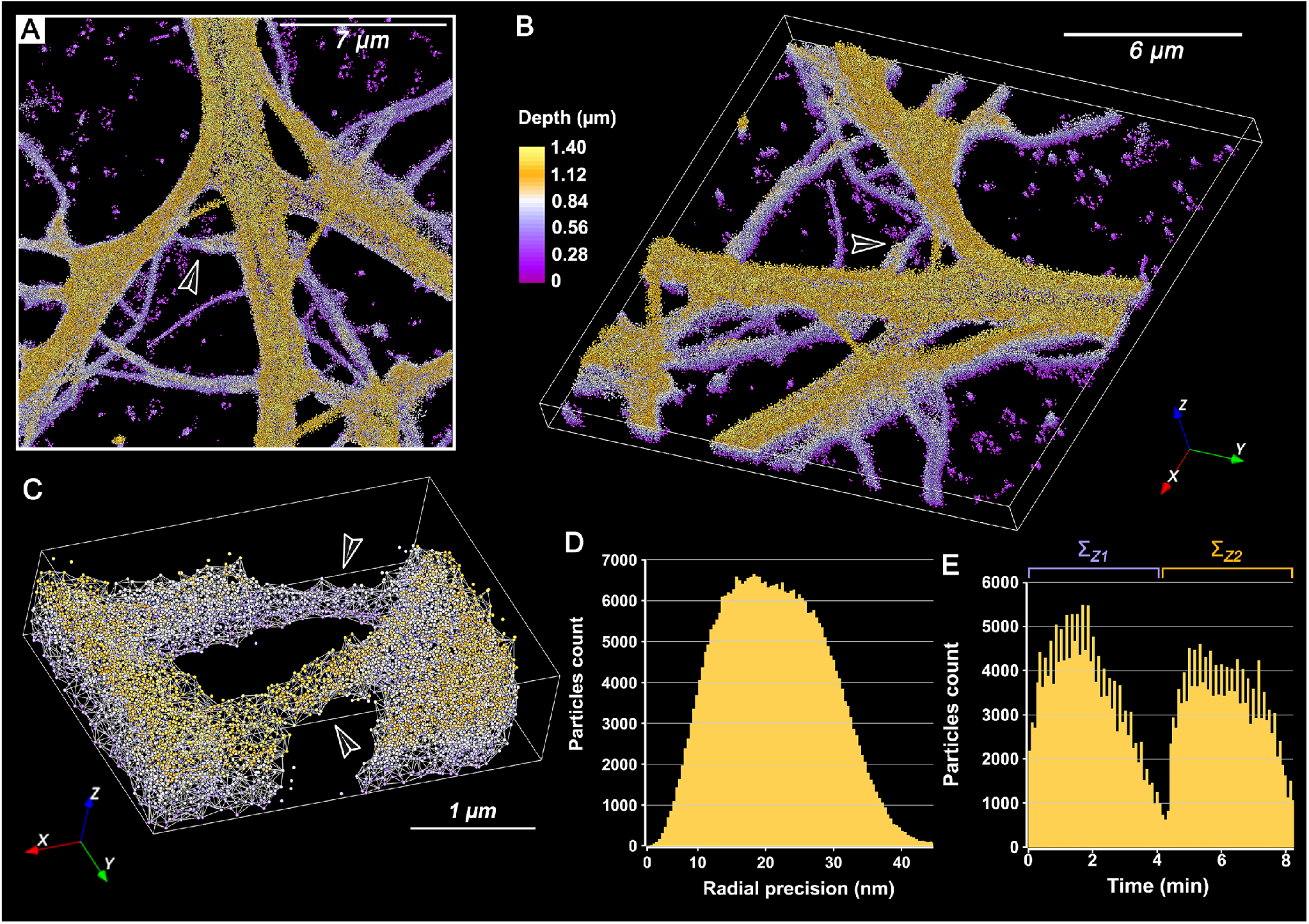
3D-STORM imaging of voluminous dendritic membrane of living neuron at 37°C in physiological buffer. (A-C) Live-cell 3D-STORM imaging, using converted cBDP-650-PM, at 37°C in physiological buffer (Krebs-Ringer) during 8 min without any addition of 405 nm wavelength. The final 3D-STORM image is made of 24,000 frames at a frame rate of 20 ms with 561 nm illumination (8.445 kW.cm^-2^). Single particles are represented as spheres with a diameter of 50 nm. The color code from purple to yellow represents the depth. To minimize photo damage, 2 Z-stacks of 16 slices (4 min each) were acquired successively. (A-B) Reconstructed 3D-STORM image with a thickness of 1.4 μm. The image is made up of two cycles of 12,000 frames which are distributed on 16 Z containing 750 frames each. (A) x,y view of 3D STORM image. (B) x,y,z view of 3D STORM image. (C) DBScan 3D segmentation (white network) of dendritic structure indicated in B, which shows two dendritic pathways: one above and one below, indicated by black arrows. (D) Radial precision of single particles detected during 3D-STORM imaging. The mean of radial precision is 21nm for 288,146 single particles. (E) Particles count during the acquisition of 24,000 frames. 2 Z-stacks of 16 pictures were acquired successively. Bars above named as ΛZ1 and Z2 represent the localization of the first and second Z-stack. Even if fewer single particles were detected during the second cycle, cBDP-650 seems to work efficiently during long and thick acquisition. Graphic bars represent a bin of 100 frames.

At last, we also checked that membrane blinking properties of **cBDP-650** was not affected by the lipid composition of plasma membrane. Indeed, lipid composition is known to change during neuronal maturation. We thus tested blinking properties on either very young neurons (Figure S21) or on conventional epithelial Hela cells (Figure S22). As shown on figure 8, 9, figure S21 and S22, **cBDP-650-PM** does not seem to be affected by lipid composition since different cell types and different maturation stage of the same cell type could be decipher without any issue. Since our results showed that **cBDP-650** can be used for more than 10 minutes in live imaging we wondered if we could combine dynamic studies with the spatial resolution of STORM imaging. To tackle this challenge, we used mitochondrial **cBDP-650-Mito** and tracked mitochondrial dynamics over two minutes (figure 10 and movie 2). Our results show that we successfully tracked moving mitochondria inside the dendrites (Figure 10. D1-D6 and E). Despites fast exposure time (10 ms), we could detect in the whole image 397,922 localizations with a mean radial precision of 28 nm (Figure 10C), and more specifically 10,390 localizations within the box (Figure 10 B, D1-D6). Finally, those results illustrate the efficiency of **cBDP-650** which allows to record dynamics of complex and thick cellular structures with both high spatial and temporal resolution.

**Figure 10.**
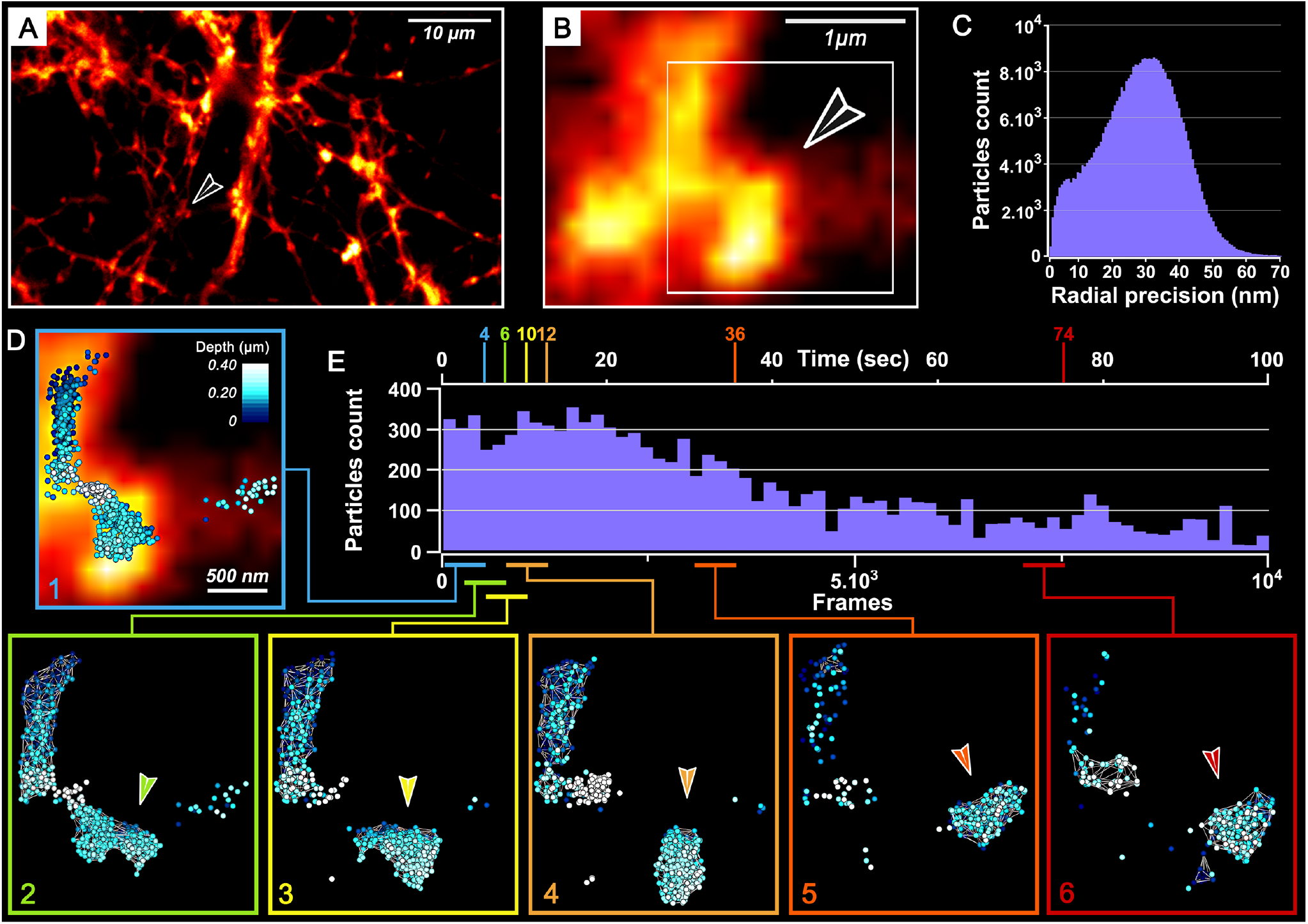
Dynamics of mitochondria displacement using cBDP-650 3D STORM imaging on live neuron. (A-B) Widefield fluorescence image of dendritic mitochondria in living 21 DiV-neuron using cBDP-650-Mito. Color code represents the intensity (Fire LUT). Black arrow indicates the expanded region of the mitochondria shown in B and D. (C) Radial precision of single particles detected during 3D-STORM imaging. The mean of radial precision is 21 nm for 288,146 single particles. (D) Live-cell 3D-STORM imaging of converted form of BDP-650 at 37°C in physiological buffer (Krebs-Ringer) during 10,000 frames at a rate of 10 ms with 561 nm illumination (8.445 kW.cm^-2^) and without any 405 nm irradiation. Single particles are represented as beads with a diameter of 50 nm. The color code from blue to white represents the depth. (D1-D6) 3D-STORM live reconstruction of single particles and their DBScan segmentation (white network) showing mitochondrial dynamics over 10,000 frames. Each time window corresponds to sliding combination of 500 frames (D1-D6), indicated by different colored box, below the frame scale of E. (D1) frames 0 to 500, (D2) frames 250 to 750, (D3) frames 500 to 1,000, (D4) frames 750 to 1,250, (D5) frames 2,500 to 3,000 and (D6) frames 7,000 to 7,500. (E) Single particles count (10,390) corresponding to the box indicated in B over 10,000 frames (100 seconds). Time (in sec) or Frame number are indicated respectively at the top and bottom of the graph. Frame time window are represented by colored horizontal line, corresponding to colored box indicated in (D1-D6). Graphic bars represent a bin of 100 frames.

To summarize our results, we have shown that both form of **BDP-650** can be used in live cell STORM imaging at 37°C without reducing buffer or UV light. Then we demonstrated that the conversion of **BDP-650** do not alter its own targeting properties. Lipid composition of the plasma membrane does not seem to influence blinking properties of **BDP-650. cBDP-650** shows robustness for long time imaging and amazing performances to reconstruct sharply every detail as well as voluminous structure of the plasma membrane.

Finally, we can conclude that **BDP-650** is a suitable tool for live imaging in STORM allowing both high spatial (20-30 nm as radial precision) and temporal resolution.

## Conclusions

Bright photoconverters adapted to microscopes settings including lasers lines and emission filters are highly valuable tools for tracking and monitoring dynamics of biological events by multicolor fluorescence imaging. Herein we showed that pyrrolyl-BODIPYs: **BDP-576** and **BDP-650** possess unique photoconversion features, displaying adapted rate of photoconversion, low phototoxicity, and providing bright and stable photoproducts that are perfectly adapted to typical channels of fluorescence microscopy. The photophysical properties of these probes were thoroughly investigated and led us to conclude that the involved mechanism of photoconversion was our recently reported Directed Photooxidation Induced Conversion (DPIC) mechanism. Hence, the rationalization of this mechanism and its exemplification from coumarin to BODIPY open new avenues to develop photoconverters based on DPIC. In addition to their appealing photoconversion properties our single molecule analysis showed that pyrrolyl-BODIPYs displayed impressive photoswitching properties providing numerous, fast and bright blinking in a short time frame. Although the photoswitching mechanism needs to be determined, we hypothesized that the fast reversibility observed at the single molecule level might be caused by transient formation of endoperoxide intermediates arising from the reversible addition of singlet oxygen. Regardless the mechanism, these impressive photoswitching features allowed live SMLM imaging of plasma membrane and mitochondria in epithelial cell as well as in fragile cells like neurons.

The complexity of SMLM imaging on living organisms is raised by all the parameters that we have to combined for optimal conditions. Here, we provided evidence showing that **BDP-650** is a tool with remarkable properties for live cell STORM imaging. Indeed, we were able to demonstrate that **BDP-650** can fit with physiological conditions such as Krebs-Ringer buffer at 37°C without any reducing buffer or UV light. Moreover **BDP-650** can be used with low laser power to avoid any cytotoxic side effects and cellular damaged. Strikingly, **BDP-650** harbour interesting properties allowing long time imaging and sharp reconstruction of thin details, within voluminous and complex organization of different cellular structures. We showed that pyrrolyl-BODIPYs can be combined with Halo Tag to benefit from genetic specificity. It can also be combined to targeting moieties, which makes the labelling of all primary cells (neuronal cells or immune cells known to be difficult to transfect), much more accessible. With 100% of labelled cells this strategy thus enabled stronger statistics on fragile live cells. On top of that it is suitable for high spatial resolution which perfectly fits with STORM imaging. Although **BDP-650** spans over two imaging channel (561 and 640 nm) that are widely used in STORM imaging, it can still be used with PA-GFP form during PALM acquisition. Finally, properties of **BDP-650** open a novel field for live cell STORM imaging. Indeed, it would be interesting in the next years to develop a native form of BDP excitable in the near infra-red channel (typically using 730 nm laser line). We thus think that a new generation of BODIPYs with improved properties will enable 3D live cell and multicolour STORM imaging.

## Materials and methods

### Synthesis

All starting materials for synthesis were purchased from Alfa Aesar, Sigma-Aldrich, or TCI Europe and used as received unless stated otherwise. pcDNA3-EGFP was purchased from addgene (Watertown, USA). NMR spectra were recorded on a Bruker Avance III 400 MHz or 500 MHz spectrometers. Mass spectra were obtained using an Agilent Q-TOF 6520 mass spectrometer. Absorption spectra were recorded on a Cary 4000 spectrophotometer (Varian). Fluorescence spectra were recorded on a Fluoromax-4 (Jobin Yvon, Horiba) spectrofluorometer. Emission measurements were systematically done at 20°C, unless indicated otherwise. All the spectra were corrected from the wavelength-dependent response of the detector. HPLC analyses were performed on Agilent 1260 Infinity II system equipped with Interchim PF5C18AQ-250/046 reversed phase column with gradual eluting from water to acetonitrile for 20 min at 1.5 mL/min. Protocols of synthesis of all new compounds are described in the Supporting Information. BDP-576 and BDP-650 were purchased from Lumiprobe GmbH (Hannover, Germany).

### Spectroscopy

The water used for spectroscopy was Milli-Q water (Millipore), and all the solvents were spectral grade. Absorption and emission spectra were recorded on a Cary 4000-HP spectrophotometer (Varian) and a FluoroMax-4 spectrofluorometer (Horiba Jobin Yvon) equipped with a thermo-stated cell compartment, respectively. For standard recording of fluorescence spectra, the emission was collected 10 nm after the excitation wavelength. All the spectra were corrected from the wavelength-dependent response of the detector. The fluorescence quantum yields φ_*F*_ were determined following the following equation:

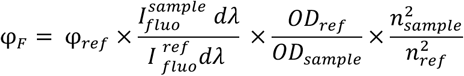

With OD the optical density at the excitation wavelength, and *n* refraction index of the solvent.

### Photoconversion

Laser spectroscopy and conversion were performed on 1 μM solution of dye in methanol using 3 × 3 mm optical path length quartz cuvettes of 45 μL. Excitation was provided by a continuous wave laser diode (488, 532 and 638 nm, Oxxius, Lannion, France) and the emission spectra were recorded by a QE pro spectrometer from Ocean Optics. All measurements were performed at room temperature. The kinetic rate of phototransformation, k_Pt_, was determined by fitting the emission decrease (integrated spectra) of the photo-converting dye over time as described in Moerner’s method,^25^ and according to the equation (1).

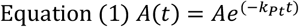

Where A(t) is the emission signal over time of the photo-converting dye.

Then the quantum yield of phototransformation ∅_Pt_ is given by the equation (3).

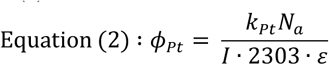

Where I is the irradiance at the sample (W.cm^-2^), ε the molar-absorption coefficient of the dye at the excitation wavelength (L.mol^-1^.cm^-1^) and N_a_ the Avogadro’s constant (mol^-1^).

Irradiance is defined by the equation (3).

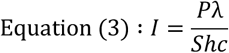

Where S is the surface irradiated (S = 0.15 cm^2^), h the Planck’s constant, c is the speed of light (m.s^-1^), P the power of the laser (in W) and λ the wavelength of the laser used for conversion (m). Based on our model, we hypothesized that cBDP-576-Mito was similar to BDP-FL and cBDP-650-Mito to BDP-SA (see SI for the comparison between the photoproducts and the models). The chemical yield (η) is obtained by comparing the fluorescence intensity of BDP before conversion (*Fl*_*A*_) and after conversion (*Fl*_*B*_) knowing Irradiance of the sample (I) and the brightness (B) defined as ε x φ_*F*_ at the laser’s wavelength of each dye by the equation (4).

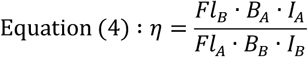

Finally, the quantum yield of photoconversion (φ_*pc*_) was obtained by multiplying the quantum yield of phototransformation (φ_*pt*_) by the chemical yield following the equation (5) and finally the quantum yield of photobleaching (φ_*Bl*_) is determined knowing φ_*pc*_ and φ_*pt*_ by the equation (6).

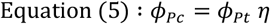

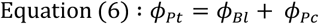

Consequently, when the dye is not convertible the quantum yields of phototransformation and photobleaching are linked following the equation (7).

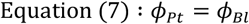

The photobleaching of the converted forms (cBDP-576-Mito) was performed at 488 nm (160 mW.cm^-2^) (after the photoconversion step) and the quantum yield of photobleaching was determined as described above.

### Singlet oxygen quantum yield

A solution of DPBF (100 μM) and the fluorophore or reference (5 μM) in MeOH were irradiated over few seconds with continuous wave laser. The emission spectra are acquired to obtain the slope of the decrease of DPBF fluorescence intensity. The singlet oxygen quantum yield is given by the following equation:

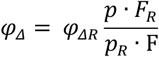

With p the decrease slope of the compound and F the corrected absorption given by the following equation:

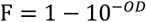

### Single molecule sample preparation

Coverslips were cleaned using argon plasma for 20 minutes followed by acidic cleaning using a solution of sulphuric acid and ammonium persulfate (100:1) under sonication for 30 minutes. Then, the coverslips were rinsed with MeOH three times and then incubated with a solution of MeOH, AcOH and APTES (20:1:0.6) for 30 minutes. The PEGylation was performed by incubating the coverslips in a solution of NaHCO_3_ 0.1 M with 10% of MeO-PEG-NHS (MW 5000 Da) and 0.2% of biotin-PEG-NHS (MW 5000 Da) overnight. Finally, a methanolic solution of biotinylated dye (1 pM) was added and allowed to remain for 5 minutes. The coverslips were rinsed three times with a MeOH.

### Single molecule analysis

Single molecule analysis was performed on a home-built setup based on a Nikon Eclipse Ti microscope with 100x 1.49 NA oil-immersion objective. The excitation was provided with a 488 nm, 561 nm and 642 nm laser lines (Oxxius). An acousto-optic tunable filter (AOTF; Opto-Electronic) was used to change the laser power. The signal was detected with on an EM-CCD camera from Hamamatsu (ImagEM). BDP-650-Biotin was excited at 642 nm (0.29 kW.cm^-2^) and 561 nm laser (0.33 kW.cm^-2^) without or after 642 nm irradiation with 30 ms of integration time for 90 s. BDP-576-Biotin was excited at 561 nm (0.33 kW.cm^-2^) and 488 nm laser (0.22 kW.cm^-2^) without or after 642 nm irradiation with 30 ms of integration time for 90 s. All images were analysed using ThunderSTORM plugin and a described protocol.^13^ Statistics were performed with homemade algorithm (see SI) to obtain the photon count, duty cycle, number of cycles, and the photoswitching time.

### Cell culture

Hela cells for photoconversion were incubated in Dulbecco’s Modified Eagle Medium (1 g·L^−1^ glucose) supplemented with 10% foetal bovine solution, 1% L-glutamine, and 1% antibiotic solution (penicillin−streptomycin) at 37 °C in a humidified atmosphere containing 5% CO_2_. KB cells were grown in Dulbecco’s modified eagle medium (DMEM, Gibco Invitrogen) and supplemented with 10% foetal bovine serum (FBS, Lonza), 1% L-glutamine (Sigma-Aldrich), 1% non-essential amino acid solution (Gibco-Invitrogen), and 1% MEM vitamin solution (Gibco-Invitrogen) at 37 °C in a humidified 5% CO_2_ atmosphere. Hela cells for SMLM were grown on poly-D-ornithine (Sigma Aldrich, P8638) coated #1.5 18 mm glass coverslips in DMEM (4.5 g/L^-1^ D-G-glucose) supplemented with 10% fetal bovine solution (Gibco™), 1% antibiotic solution (penicillin Streptomycin) and 1% Glutamax. They were incubated in humidified incubator with 5% CO_2_. Neurons were prepared from embryonic day 18 (E18) Sprague Dawley rat embryos. Brains were extracted from the embryos and hippocampi were dissociated in 0.25% trypsin solution at 37°C for 15 min and then seeded in MEM medium (Gibco™) supplemented with 10% horse serum, 1% Glutamax, 3% Glucose (0.2 g.L^-1^) and 1% antibiotic at 37 °C at an approximate humidity of 95–98% with 5% CO_2_. 50,000 cells were seeded in 12 well plates on poly-D-ornithine (1 mg/mL) (Sigma Aldrich, P8638) coated #1.5 (18 mm) glass coverslips (Menzel-Gläser, Thermo Scientific). After 4 h when the cells were attached to the substrate, the medium was replaced with Neurobasal-B27 medium conditioned previously on confluent glial feeder layer (neurobasal medium (Gibco) containing 2% B27 supplement (Gibco), and 500 μM L-Glutamine (Sigma-Aldrich). Cultures were maintained up to 3 weeks at 37 °C in a humidified atmosphere of 95% air and 5% CO_2_ to obtain mature hippocampal network. Neurobasal medium was conditioned overnight on a confluent astrocyte feeder layer. One third of the neuronal medium was then replaced with this fresh conditioned medium once a week.

### Synthesis of the plasmid coding for Halo-NPM1

The sequence coding for Halo tag was amplified from pSEMS-Halo7Tag-hFis (111136 Addgene) vector by using primers (5’-GCAATTCGATATGGGATCCGAAATCGGTACTGGCTTTCC-3’ and 5’-GGCCTCGAGATTAACCGGAAATCTCCAGAGTAG-3’).

After purification the PCR products were digested by Bam-HI/Xho restriction enzymes and inserted in pcDNA3.1 (zeo) vector. Obtained pcDNA-Halo was further digested with HindIII/Bam-HI restriction enzymes and ligated with the NPM insert (obtained by PCR amplification from eGFP-NPM (17578 Addgene) with our primers 5’-CCCAAGCTTCCACCATGGAAGATTCGATGGACATGG-3’ and 5’-CGGGATCCAAGAGACTTCCTCCACTGCC-3’, then digested with HindIII/Bam-HI). Final pcDNA-NPM-Halo was ligated and sequenced for verification.

### Confocal imaging

Cells were incubated with **BDP-576-Mito** or **BDP-650-Mito** (200 nM) in opti-MEM for 30 minutes. The cells were then washed with opti-MEM before being imaged. **BDP-576-PM** or **BDP-650-PM** (200 nm) was added in opti-MEM for 5 minutes before being imaged. Cells were imaged with a Leica TSC SP8 laser scanning confocal microscope with a 63× objective.

### Photoconversion *in cellulo*

The photoconversion were performed by zooming on a region of interest to be photoconverted. About 20 successive scans at 2% laser power were applied and the intensity histogram was checked until it shows no more diminution of the signal of the initial form. The photoconverted channel was acquired every 5 scans of the non-converted channel to monitor the increase of signal.

### Cells fixation

The cells were covered with 4% of paraformaldehyde in PBS for 15 minutes at room temperature. The solution was removed and the cells were washed with PBS. Then Triton (0.1%) was added for 10 minutes. Then the cells were washed with PBS. The fixed cells were incubated with **BDP-650-Actin** (200 nm) in opti-MEM for 30 minutes. The cells were then washed with opti-MEM before being imaged.

### BDP-650-Halo and transfection

Hela cells have been transfected 24 h after being seeded in Ibidi chambers. Transfection mix have been prepared by preparing extemporaneously solution A containing jetPEI^®^ (2 μL, Polyplus transfection^®^) in 150 mM NaCl (50 μL) and solution B of pDNA (1.6 μg) and pcDNA3-EGFP (0.2 μg) in 150 mM NaCl (50 μL). After 5 min at room temperature, solutions A and B were mixed thoroughly and were allowed to stay 5 min at room temperature. Transfection mix was added on cells with 1 mL growing media. Cells were incubated at 37 °C for 24 h. The medium was removed and the cells were washed twice with opti-MEM, then **BDP-650-Halo** (50 nM) was added in opti-MEM and incubated for 30 minutes. To remove non-specific labelling the medium was removed and washed twice with growing medium and left at 37°C for 1 hour. Then the medium was removed, washed twice with opti-MEM before imaging in opti-MEM. The plasmids coding for Halo-tagged proteins and used to target various cellular substructures were: Halo-NPM1: nucleophosmine 1 (Nucleolus); Halo-hFIS1: FIS1 (Mitochondria); pAG848 H2B-Halotag; LifeAct-Halotag (actin); pAG847 golgi-Halotag (Golgi apparatus).

### Cytotoxicity and phototoxicity assays

HeLa cells were incubated for 1 h with BDP-Mito (200 nM) in a 96 wells plate (6 wells per condition) and were washed in Opti-MEM. For “dark condition” the cells were not irradiated. For phototoxicity assay, the conversion was performed by irradiating each well using a 5× objective with a green LED (560/80 nm LED from X-cite 110 Led illumination system) for 5 min for BDP-576-Mito and MitoTracker Red, and a 4× objective with a far-red LED (628/40 nm, EVOSTM Light Cube, Cy5 2.0) 5 min for BDP-650-Mito and MitoTracker Deep Red. The cells were then incubated at 37°C for 1 h before performing cell viability test using the 3-(4,5-Dimethylthiazol-2-yl)-2,5-Diphenyltetrazolium Bromide (MTT) assay. The medium was removed and MTT (Sigma Aldrich) at 0.5 mg/mL in PBS was added to cells that were further incubated for 3 hours. After the formation of the subtract to a chromogenic product by metabolically active cells, the medium was removed and 100 μL of DMSO was added to solubilized the crystals. The absorbance of each samples was measured with a plate reader (TECAN) spectrometer at 570 nm. Cell viability was reported as relative decrease compared to the absorbance of the positive control (Cells without BDP and light) considered as 100% of viable cells and negative control (Cells with Triton 0.1%) considered as 0% of viable cells. Data are presented as mean ± standard error of the mean of six independent experiments. ns: non-significative, ****p<0.0001.

### Live Single Molecule Localization Microscopy

BDP-650-Mito: Cells were incubated with BDP-650-Mito in cell media at pH 7.42 and 37°C during 2h, then washed 3 times with Krebs-Ringer solution (Krebs-Ringer solution: NaCl: 7.90 g/L, KCl: 0.37 g/L, CaCl_2_·2H_2_O: 0.13 g/L, MgCl_2_·6H_2_O: 0.24 g/L, NaH_2_PO_4_·H_2_O: 2.07 g/L, D(+)-Glucose monohydrate: 5 mM and at pH 7.42, warmed at 37°C with HEPES) and finally mounted in chamber (cf. STORM imaging) with Krebs-Ringer solution. For membrane version of BDP-650, BDP-650-PM: Cell media was removed and cells were washed 3 times with Krebs-Ringer. The cells were incubated during few minutes with BDP-650-PM in Krebs-Ringer and temperature at 37°C. Cells was washed 3 times in a row with Krebs-Ringer solution warmed at 37°C and then mounted in Chamlide chamber (cf. STORM imaging).

SMLM imaging was performed on a Vutara biplane microscope (Bruker) with a high-numerical aperture (NA) objective (60×, silicon, NA 1.3, Olympus). The 488, 561, 640 laser lines were all 1,000 mW. The 405 nm was 100 mW. 170 nm coverslips (Menzel glaser 18 mm diameter #1.5) were mounted in Chamlide chamber for 18 mm coverslips. Chamlide chamber were filled with imaging physiological buffer Krebs-Ringer solution warmed at 37°C placed in chamber at 37°C and imaged. Neurons were isolated with widefield microscopy (Coolsnap camera) and then imaged for STORM with a FLASH4 CMOS camera (20 × 20 microns). Irradiance was measured for each laser line (640, 561, 405 nm) with a Thorlabs powermeter. The probe (S121C) was set in place of the sample behind a #1.5 glass coverslip at the exit of the 60X objective covered with silicon oil. 3D-STORM imaging was done using the bi-plane module allowing the localization in the xyz direction. Sample were illuminated with: a 405 nm laser during experiment Figure 7 and 8 only, 640 nm laser for imaging BDP-650 in its non-converted form and 561 nm laser for imaging BDP-650 in converted form (cBDP-650). The SRX software (Bruker) was used to localize particles in 3D, 3D STORM images reconstruction and DBScan segmentation. Graphic of radial precision and particles count per frames were extract from SRX software.

## ASSOCIATED CONTENT

### Supporting Information

Characterizations (^1^H NMR ^13^C NMR, HPLC-High-resolution mass spectrometry) of the synthesized probes can be found in supporting information as well as additional experiments, controls, algorithm for single molecule analysis, single molecule analysis, cytotoxicity and photocytotoxicity assays, spectroscopy, analysis of photoproducts, signal/noise image analysis, photoconversion of live cells, live SMLM supplementary figures.

**Movie 1**. Tracking of single intracellular vesicles over 21 min.

**Movie 2**. Tracking mitochondrial dynamics in live SMLM imaging over 2 min.

## AUTHOR INFORMATION

### Author Contributions

LS synthesized the probes, performed all the spectroscopic experiments, the single molecule characterizations *in vitro* and the cell conversion experiments. PD supervised the kinetics studies and single molecule analysis. TL synthesized the plasmid corresponding to Halo-NPM1. VB and LD realized SMLM imaging in both neuronal and epithelial cells and analyzed biophysical properties of probes within living cells. LD and MC supervised the work. MC wrote the manuscript together with LD and VB for the SMLM imaging part. LD and MC founded the project. / All authors have given approval to the final version of the manuscript. / ‡These authors contributed equally.

## ACKNOWLEDGMENT

The authors would like to thank Pr. Arnaud Gautier for having provided plasmids coding for Halo tagged proteins: pAG847 Golgi-Halotag and LifeAct-Halotag, Dr. Halina Anton for the protocol of. Halo-NPM1, Dr. Andrey Klymchenko and Dr. Sophie Martin for granting access to equipment. We would like to thank Philippe Bun, David Geny, Laurianne Beynac for maintenance of NeurImag systems and Julie Nguyen and David Boulet for having provided hippocampal neurons.

