## Supplementary Information for "Targeted Photoconvertible BODIPYs Based on Directed Photooxidation Induced Conversion for Applications in Photoconversion and Live Super Resolution Imaging"

### Synthesis protocol and characterizations.

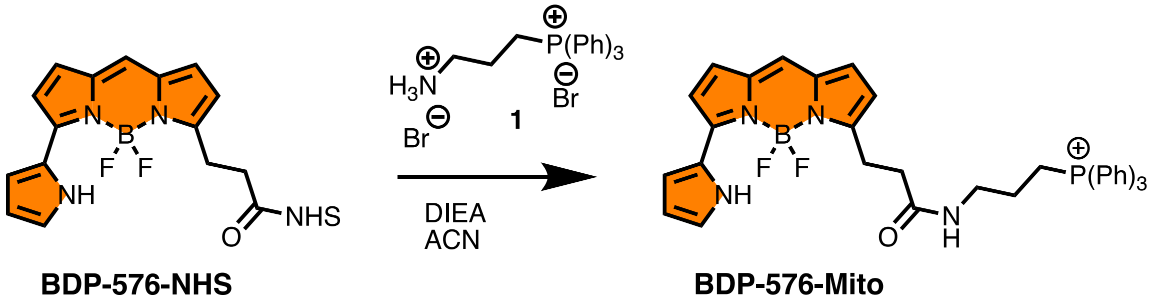


**BDP-576-Mito**. To a solution of BDP-576-NHS (11.7 mg, 24 µmol, 2 eq) and Diisopropylethylamine (30 µL, 171 µmol, 14 eq) in ACN (1 mL) was added 1 (5 mg, 12 µmol, 1 eq). The reaction mixture was left to stir at RT for 1 hour. The crude was purified was preparative TLC (DCM/MeOH: 9/1) to give BDP-576-Mito. Rf = 0.33 (DCM/MeOH: 9/1). HRMS (ESI+) calculated for $C_{37}H_{35}BF_2N_4OP$ [M]+ 631.2610, found 631.2589.

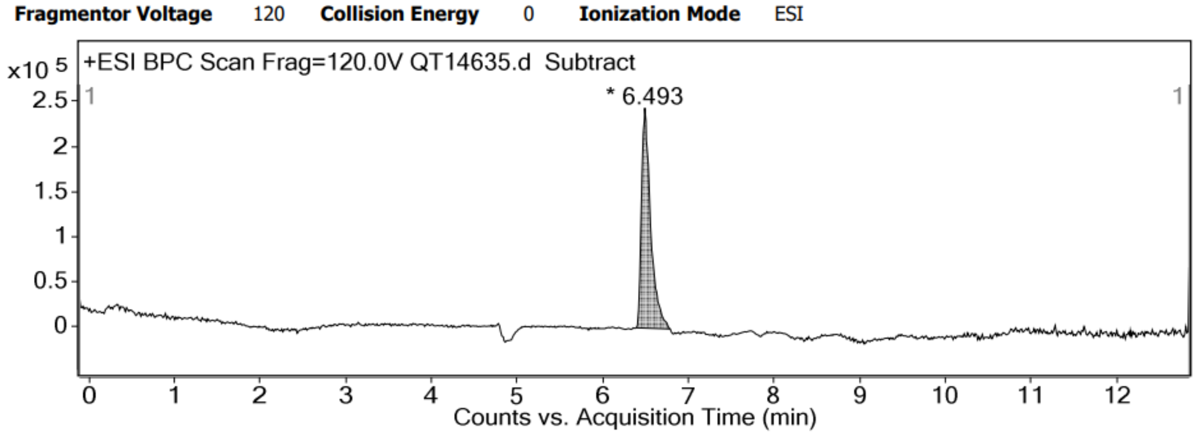


HPLC trace of **BDP-576-Mito**

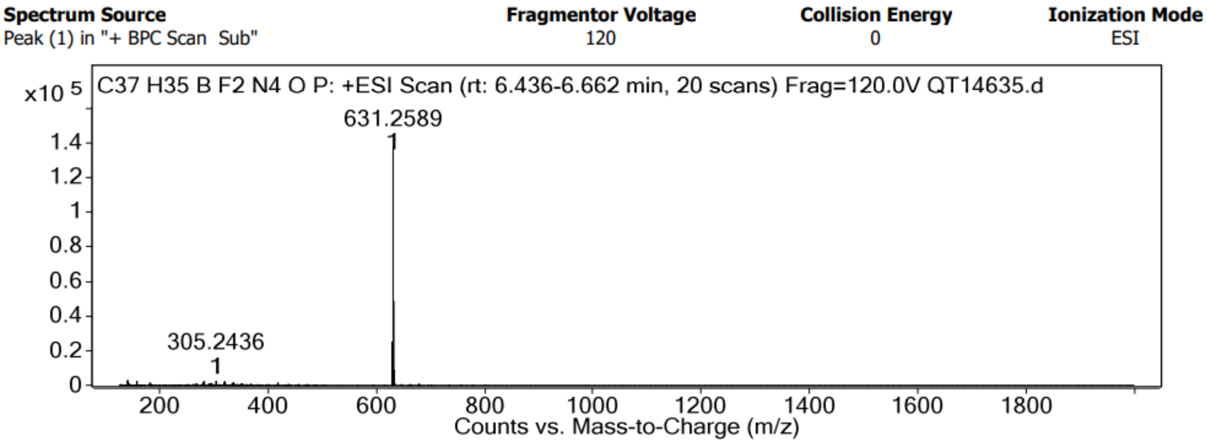


HRMS spectrum of **BDP-576-Mito**

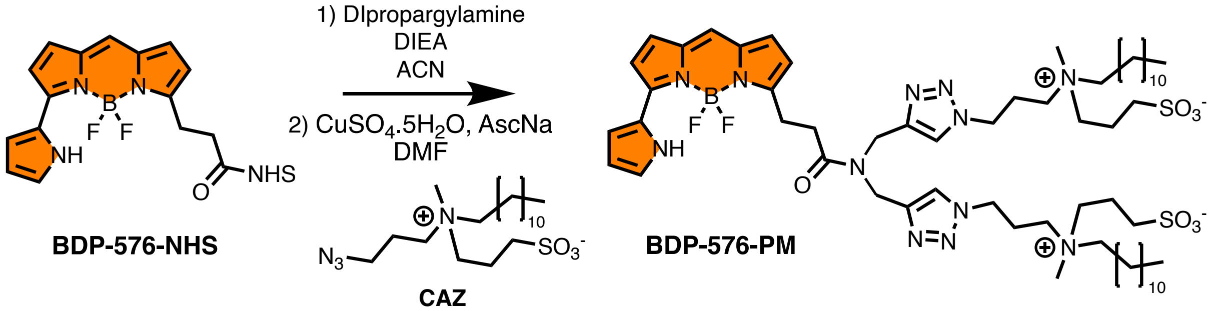


**BDP-576-PM.** To a solution of BDP-576-NHS ester (5 mg, 23.5 µmol, 1 eq) in ACN (3 mL) was added dipropargylamine (2.4 µL, 23.5 µmol, 1 eq). The reaction mixture was left to stir at RT for 4 hours. The crude was concentrated under reduced pressure and solubilized in DMF (3 mL) with **CAZ**[1] (21 mg, 51.7 µmol, 2.2 eq). 200 µL of an aqueous solution of $CuSO_4.5H_2O$ (7.2 mg, 30 µmol, 1.3 eq) and ascorbic acid (6 mg, 36 µmol, 1.5 eq) was then added. The reaction mixture was left to stir at 50°C overnight. The solvent was evaporated and the crude was purified by size exclusion column using DCM/MeOH, 1/1 to give **BDP-576-PM** after evaporation. HRMS (ESI+) calculated for $C_{60}H_{98}BF_2N_{12}O_7S_2Na$ $[M+Na]^+$ 1234.7082, found 1235.7220.

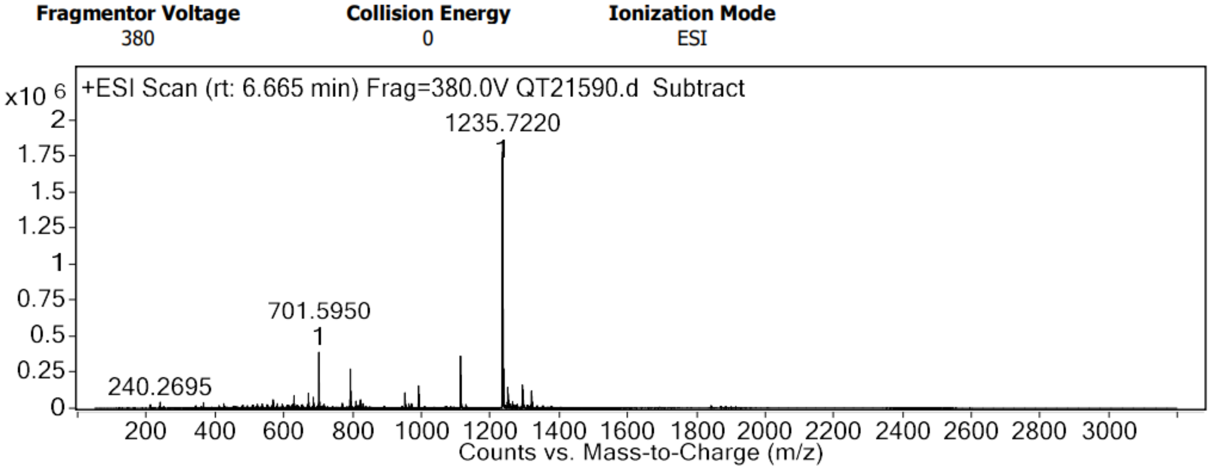


HRMS spectrum of **BDP-576-PM**

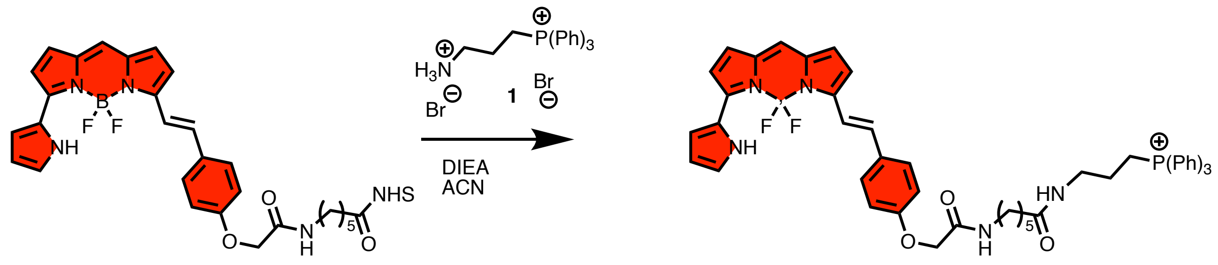


**BDP-650-Mito.** To a solution of BDP-650-NHS ester (5 mg, 7.8 µmol, 1 eq) and **1** (7.3 mg, 15 µmol, 1.9 eq) in ACN (1 mL) was added diisopropylethylamine (20 µL, 117 µmol, 15 eq). the reaction mixture was left to stir at RT for 4 hours. The crude was concentrated under reduced pressure and purified by column chromatography on silica gel (DCM/MeOH: 9/1 to 8/2) to obtain **BDP-650-Mito** (4 mg, 55%). $^1$H NMR (400 MHz, $CDCl_3$) δ 10.45 (s, 1H, NH Pyrrole), 8.65 (s, 1H, NH Amide), 7.82 – 7.62 (m, 15H, $PPH_3$), 7.58 – 7.47 (m, 3H, ArH), 7.21 (d, J = 9.7 Hz, 1H, ArH), 7.05 – 6.83 (m, 9H, ArH), 6.37 (d, J = 6.3 Hz, 1H, ArH), 4.52 (s, 2H, H Amide), 3.71 (d, J = 12.5 Hz, 2H, H Amide), 3.45 (d, J = 4.3 Hz, 2H, H Amide), 3.34 (q, J = 6.9 Hz, 2H, H Amide), 2.41 (d, J = 7.8 Hz, 2H, H $CH_2PPH_3$), 1.75 – 1.54 (m, 8H, $CH_2$). HRMS (ESI+) calculated for $C_{50}H_{50}BF_2N_5O_3P$ $[M]^+$ 848.3712, found 848.3741.

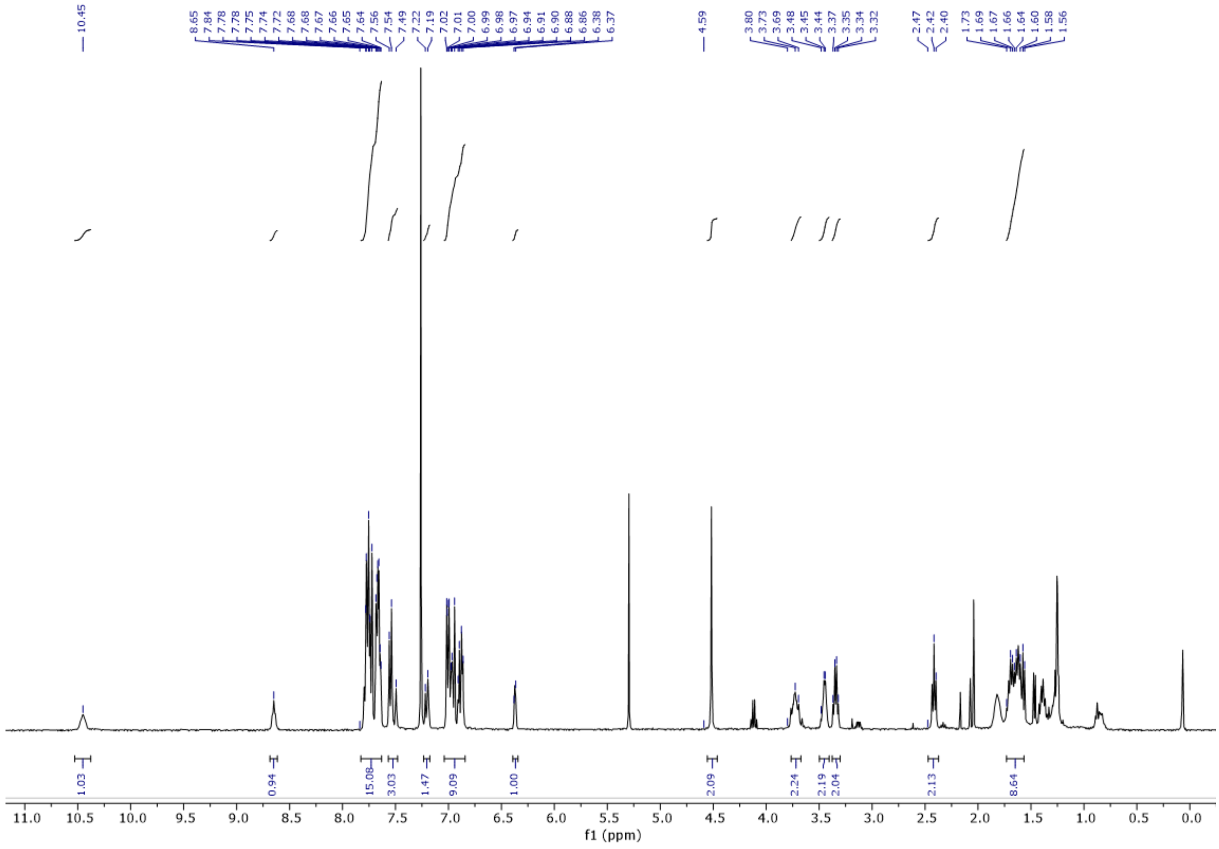


$^1$H NMR spectrum of **BDP-650-Mito**

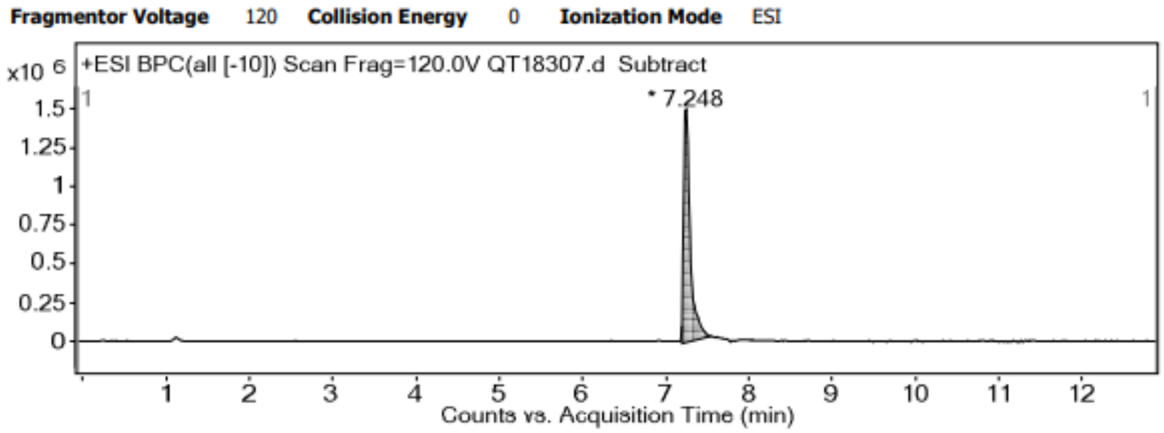


HPLC trace of **BDP-650-Mito**

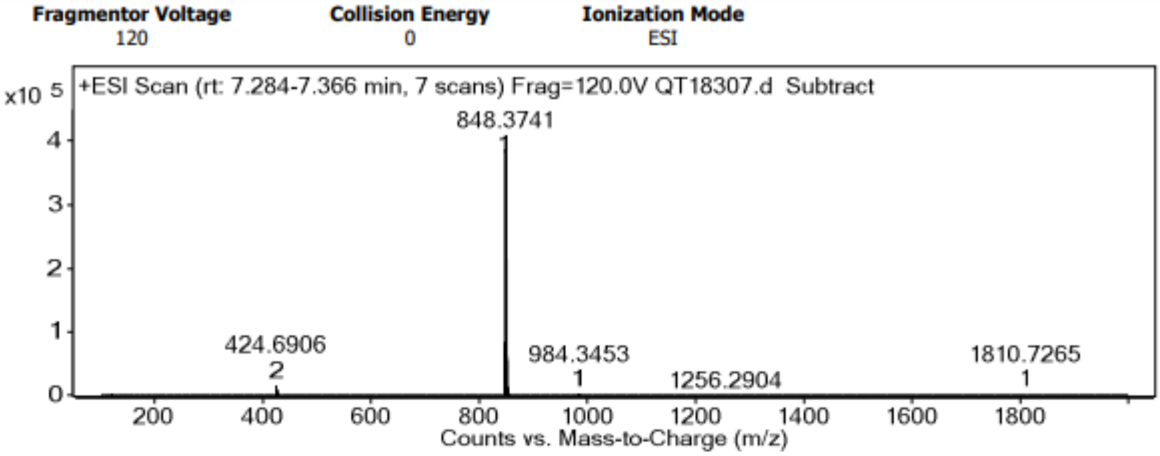


HRMS spectrum of **BDP-650-Mito**

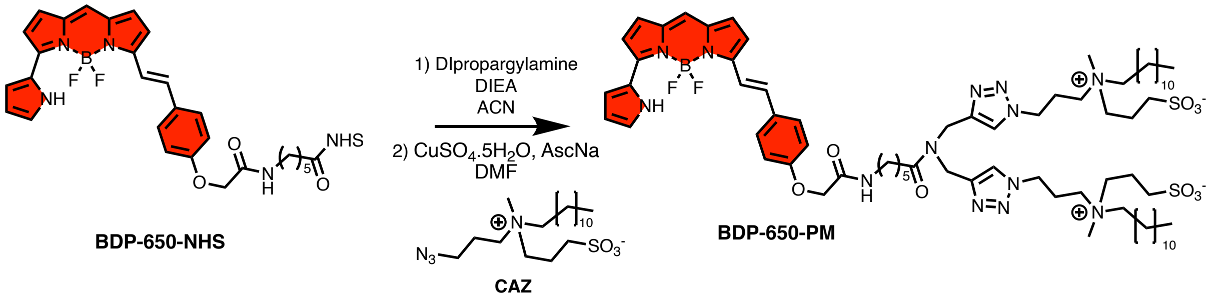


**BDP-650-PM.** To a solution of BDP-650 NHS ester (5 mg, 7.8 µmol, 1 eq) in ACN (3 mL) was added dipropargylamine (0.8 µL, 7.8 µmol, 1 eq). The reaction mixture was left to stir at RT for 4 hours. The crude was concentrated under reduced pressure and solubilized in DMF (3 mL) with **CAZ** (7 mg, 17.4 µmol, 2.2 eq). 200 µL of an aqueous solution of $CuSO_4.5H_2O$ (2.4 mg, 10 µmol, 1.3 eq) and ascorbic acid (2 mg, 12 µmol, 1.5 eq) was then added. The reaction mixture was left to stir at 50°C overnight. The solvent was evaporated and the crude was purified by size exclusion column using DCM/MeOH, 1/1 to give **BDP-650-PM** after evaporation (400 µg, 4%). HRMS (ESI+) calculated for $C_{73}H_{115}BF_2N_{13}O_9S_2$ $[M+H]^+$ 1430.8443, found 1430.8416.

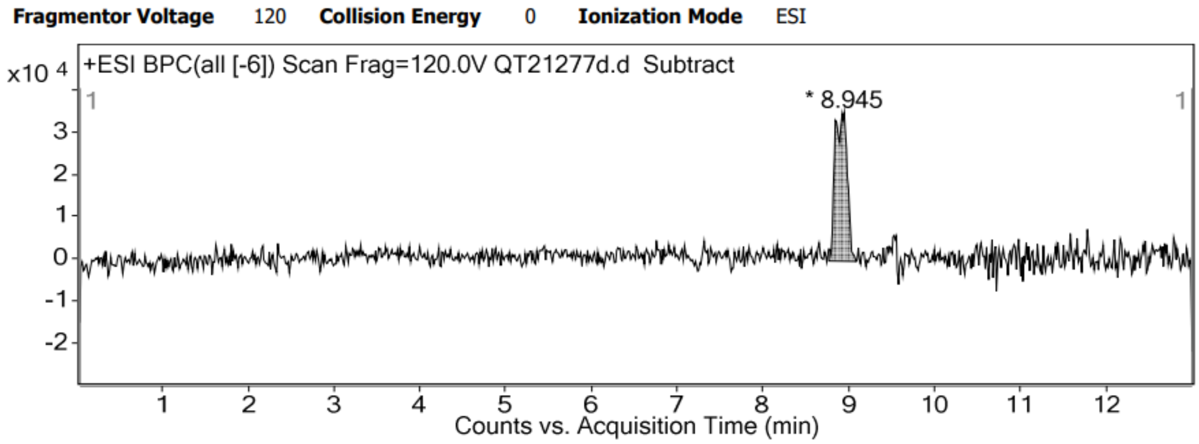


HPLC trace of **BDP-650-PM**

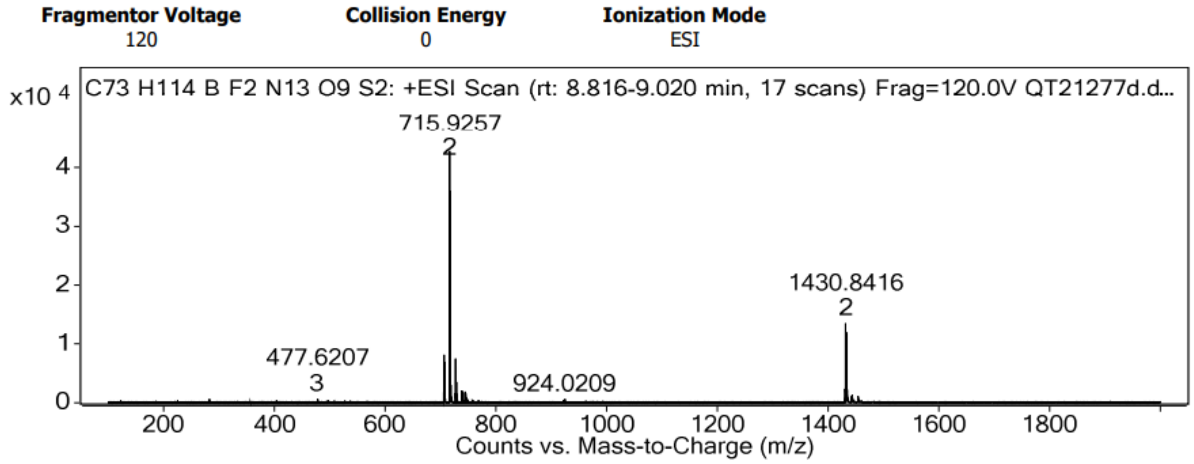


HRMS spectrum of **BDP-650-PM**

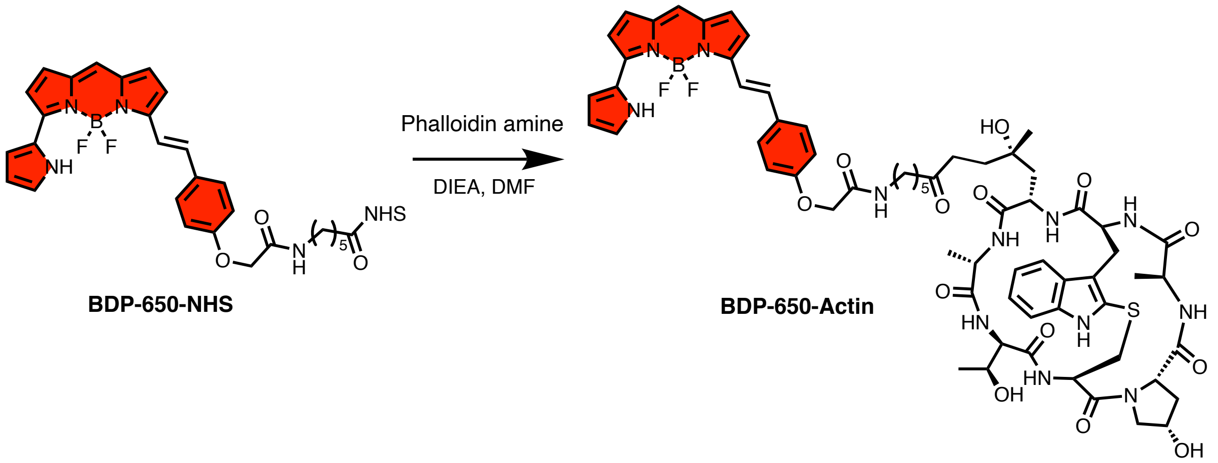


**BDP-650-Actin**. To a solution of BDP-650-NHS ester (1 mg, 1.3 µmol, 1 eq) and phalloidin amine (800 µg, 1.3 µmol, 1 eq) in DMF (1 mL) was added diisopropylethylamine (3 µL, 19 µmol, 15 eq). the reaction mixture was left to stir at RT for 4 hours. The crude was concentrated under reduced pressure and purified by HPLC ($H_2O$/ACN: 95/5 to 5/95) to obtain **BDP-650-Actin** (1.2 mg, 72%). HRMS (ESI+) calculated for $C_{64}H_{77}BF_2N_{13}O_{13}S$ $[M+H]^+$ 1313.5546, found 1316.5518.

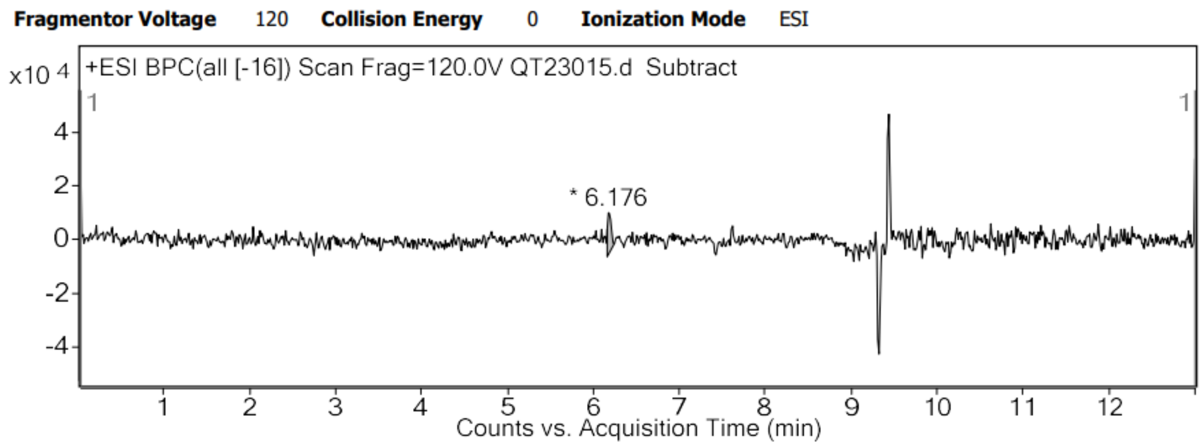


HPLC trace of **BDP-650-Actin**

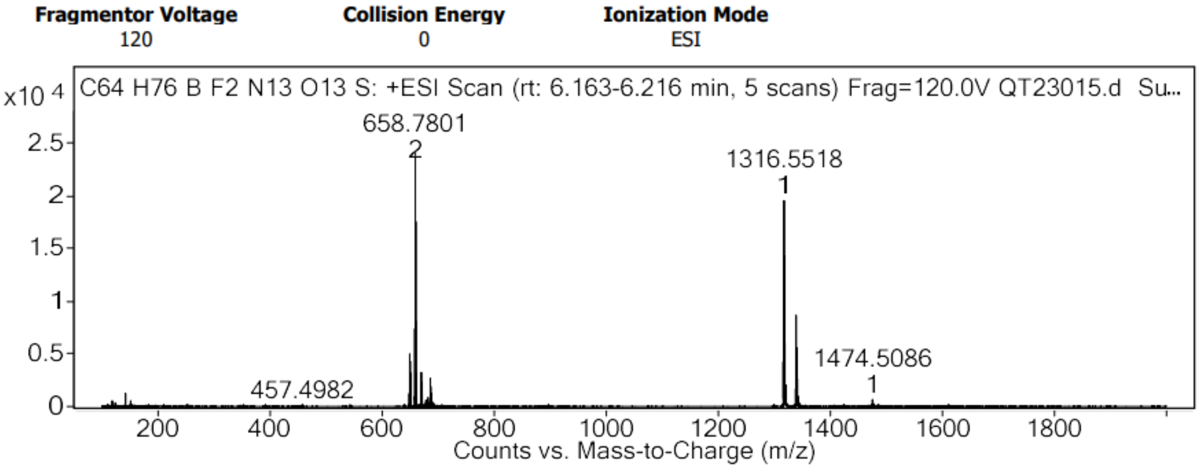


HRMS spectrum of **BDP-650-Actin**

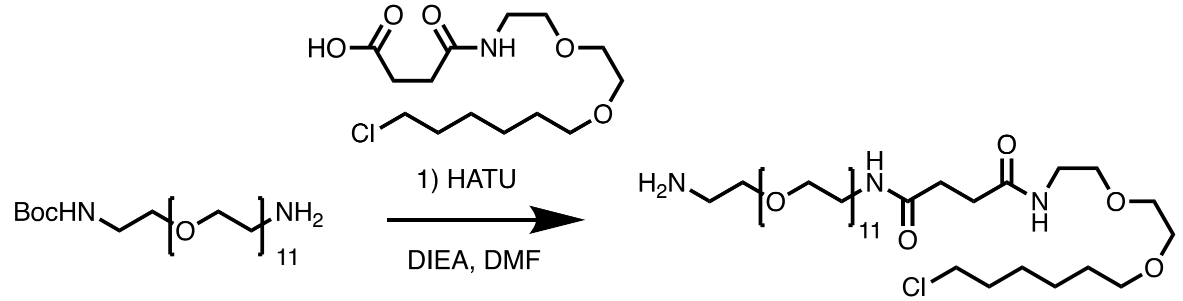


**Boc-*N*-amido-PEG$_{12}$-amine** **PEG$_{12}$-Halo Tag**

**PEG$_{12}$-Halo Tag**. To a solution of Boc-*N*-amido-PEG$_{12}$-amine (100 mg, 154 µmol, 1 eq) in DMF (2 mL) was added 4-[[2-[2-[(6-Chlorohexyl)oxy]ethoxy]ethyl]amino]-4-oxo-butanoic Acid (50 mg, 154 µmol, 1 eq), HATU (76 mg, 201 µmol, 1.3 eq) and DIEA (81 µL, 463 µmol, 3 eq). The solution was allowed to stir for 3 h at RT. The crude was concentrated under reduced pressure and purified by column chromatography on silica gel (DCM/MeOH: 9/1 to 8/2). Surprisingly, the **PEG$_{12}$-Halo Tag** was obtained (50 mg, 38%) as free amine, where the Boc protecting has been removed. $^{1}$H NMR (400 MHz, $CDCl_3$) δ 7.90 (s, 2H), 6.90 (t, J = 5.6 Hz, 1H), 6.64 (t, J = 5.6 Hz, 1H), 3.82 – 3.75 (m, 2H), 3.67 (dd, J = 5.6, 2.7 Hz, 2H), 3.65 – 3.46 (m, 50H), 3.46 – 3.33 (m, 6H), 3.15 (t, J = 4.9 Hz, 2H), 2.48 (s, 4H), 1.80 – 1.68 (m, 2H), 1.57 (p, J = 6.8 Hz, 2H), 1.48 – 1.27 (m, 4H). $^{13}$C NMR (101 MHz, $CDCl_3$) δ 71.23, 70.37, 70.35, 70.29, 70.26, 70.21, 70.19, 70.11, 70.05, 70.01, 69.99, 69.96, 69.87, 69.82, 69.69, 67.07, 45.03,

39.98, 39.21, 32.50, 31.68, 31.55, 29.42, 26.65, 25.38. HRMS (ESI+) calculated for $C_{38}H_{77}ClN_3O_{15}$ $[M+H]^+$ 850.5043, found 850.5036.

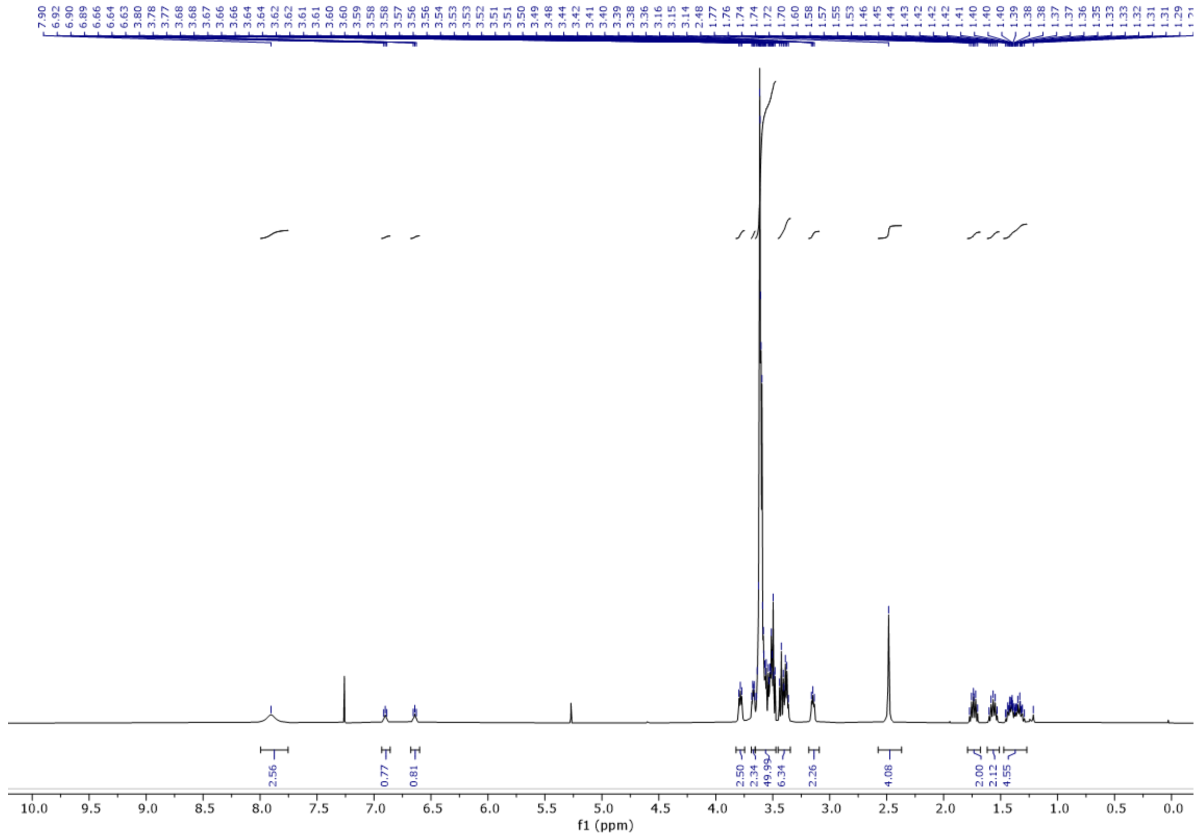

$^1$H NMR spectrum of **PEG$_{12}$-Halo Tag**

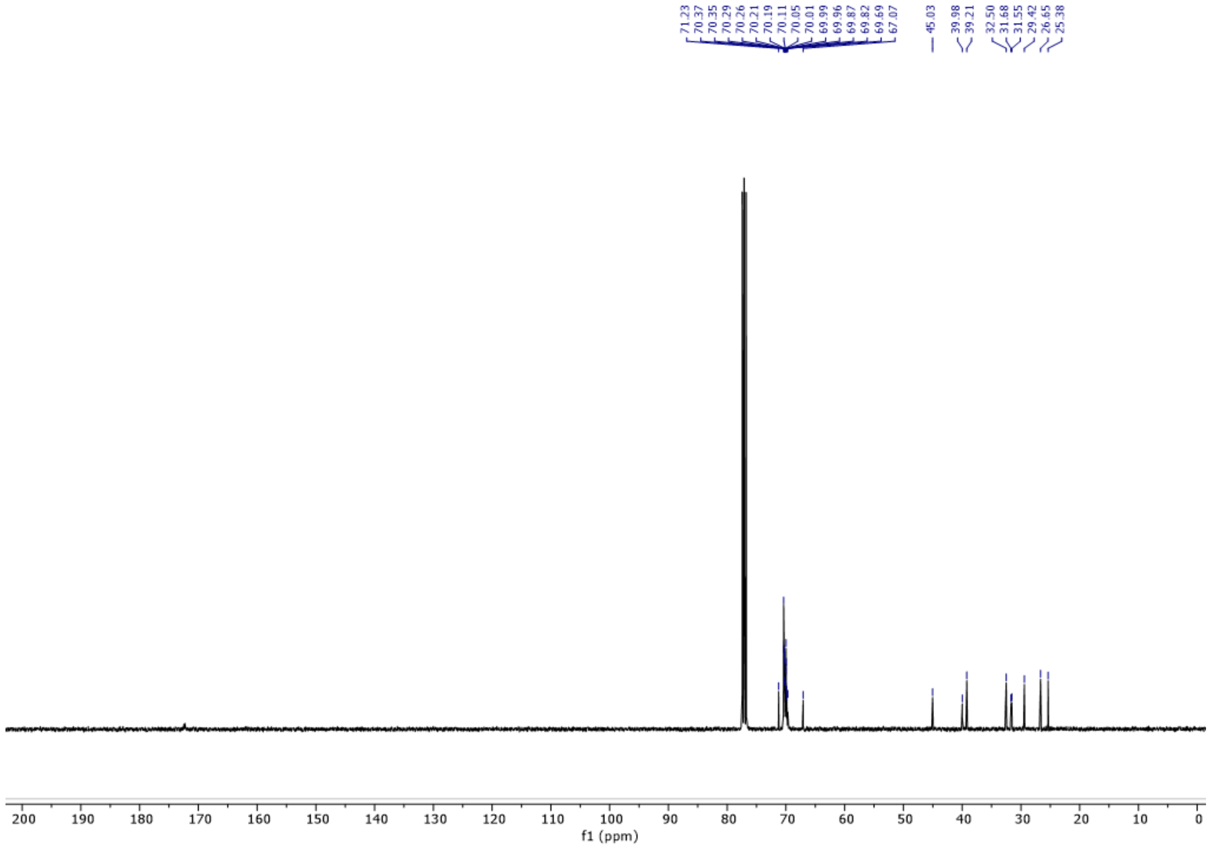

$^{13}$C NMR spectrum of **$PEG_{12}$-Halo Tag**

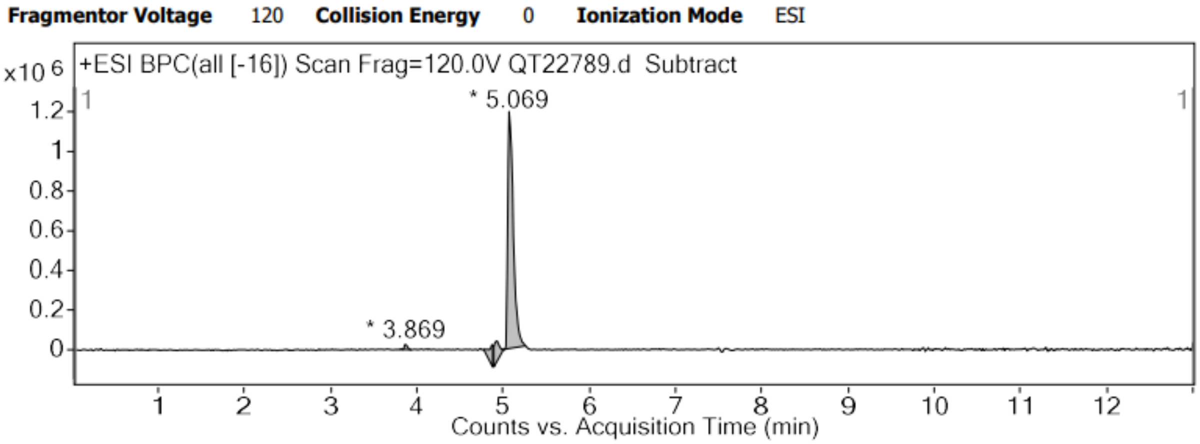


HPLC trace of **$PEG_{12}$-Halo Tag**

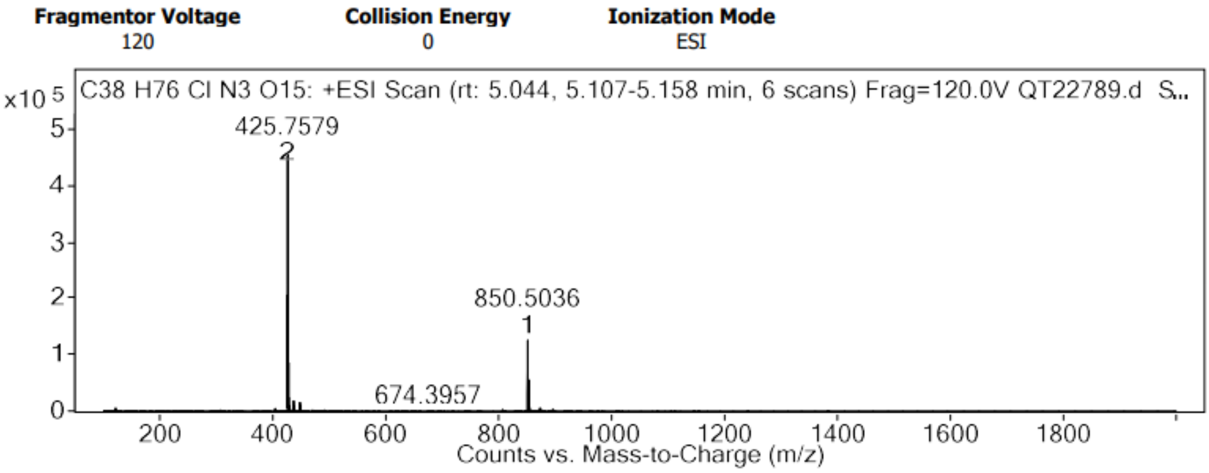


HRMS spectrum of **$PEG_{12}$-Halo Tag**

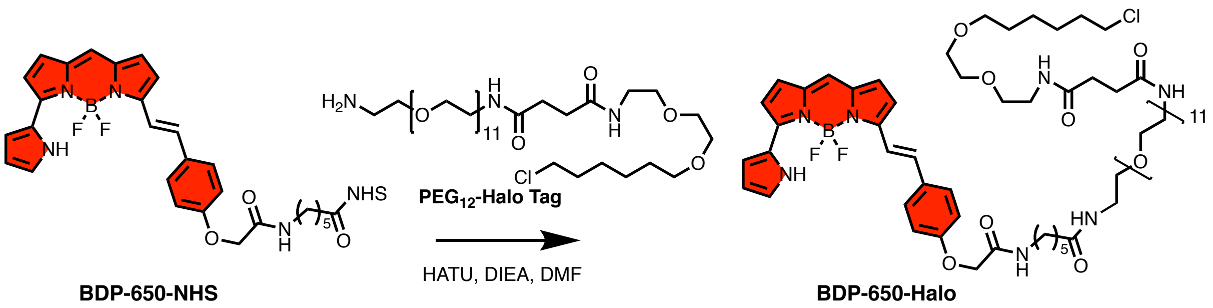


**BDP-650-Halo**. To a solution of BDP-650-NHS (5 mg, 8 µmol, 1 eq) in DMF (3 mL) was added **$PEG_{12}$-Halo Tag** (6.6 mg, 8 µmol, 1 eq), HATU (3.8 mg, 10 µmol, 1.3 eq) and DIEA (4 µL, 23 µmol, 3 eq). The solution was left to stir for 3 hours. The crude was concentrated under reduced pressure and purified by column chromatography on silica gel (DCM/MeOH: 9/1) to obtain **BDP-650-Halo** (1.5 mg, 14%). HRMS (ESI+) calculated for $C_{67}H_{103}BClF_2N_7NaO_{18}$ $[M+Na]^+$ 1400.7007, found 1400.6966.

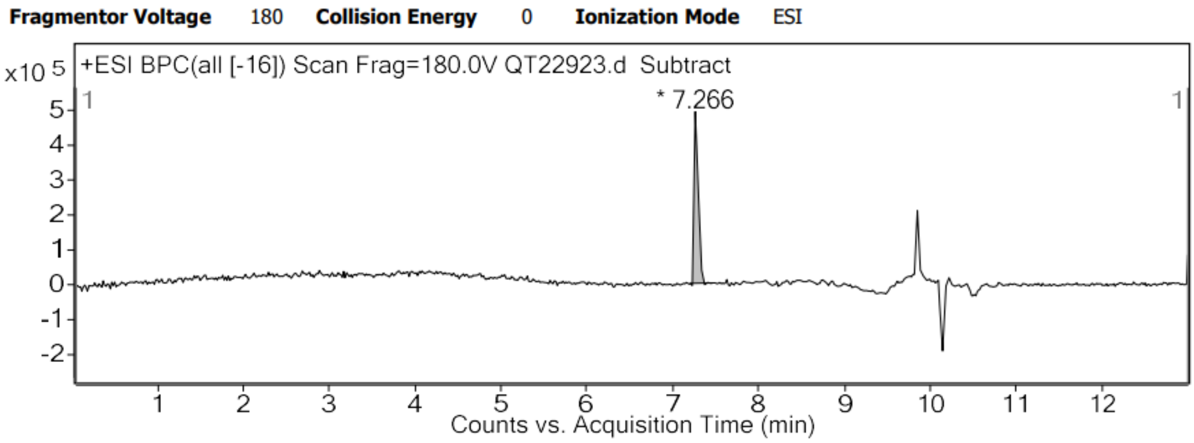


HPLC trace of **BDP-650-Halo**

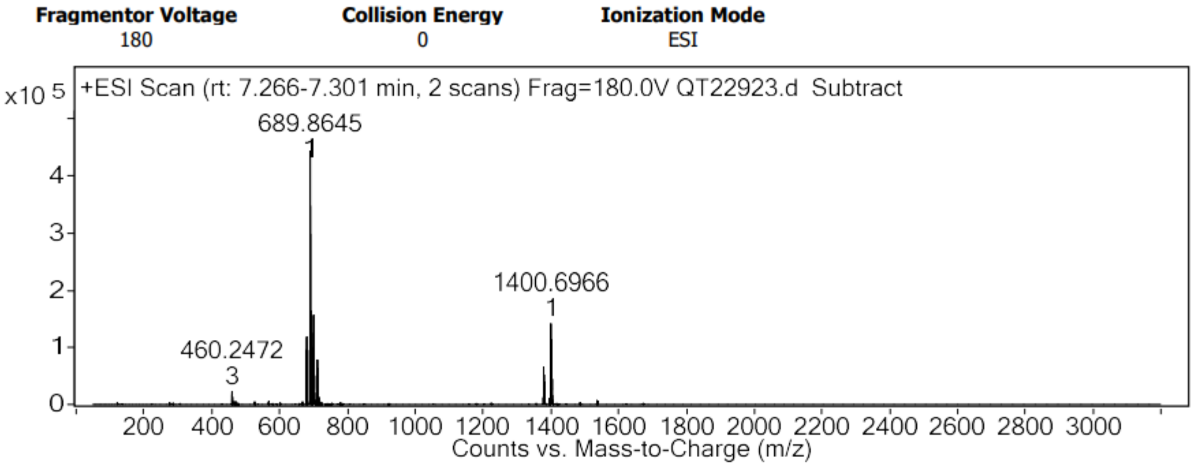


HRMS spectrum of **BDP-650-Halo**

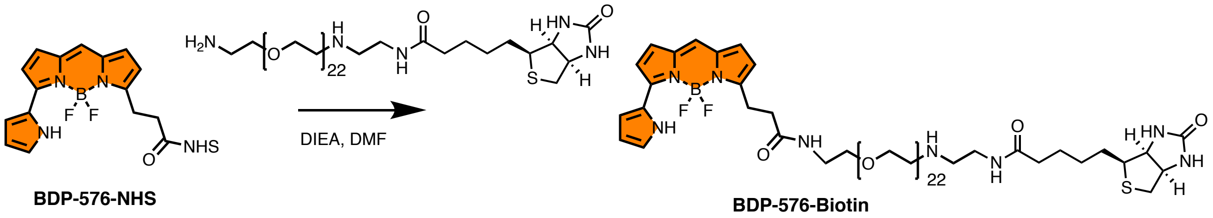


**BDP-576-Biotin**. To a solution of BDP-576-NHS (2.5 mg, 3.9 µmol, 1 eq) in DMF (2 mL) was added (+)-biotin-PEG$_{23}$-CH$_2$CH$_2$NH$_2$ (4.0 mg, 3.8 µmol, 1.2 eq) and DIEA (6 µL, 33 µmol, 10 eq). The solution was left to stir for 3 hours. The crude was concentrated under reduced pressure and purified by column chromatography on silica gel (DCM/MeOH: 9/1) to obtain **BDP-650-Biotin** (5 mg, 93%).

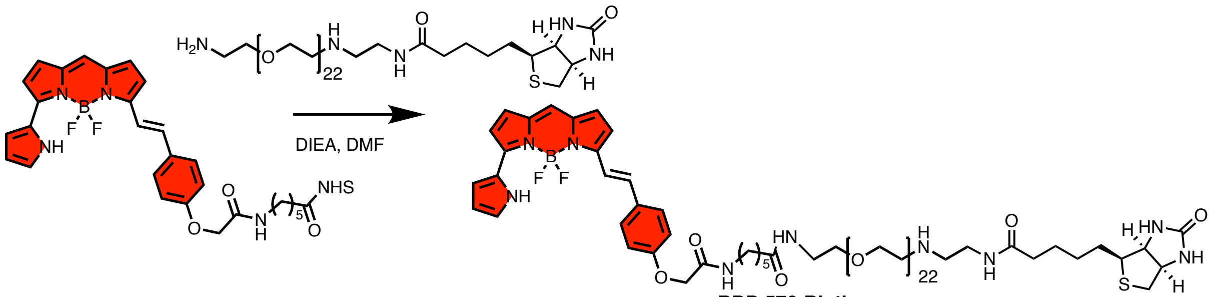


**BDP-650-Biotin**. To a solution of BDP-650-NHS (2.5 mg, 3.9 µmol, 1 eq) in DMF (2 mL) was added (+)-Biotin-PEG$_{23}$-$CH_2CH_2NH_2$ (4.8 mg, 4.6 µmol, 1.2 eq) and DIEA (7 µL, 39 µmol, 10 eq). The solution was allowed to stir for 3 hours. The crude was concentrated under reduced pressure and purified by column chromatography on silica gel (DCM/MeOH: 9/1) to obtain **BDP-650-Biotin** (5 mg, 80%).

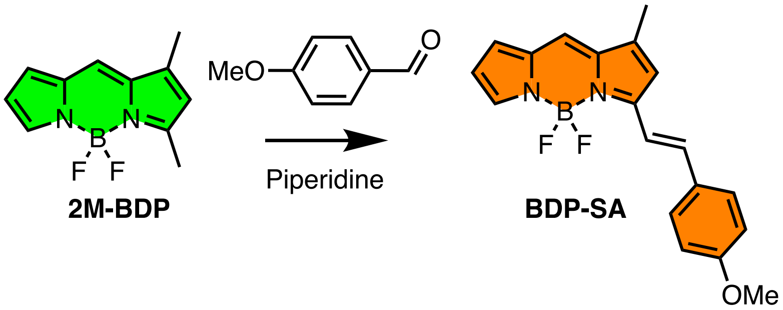


**BDP-SA**. To solution of **2M-BDP**,[2] (180 mg, 0.8 mmol, 1 eq) in piperidine (1 mL, 0.8 mmol, 1 eq) was added 4-Methoxybenzaldehyde (134 mg, 982 mmol, 1.2 eq). The solution was allowed to stir at RT for 10 minutes. The solution was concentrated under reduced pressure and purified by column chromatography on silica gel (Heptan/DCM/acetone : 90/5/5) to give **BDP-SA** (18 mg, 7%). $^1$H NMR (400 MHz, $CDCl_3$) δ 7.65 (s, 1H), 7.59 – 7.54 (m, 2H), 7.50 (dt, J = 16.3, 1.6 Hz, 1H), 7.33 (d, J = 16.2 Hz, 1H), 7.11 (d, J = 0.8 Hz, 1H), 6.95 – 6.90 (m, 2H), 6.90 – 6.86 (m, 1H), 6.72 (d, J = 1.3 Hz, 1H), 6.44 (dd, J = 3.9, 2.1 Hz, 1H), 3.85 (s, 3H), 2.28 (d, J = 1.0 Hz, 3H). $^{13}$C NMR (101 MHz, $CDCl_3$) δ 161.31, 159.52, 144.59, 140.17, 138.05, 138.03, 132.95, 129.71, 128.67, 125.21, 122.34, 117.22, 116.36, 116.05, 114.45, 55.44, 11.47. HRMS (ESI+) calculated for $C_{73}H_{115}BF_2N_{13}O_9S_2$ $[M+H]^+$ 319.1418, found 319.1418.

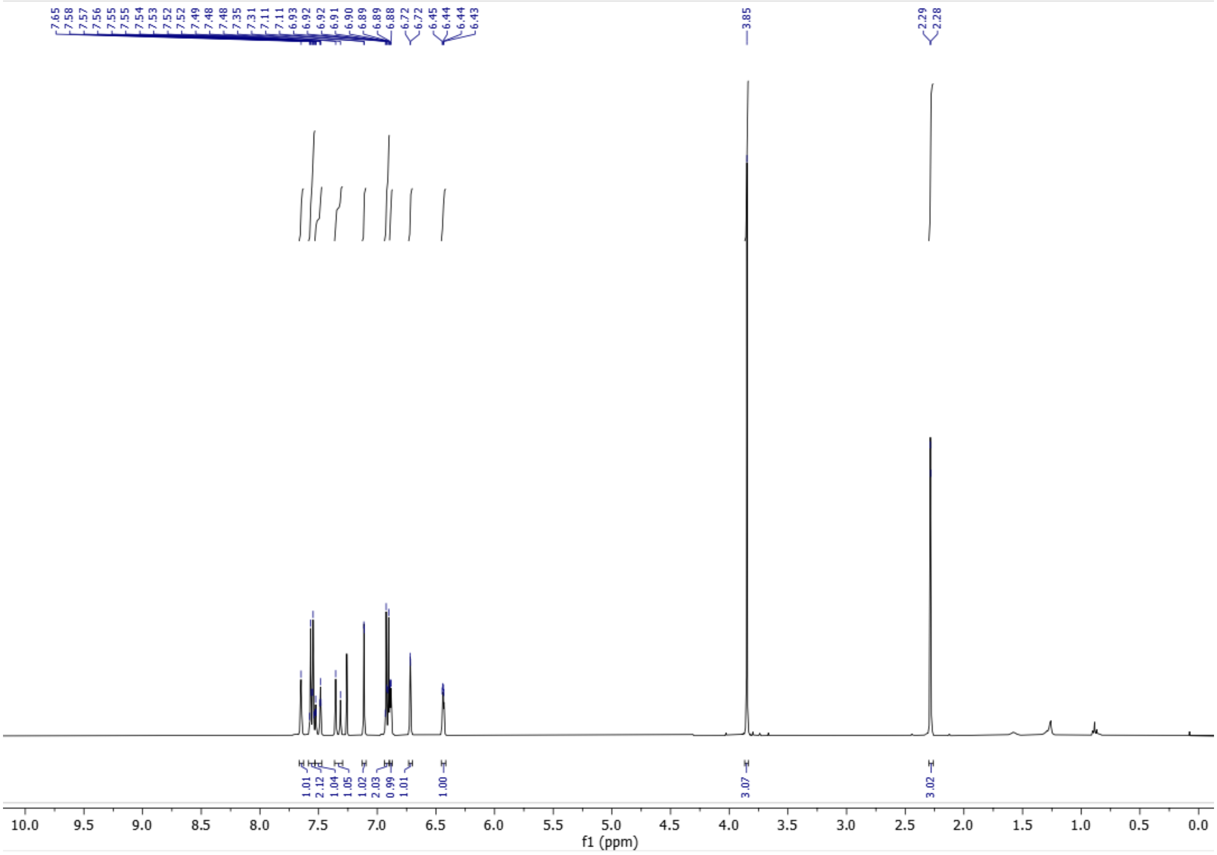


$^{1}H$ NMR spectrum of **BDP-SA**

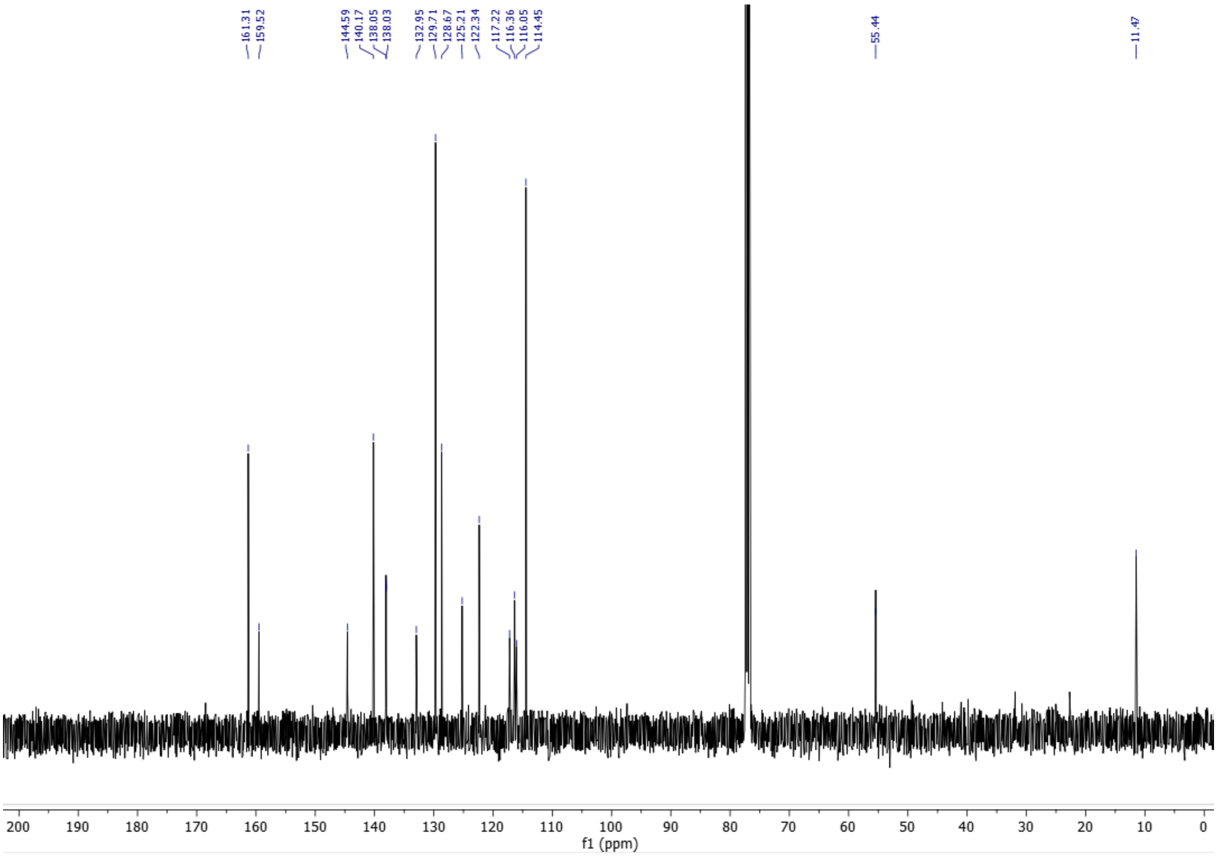


$^{13}C$ NMR spectrum of **BDP-SA**

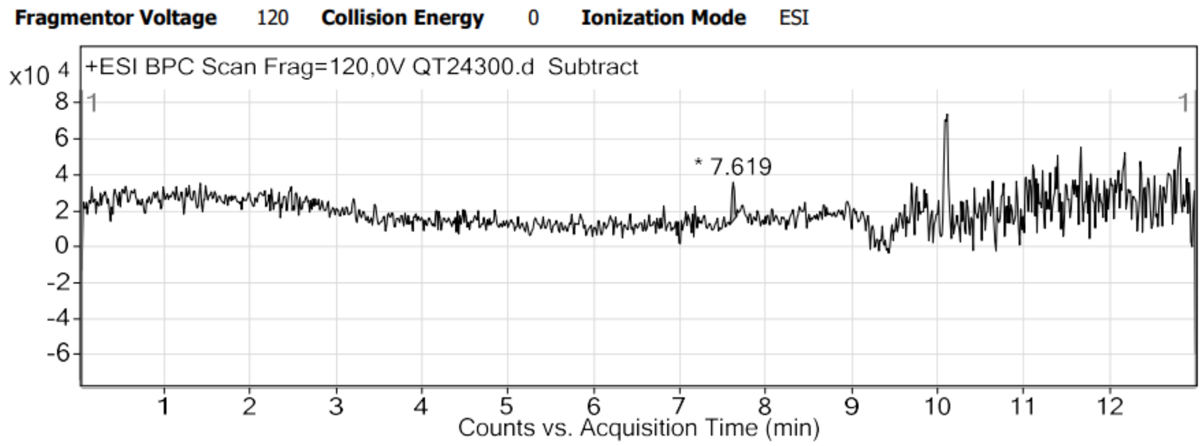


HPLC trace of **BDP-SA**

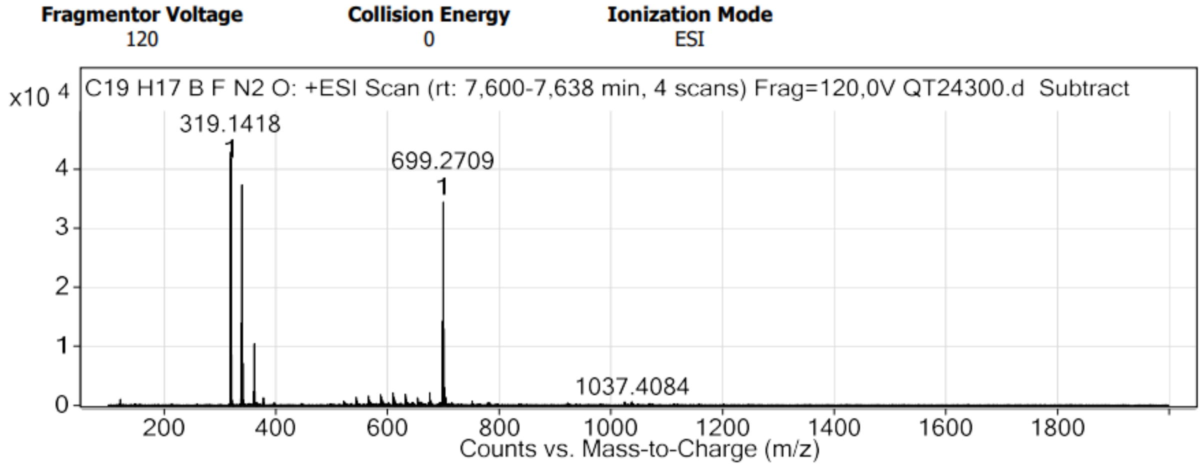


HRMS spectrum of **BDP-SA**

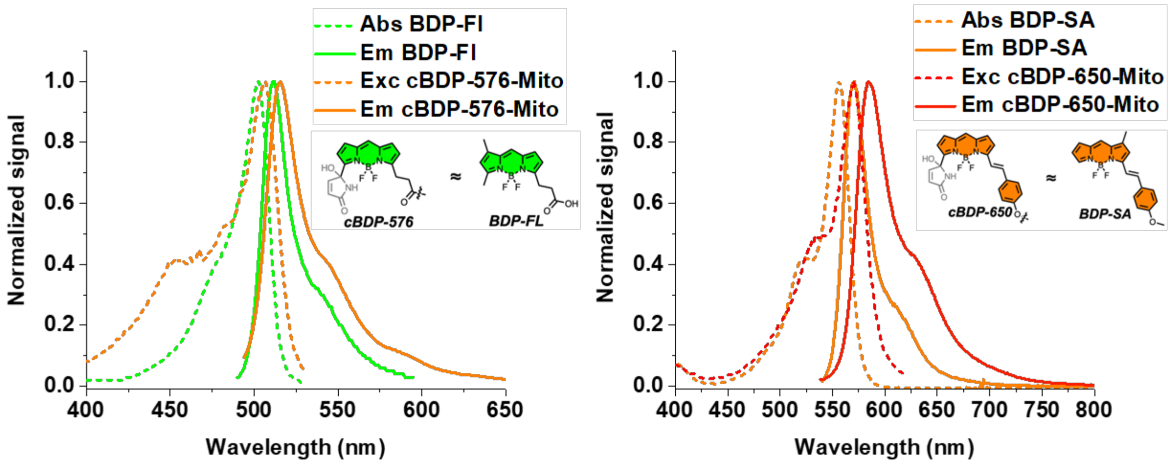


| Dye | $\lambda_{abs}$ (nm) | ε ($M^{-1}.cm^{-1}$) | $\lambda_{em}$ (nm) | $\phi_f$ (%) |
|---|---|---|---|---|
| **cBDP-576** | 506 | N/A | 515 | N/A |
| **BDP-FL** | 502 | 82,000 | 511 | 90 |
| **cBDP-650** | 570 | N/A | 584 | N/A |
| **BDP-SA** | 556 | 97,000 | 571 | 79 |

**Figure S1. Absorption and emission of model of converted form.** The spectra of converted forms were obtained after the conversion step at 1µM in MeOH. Excitation spectra of **cBDP-576** was monitored at $\lambda_{em}$ = 535 nm and **cBDP-650** at $\lambda_{em}$ = 630 nm. Emission spectra were obtained for **cBDP-576** with $\lambda_{ex}$ = 488 nm, **cBDP-650** with $\lambda_{ex}$ = 532 nm and **BDP-SA** with $\lambda_{ex}$ = 530 nm. The spectra of **BDP-FL** were obtained using Spectraviewer tool (ThermoFisher Scientific). These results show that **cBDP-576** and **BDP-Fl** present $\Delta\lambda_{abs}$ = 3 nm and $\Delta\lambda_{em}$ = 4 nm which make **BDP-FL** a very good model of the converted form of **BDP-576** according its photo-physical properties and structure. **cBDP-650** and **BDP-SA** present $\Delta\lambda_{abs}$ = 14 nm and $\Delta\lambda_{em}$ = 13 nm, which make **BDP-SA** a pretty good model of the converted form of **BDP-650** according to photo-physical properties and structure.

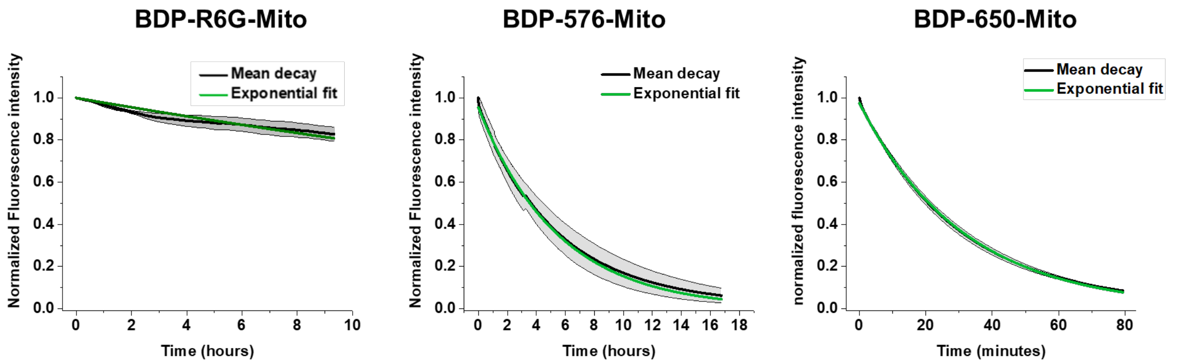


**Figure S2.** Mean Decay of BDPs upon laser irradiation with the exponential fits.**BDP-R6G-Mito** and **BDP-576-Mito** were irradiated in MeOH (1 µM) with 532 nm laser (61 mW.cm$^{-2}$). **BDP-650-Mito** was irradiated in MeOH (1 µM) with 638 nm laser (142 mW.cm$^{-2}$). The graphs present the mean and standard deviation of three independents measurements as well as the exponential fit.

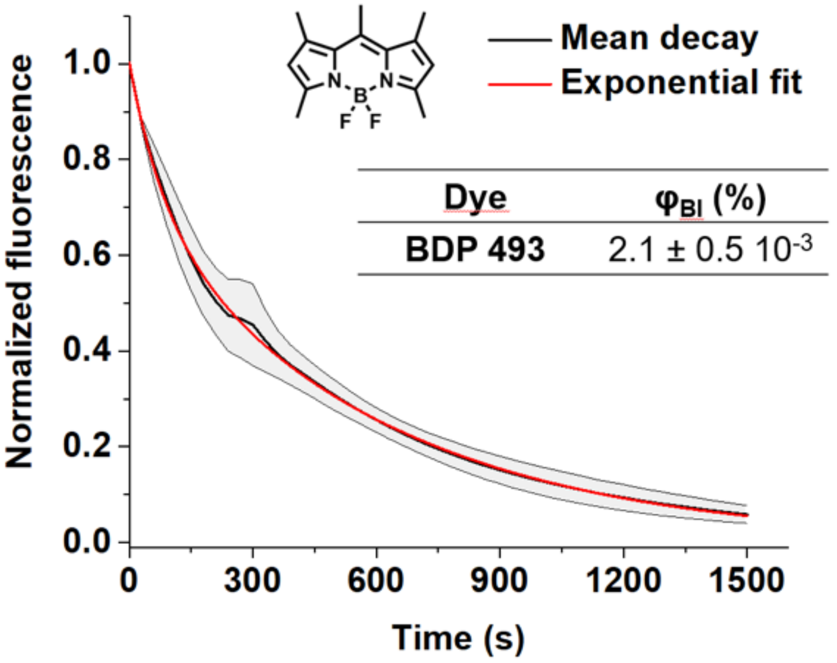


**Figure S3.** Fluorescence decay of BDP-493 upon irradiation with a 488 nm laser providing the quantum yield of photobleaching ($\phi_{Bl}$), depicting the photostability.

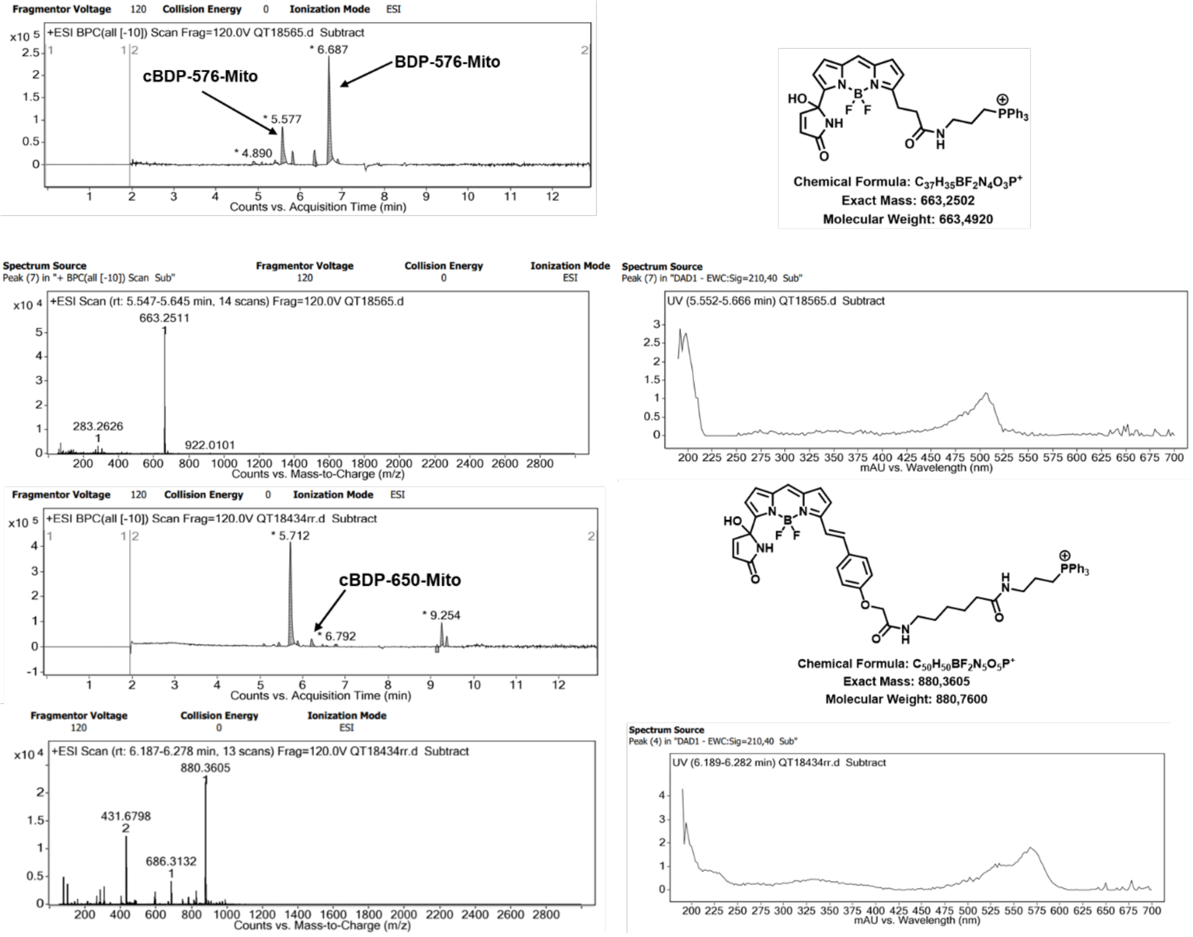


**Figure S4. HPLC-mass analysis of the photoproducts for BDP-576-Mito and BDP-650-Mito.** The results showed the presence of **cBDP-576-Mito and cBDP-650-Mito** expected mass in accordance with their absorbance spectrum.

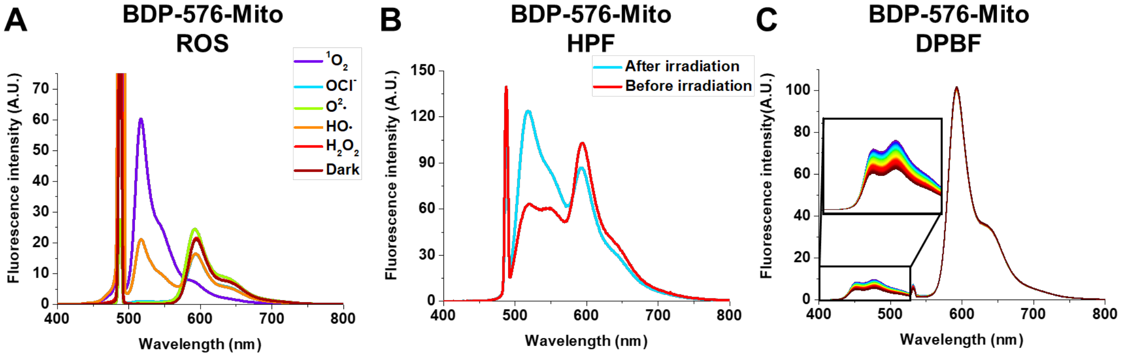


**Figure S5. ROS analysis of BDP-576-Mito**. **A)** Emission spectra of **BDP-576-Mito** in the presence of various ROS, without ROS in the dark (negative control). Selectivity tests were performed adding various ROS generators in water in a methanolic solution of **BDP-576-Mito** (1 µM). $^1O_2$: 5 µM of Aluminium Phthalocyanine with 638 nm laser for 30 minutes: $O_2^{\bullet -}$: 2 mM of $KO_2$ for 30 minutes in the dark; $OCl^-$ : 2 mM of NaOCl for 30 minutes in the dark; $^{\bullet}OH$: 2 mM of $FeSO_4$ + 2 mM $H_2O_2$ for 30 minutes in the dark; $H_2O_2$: 2 mM of $H_2O_2$ for 30 minutes in the dark. The emission spectra were acquired with 488 nm laser. **B)** Emission spectra of **BDP-576-Mito** (1µM) and HPF (20 µM) before and after 30 minutes irradiation at 532 nm. C)

Emission spectra of **BDP-576-Mito** (1 µM) and DPBF (100 µM) during irradiation at 532 nm and 405 nm.

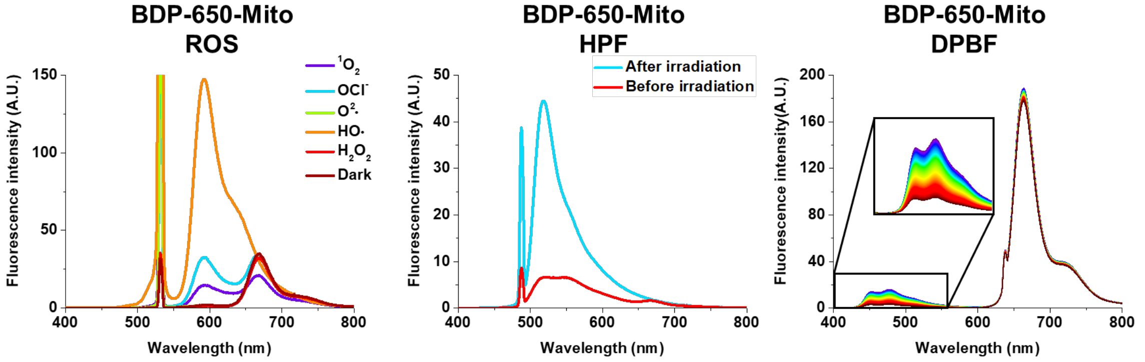


**Figure S6. ROS analysis of BDP-650-Mito**. A) Emission spectra of **BDP-650-Mito** in the presence of various ROS, without ROS in the dark (negative control). Selectivity tests were performed adding various ROS generators in water in a methanolic solution of **BDP-650-Mito** (1 µM). $^1O_2$: 5 µM of IR800 with 730 nm laser for 30 minutes: $O_2^{\bullet-}$ : 2 mM of $KO_2$ for 30 minutes in the dark; OCl- : 2 mM of NaOCl for 30 minutes in the dark; $^{\bullet}OH$: 2 mM of $FeSO_4$ + 2 mM $H_2O_2$ for 30 minutes in the dark; $H_2O_2$: 2 mM of $H_2O_2$ for 30 minutes in the dark. The emission spectra were acquired with 532 nm laser. B) Emission spectra (ex 488 nm) of **BDP-650-Mito** (1µM) and HPF (20 µM) before and after 30 minutes irradiation at 638 nm. C) Emission spectra of **BDP-576-Mito** (1 µM) and DPBF (100 µM) during irradiation at 638 nm and 405 nm.

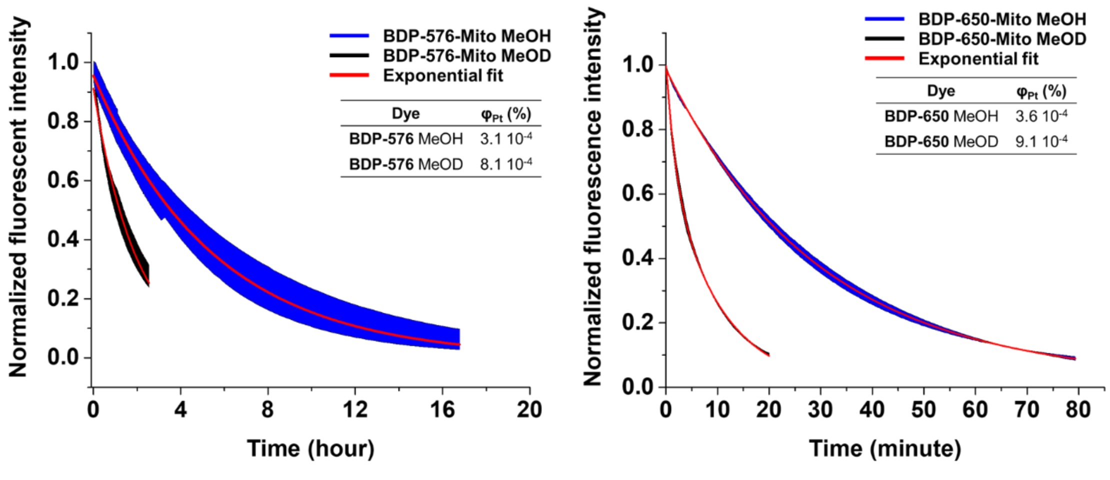


**Figure S7.** Photoconversion of **BDP-576-Mito** and **BDP-650-Mito** in MeOH and MeOD showing the influence of singlet oxygen in the conversion process.

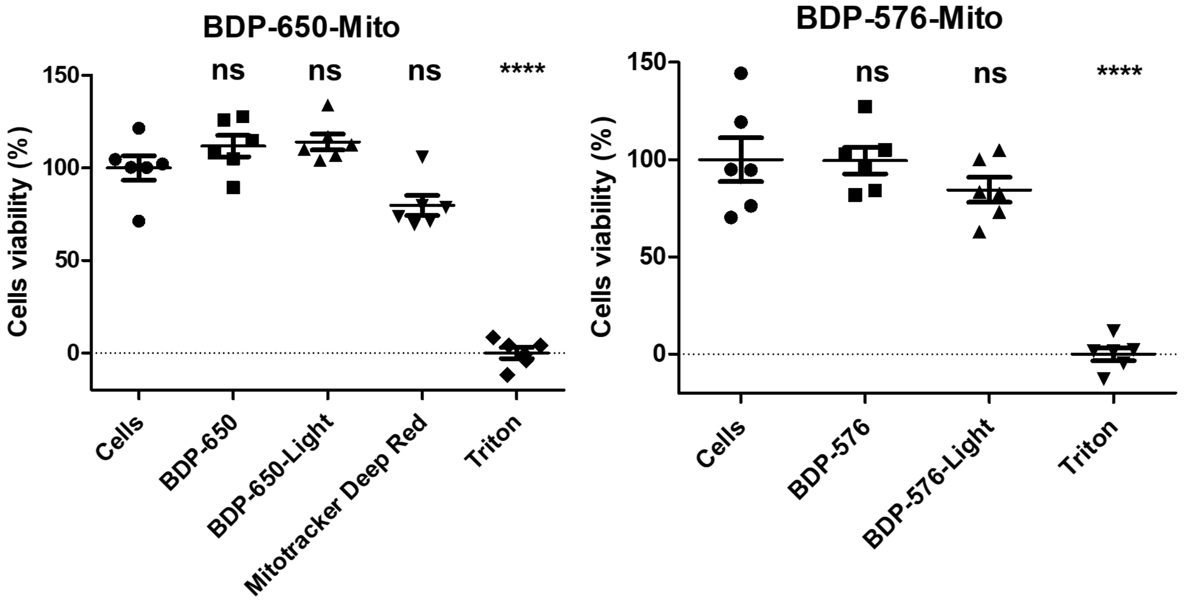


**Figure S8. Cytotoxicity and photocytotoxicity assays for BDP-576-Mito and BDP-650-Mito.** The results showed that both probes where not cytotoxic nor phototoxic upon photoconversion. The MitoTracker™ Deep Red probe was irradiated in the same conditions than for BDP-650-Mito.

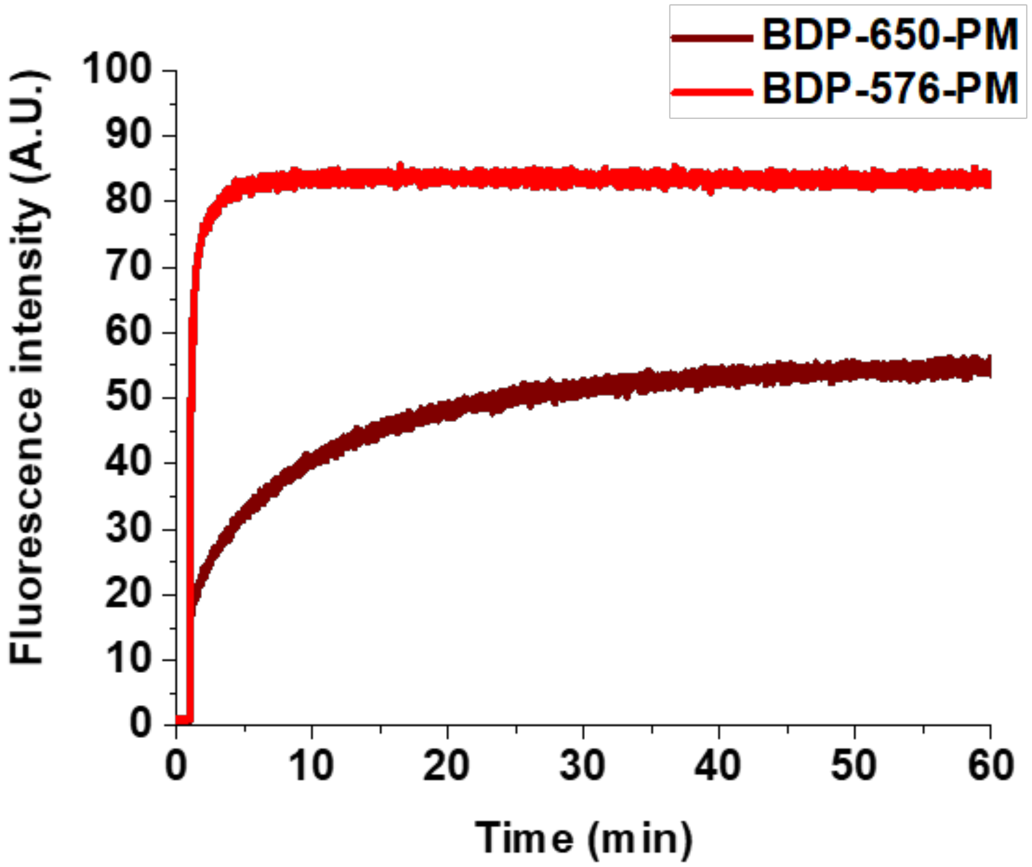


**Figure S9**. Kinetic of binding to DOPC Large Unilamellar Vesicles (LUVs) of the plasma membrane probes **BDP-576-PM** and **BDP-650-PM.** The probes (1 µM) were added after 1 min to a solution of Large Unilamelar vesicles LUVS (200 µM DOPC in PBS, size 100 nm). The fluorescence intensity was monitored at 660 nm for BDP-650-PM ($\lambda_{Ex}$= 640 nm) and 580 nm for BDP-576-PM ($\lambda_{Ex}$= 560 nm) over 1h.

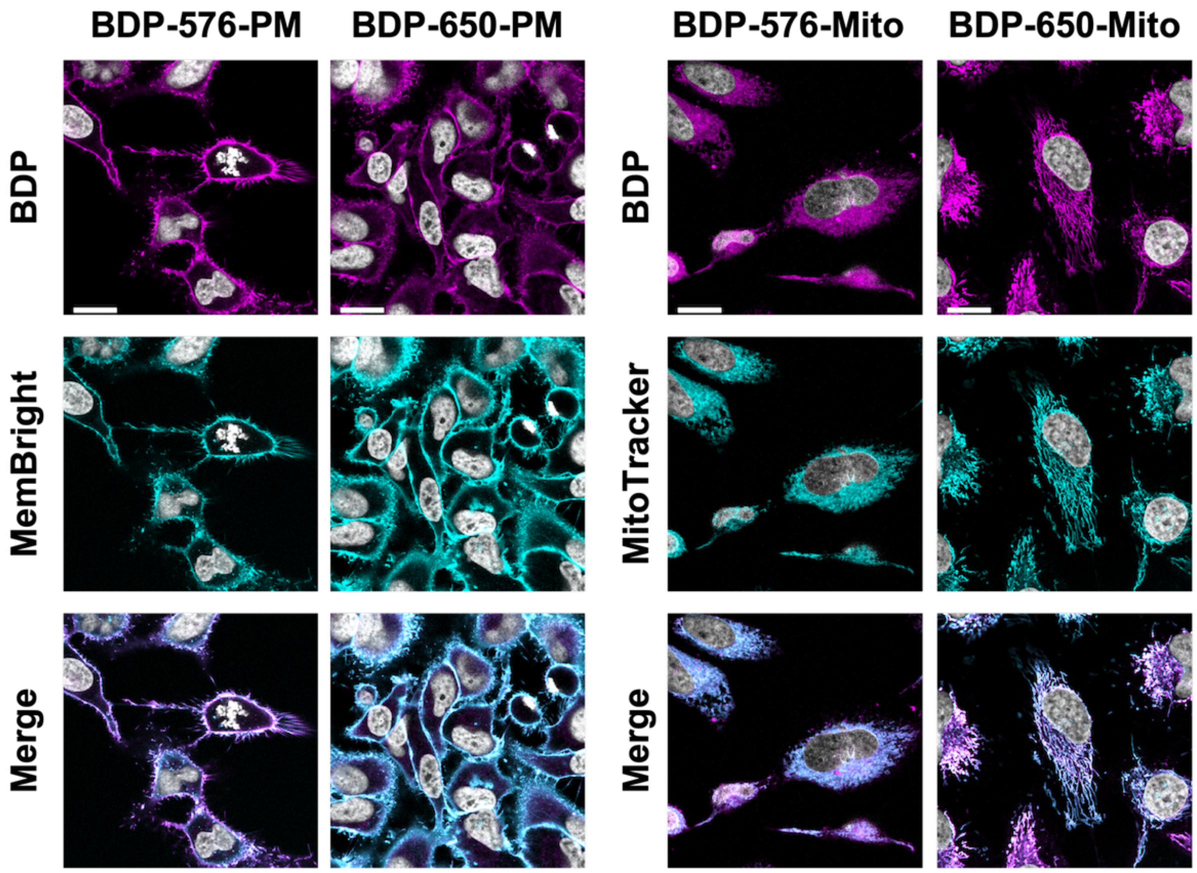


**Figure S10. Colocalization of targeted. pyrrolyl-BDPs**. **BDP-576-PM** (200 nM) was colocalized with MBcy5 (200 nM) (Pearson's coefficient: 0.884). **BDP-650-PM** (200 nM) was colocalized with MemBright 488 (200 nM) (Pearson's coefficient: 0.842). **BDP-576-Mito** (200 nM) was colocalized with MitoTracker Deep Red (200 nM) (Pearson's coefficient: 0.800). **BDP-650-Mito** (200 nM) was colocalized with MitoTracker Green (200 nM) (Pearson's coefficient: 0.873). Scale bar is 20 µm.

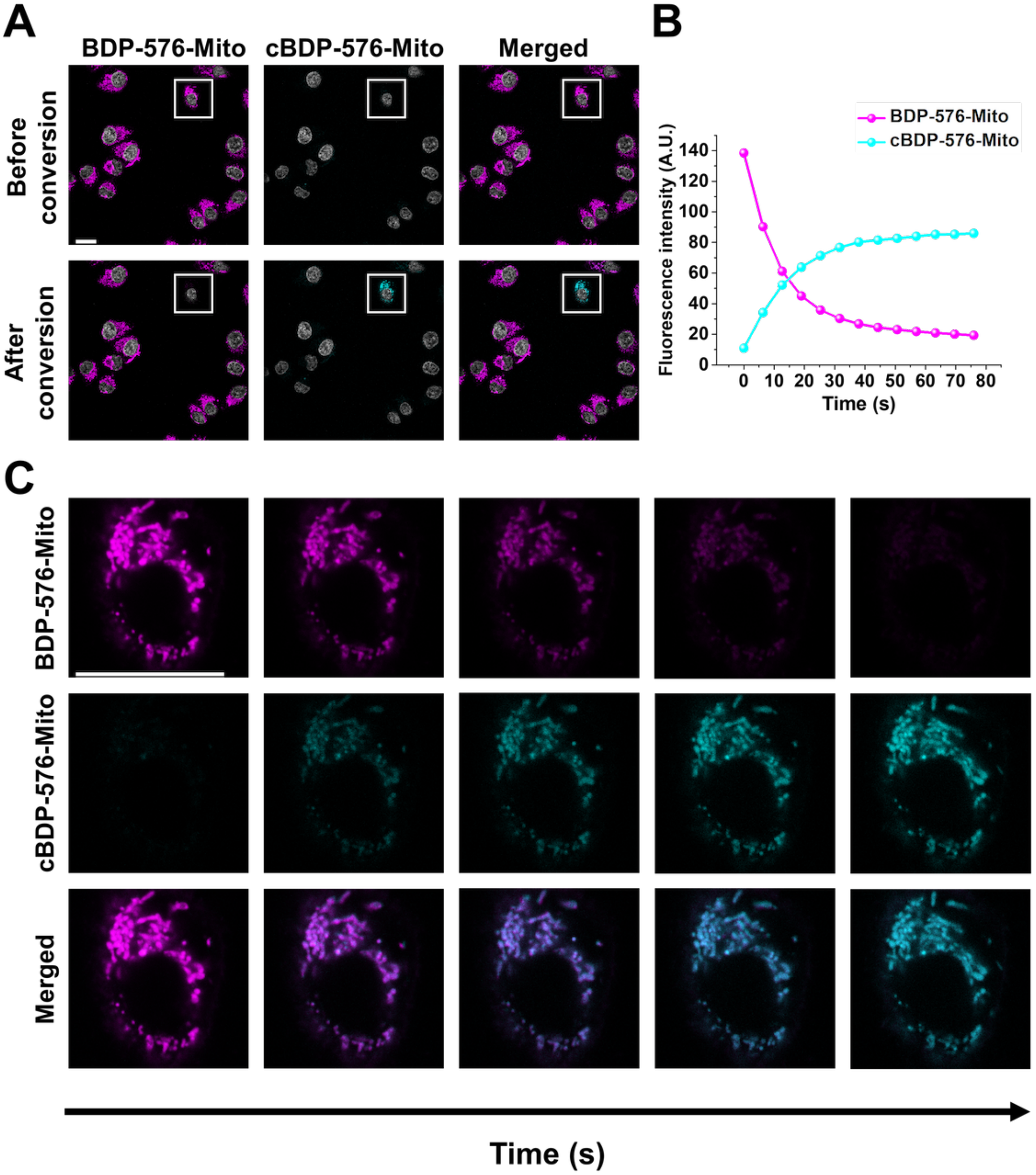


**Figure S11. Photoconversion of BDP-576-Mito**. (A) Laser scanning confocal images of HeLa incubated with **BDP-576-Mito** (200 nM). **BDP-576-Mito** (in magenta) and its converted form **cBDP-576-Mito** (in cyan) were, respectively, excited at 552 nm (fluorescence signal: 560-625 nm) and 488 nm (fluorescence signal: 500-550 nm). Nuclei are stained with Hoechst (in grey). (B) Fluorescence intensity in the ROI over time of these two channels. (C) Images in the ROI over time. Scale bar is 20 µm.

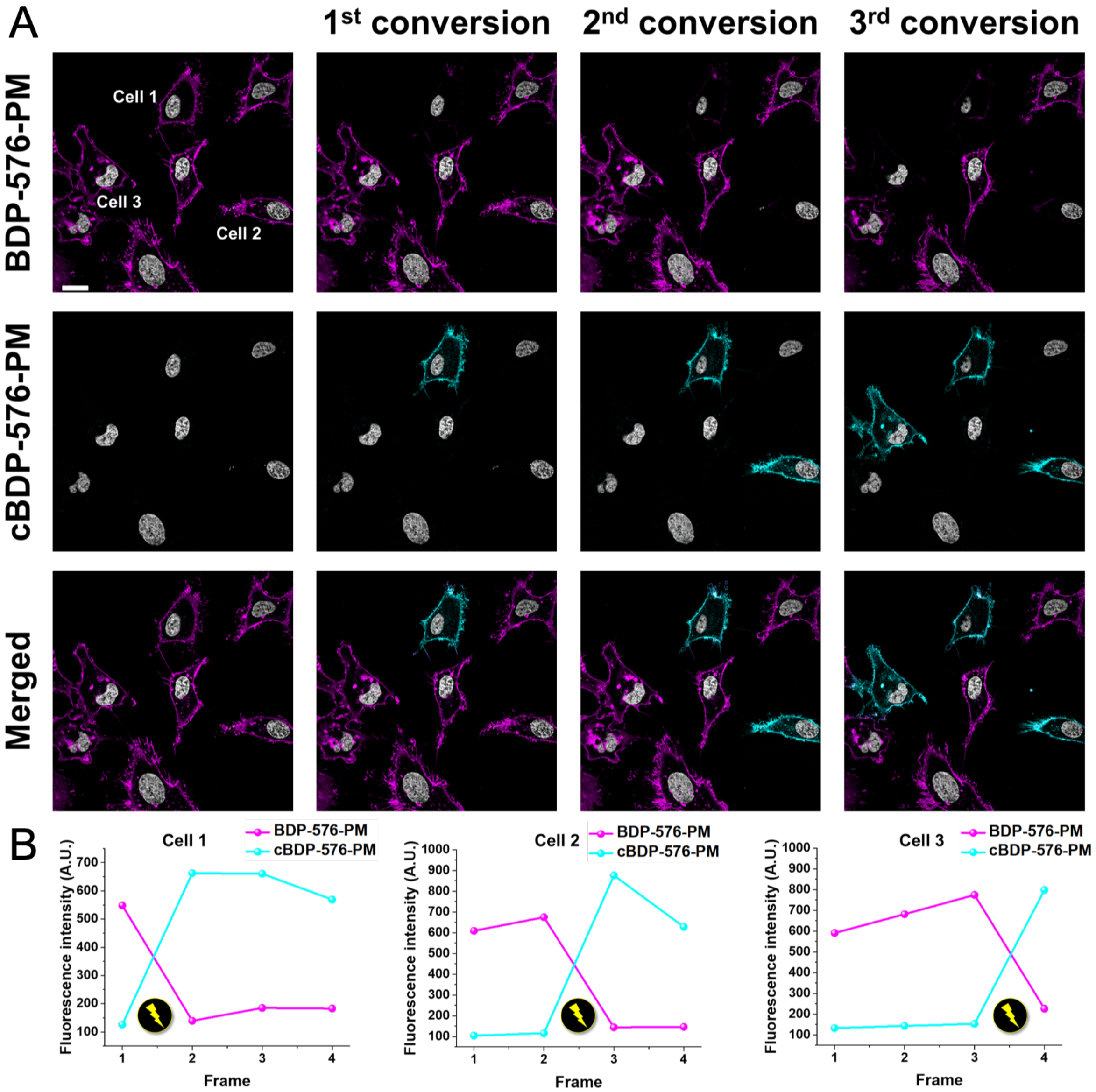


**Figure S12.** Sequential photoconversion of live cells' plasma membrane. (A) Laser scanning confocal microscope images of live HeLa cells stained with BDP-576-PM (200 nM). The cells were sequential irradiated using the 488 nm laser line to convert the BDP-576-PM-stained plasma membranes (magenta) into cBDP-576-PM (cyan) The nucleus (grey) was stained with Hoechst 33258 (5 µg. $mL^{-1}$). Scale bar is 20 µm. (B) Fluorescence signal (mean intensity of individual cells) in the BDP-576 channel (Magenta) and the cBDP-576 one (cyan) over time. Time interval between conversions was 20 s.

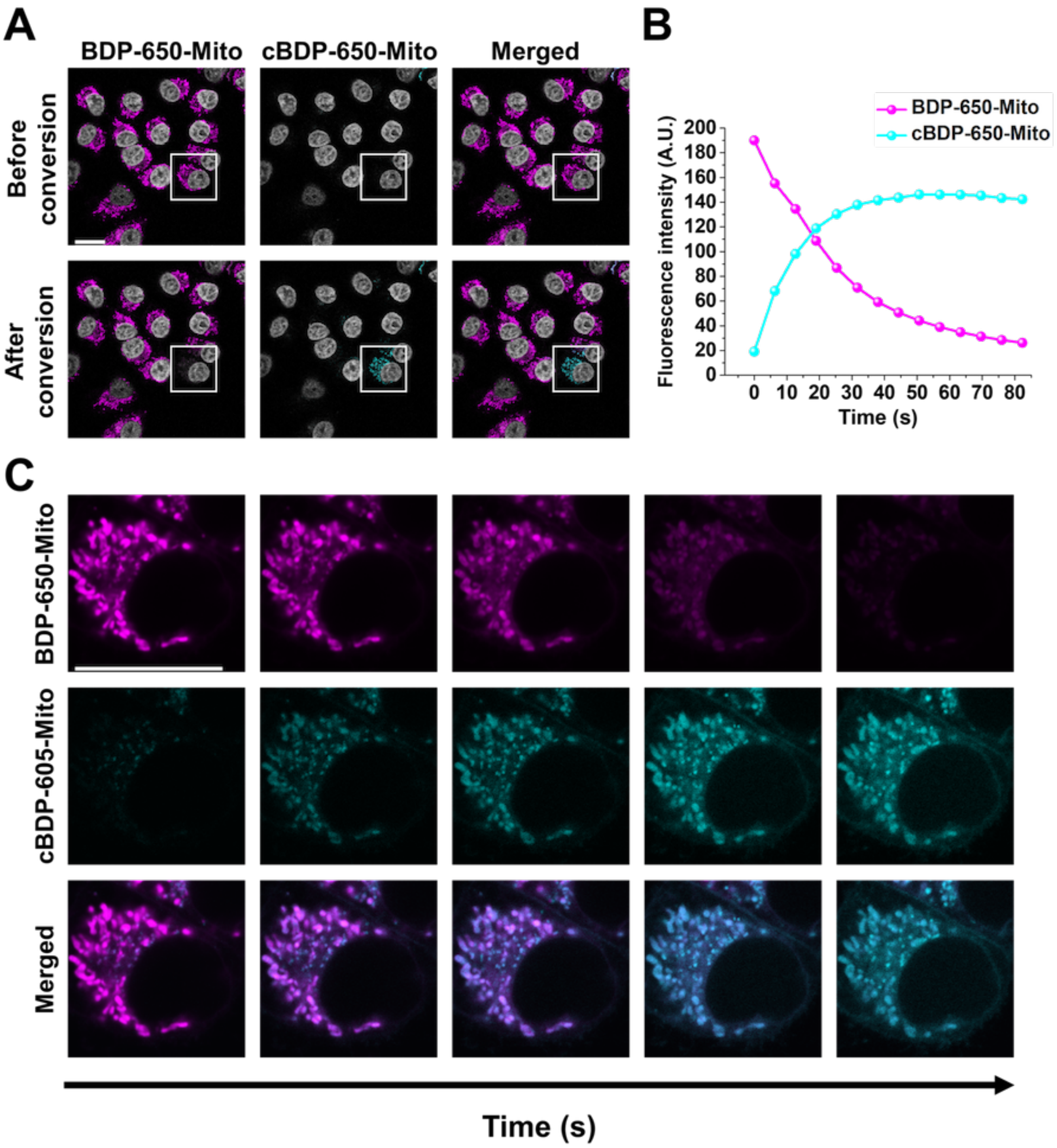


**Figure S13. Photoconversion of BDP-650-Mito**. (A) Laser scanning confocal images of HeLa incubated with **BDP-650-Mito** (200 nM). **BDP-650-Mito** and its converted form **cBDP-650-Mito** were, respectively, excited at 638 nm (fluorescence signal: 660-750 nm) and 552 nm (fluorescence signal: 560-625 nm). (B) Fluorescence intensity in the ROI over time of these two channels. (C) Images in the ROI over time. Scale bar is 20 µm.

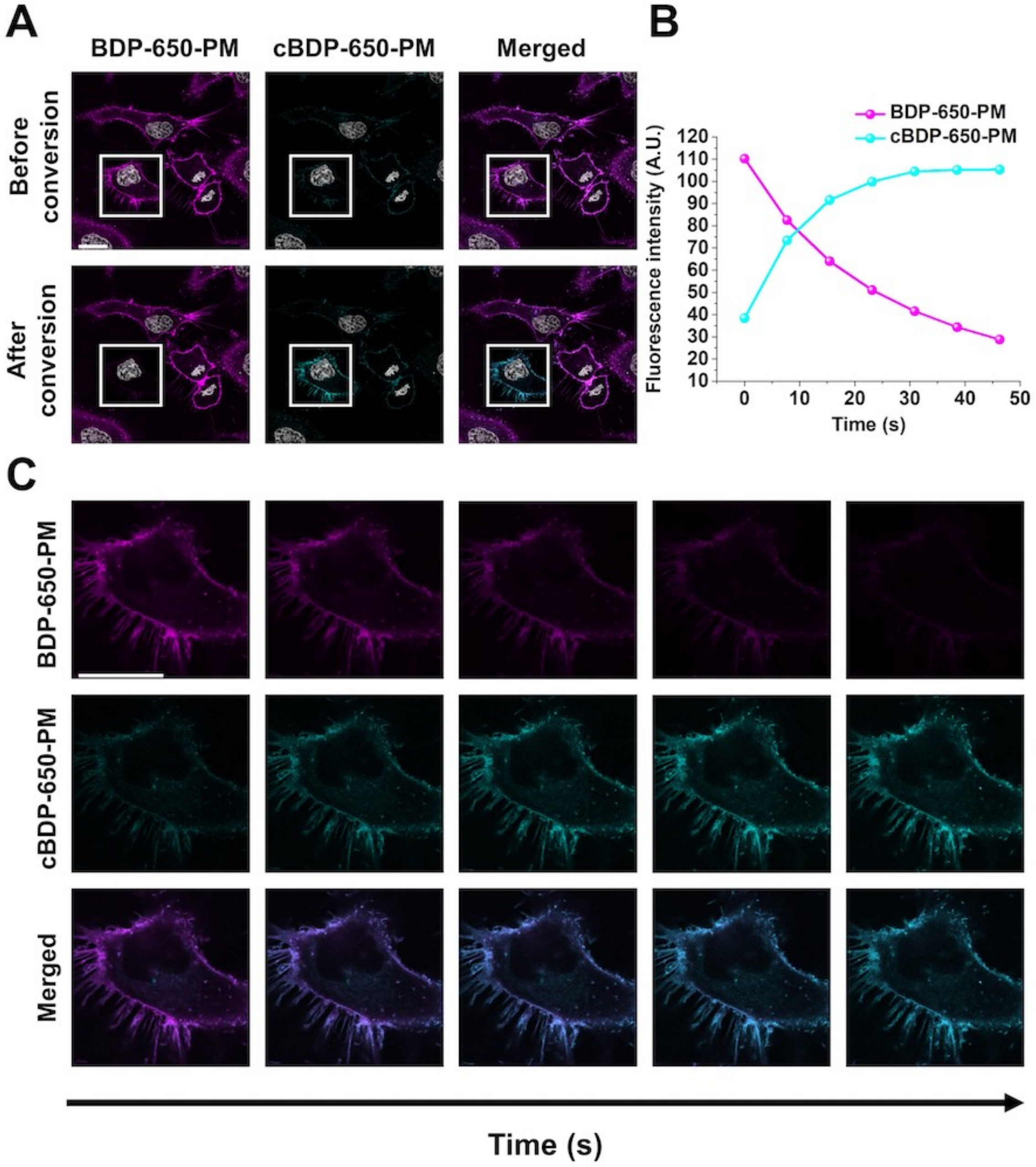


**Figure S14. Photoconversion of BDP-650-PM**. (A) Laser scanning confocal images of HeLa incubated with **BDP-650-PM** (200 nM). **BDP-650-PM** and its converted form **cBDP-650-PM** were, respectively, excited at 638 nm (fluorescence signal: 660-750 nm) and 552 nm (fluorescence signal: 560-625 nm). (B) Fluorescence intensity in the ROI over time of these two channels. (C) Images in the ROI over time. Scale bar is 20 µm.

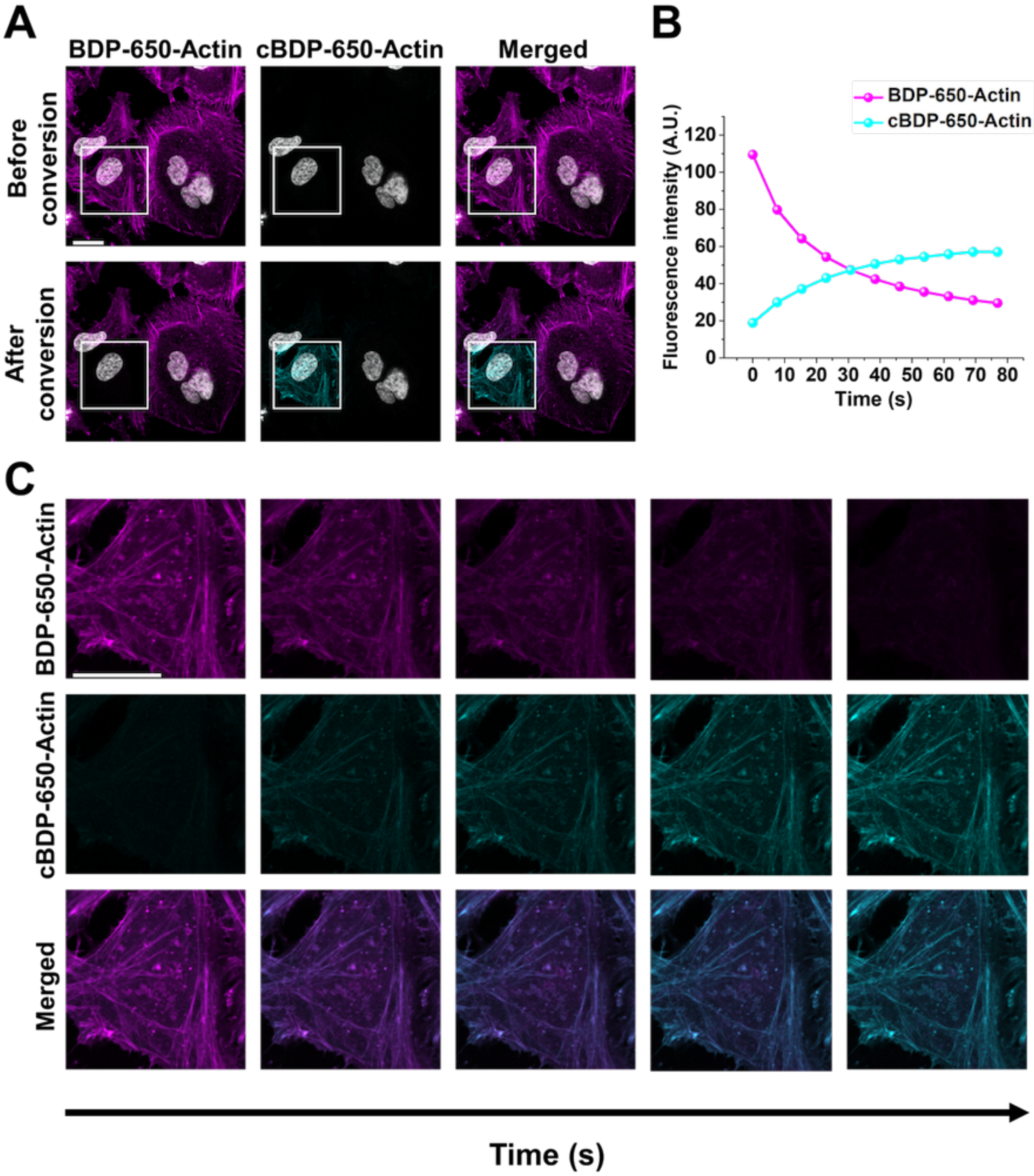


**Figure S15. Photoconversion of BDP-650-Actin**. (A) Laser scanning confocal images of HeLa incubated with **BDP-650-Actin** (200 nM). **BDP-650-Actin** and its converted form **cBDP-650-Actin** were, respectively, excited at 638 nm (fluorescence signal: 660-750 nm) and 552 nm (fluorescence signal: 560-625 nm). (B) Fluorescence intensity in the ROI over time of these two channels. (C) Images in the ROI over time. Scale bar is 20 µm.

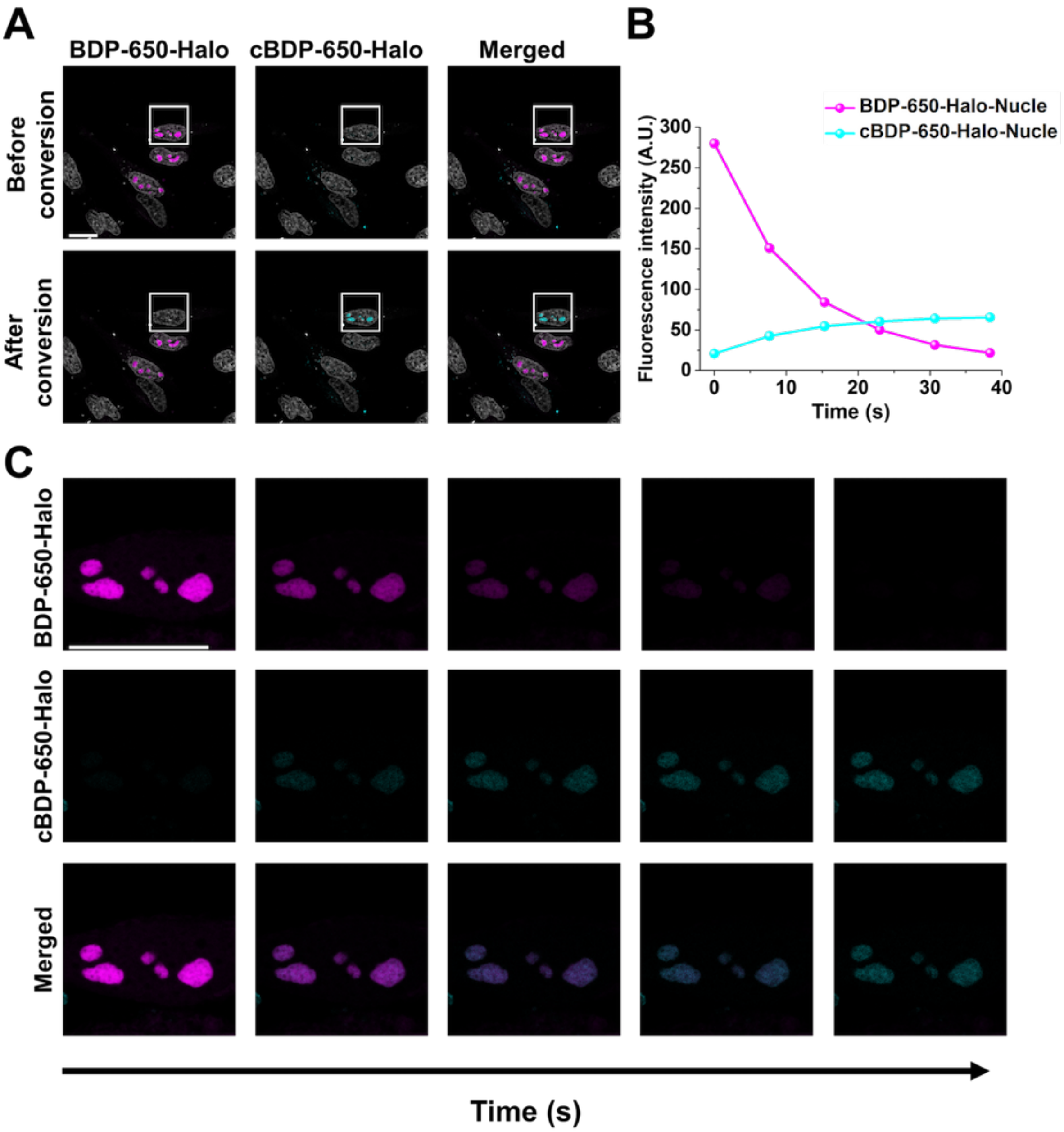


**Figure S16A. Photoconversion of BDP-650-Halo in nucleoli**. (A) Laser scanning confocal images of HeLa incubated with **BDP-650-Halo** (50 nM). **BDP-650-Halo** and its converted form **cBDP-650-Halo** were, respectively, excited at 638 nm (fluorescence signal: 660-750 nm) and 552 nm (fluorescence signal: 560-625 nm). (B) Fluorescence intensity in the ROI over time of these two channels. (C) Images in the ROI over time. Scale bar is 20 µm.

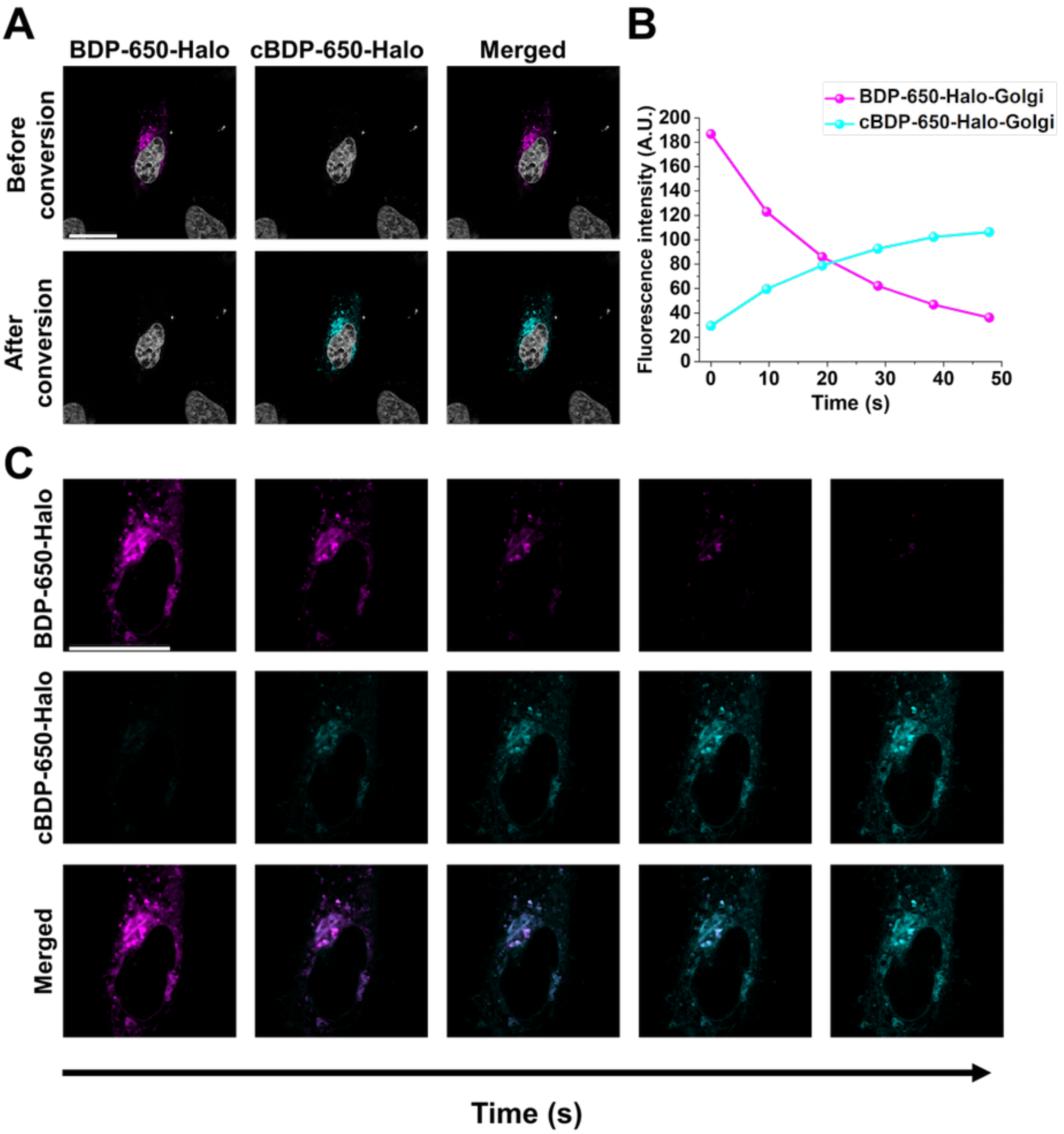


**Figure S16B. Photoconversion of BDP-650-Halo in Golgi apparatus**. (A) Laser scanning confocal images of HeLa incubated with **BDP-650-Halo** (50 nM). **BDP-650-Halo** and its converted form **cBDP-650-Halo** were, respectively, excited at 638 nm (fluorescence signal: 660-750 nm) and 552 nm (fluorescence signal: 560-625 nm). (B) Fluorescence intensity in the ROI over time of these two channels. (C) Images in the ROI over time. Scale bar is 20 μm.

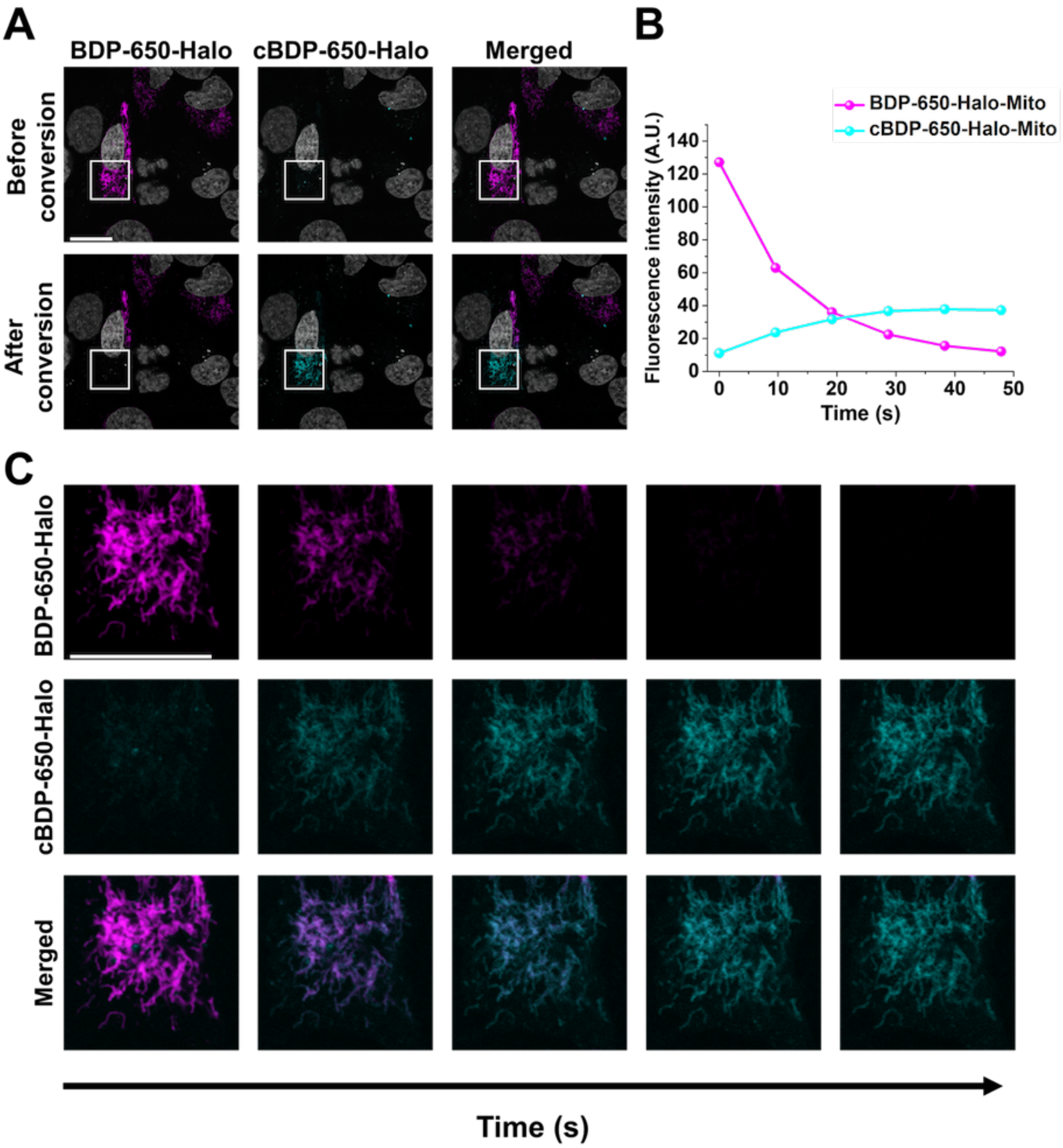


**Figure S16C. Photoconversion of BDP-650-Halo in mitochondria.** (A) Laser scanning confocal images of HeLa incubated with **BDP-650-Halo** (50 nM). **BDP-650-Halo** and its converted form **cBDP-650-Halo** were, respectively, excited at 638 nm (fluorescence signal: 660-750 nm) and 552 nm (fluorescence signal: 560-625 nm). (B) Fluorescence intensity in the ROI over time of these two channels. (C) Images in the ROI over time. Scale bar is 20 µm.

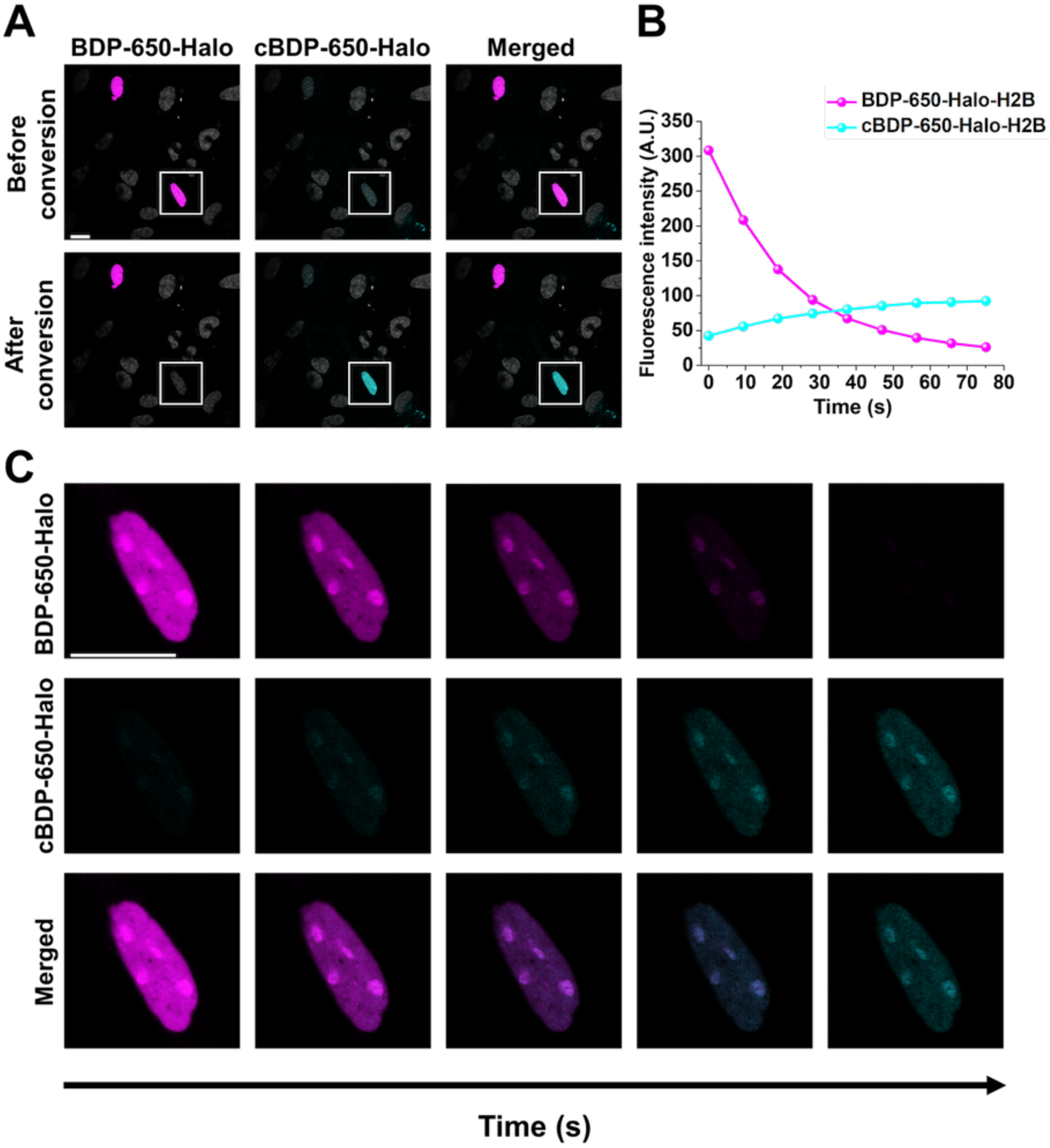


**Figure S16D. Photoconversion of BDP-650-Halo in nucleus**. (A) Laser scanning confocal images of HeLa incubated with **BDP-650-Halo** (50 nM). **BDP-650-Halo** and its converted form **cBDP-650-Halo** were, respectively, excited at 638 nm (fluorescence signal: 660-750 nm) and 552 nm (fluorescence signal: 560-625 nm). (B) Fluorescence intensity in the ROI over time of these two channels. (C) Images in the ROI over time. Scale bar is 20 µm.

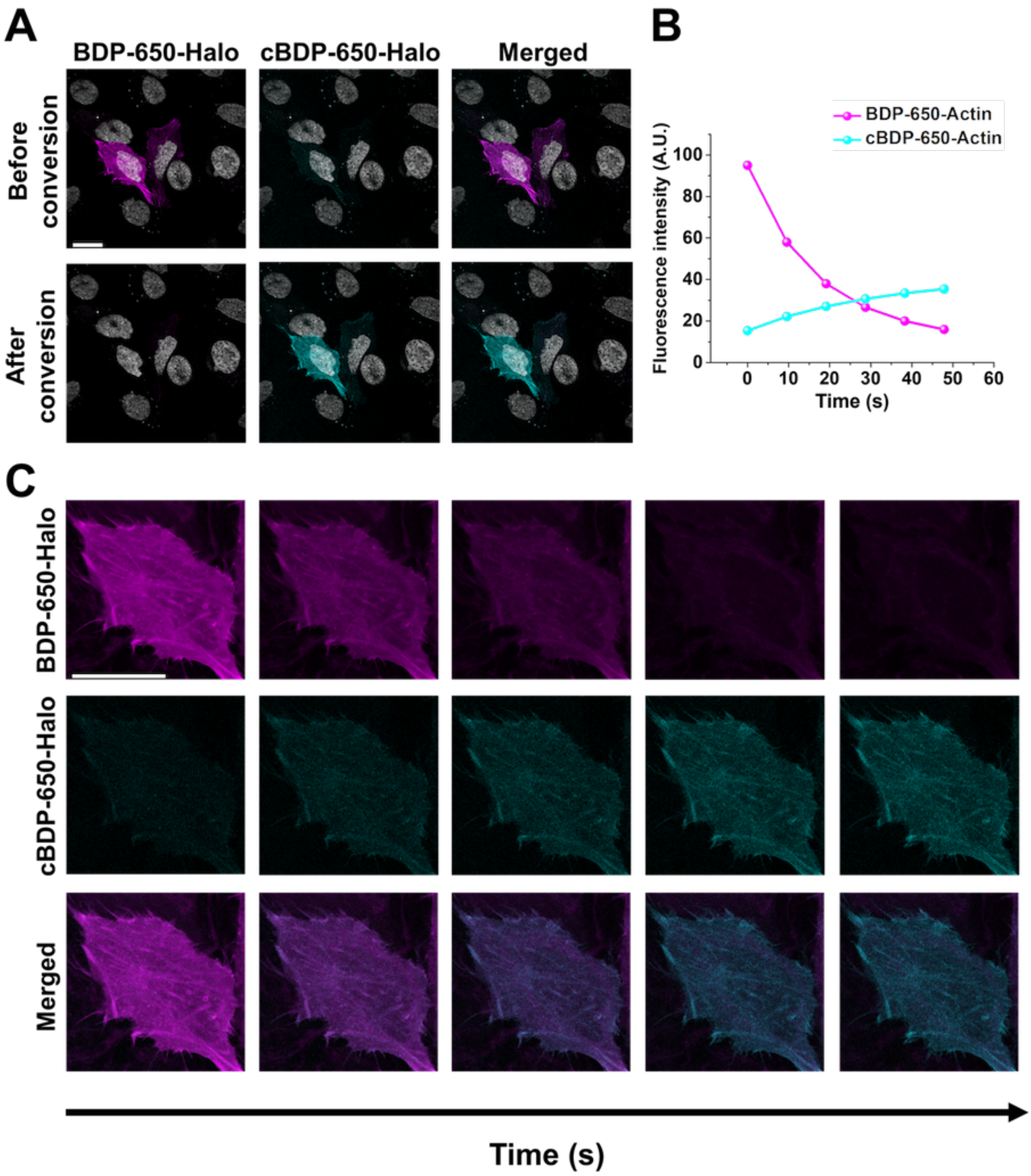


**Figure S16E. Photoconversion of BDP-650-Halo in actin filaments**. (A) Laser scanning confocal images of HeLa incubated with **BDP-650-Halo** (50 nM). **BDP-650-Halo** and its converted form **cBDP-650-Halo** were, respectively, excited at 638 nm (fluorescence signal: 660-750 nm) and 552 nm (fluorescence signal: 560-625 nm). (B) Fluorescence intensity in the ROI over time of these two channels. (C) Images in the ROI over time. Scale bar is 20 μm.

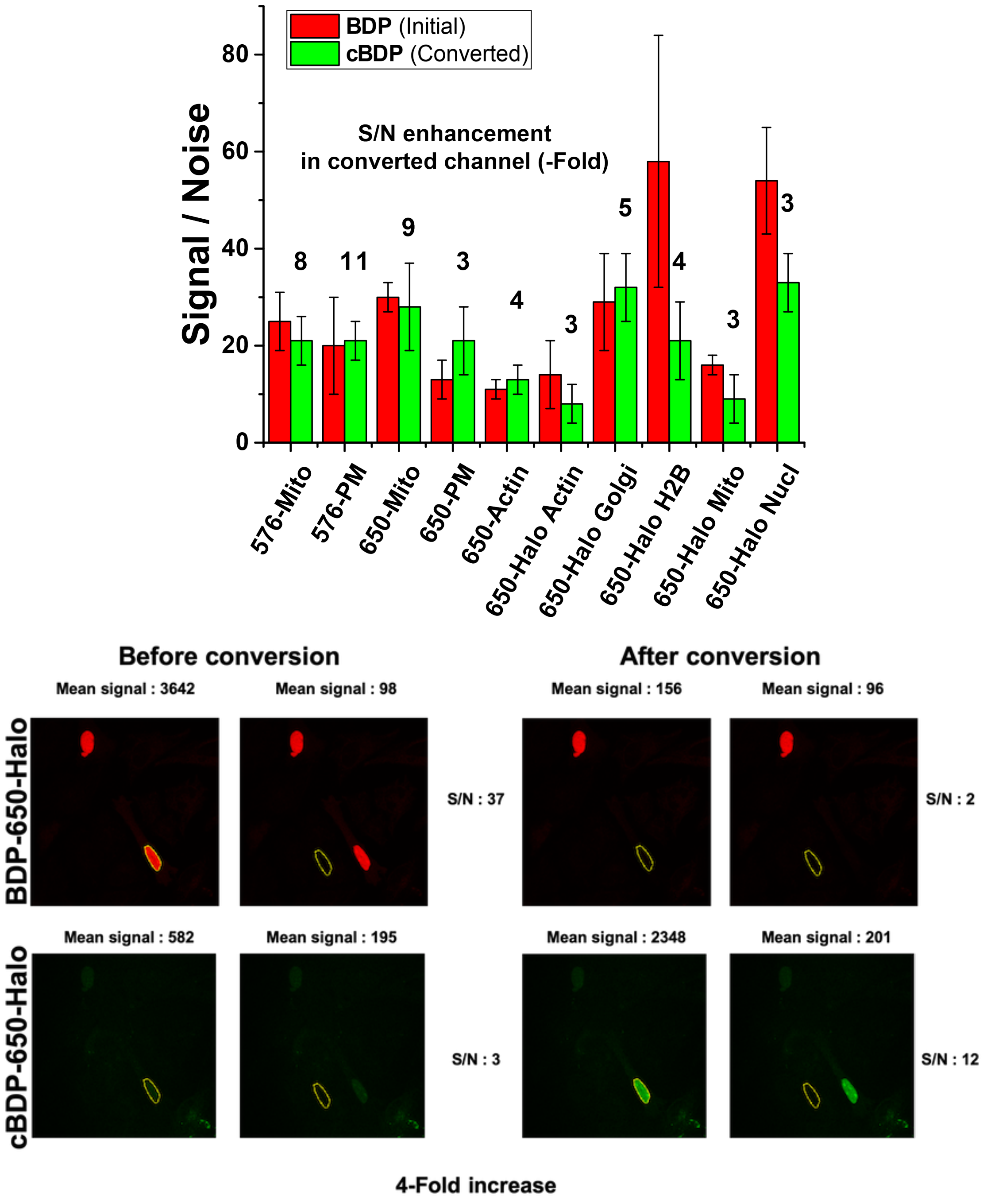


**Figure S17. Images quality before and after photoconversion.** (top) Signal to noise ratio obtained in laser scanning confocal imaging with pyrrolyl-BODIPYs (BDP) and their converted forms (cBDP). Labels on top of bars are the signal to noise ratio enhancement of the converted form after conversion. (Bottom) Example (BDP-650-Halo in nucleus) of the protocol used to determine the S/N ratios. The signal was monitored in the ROI (Yellow) were the probe is localized on the non-converted channel. The ROI was then moved to a dark area to obtain the background signal. The signal of the same ROIs was then monitored after the conversion step.

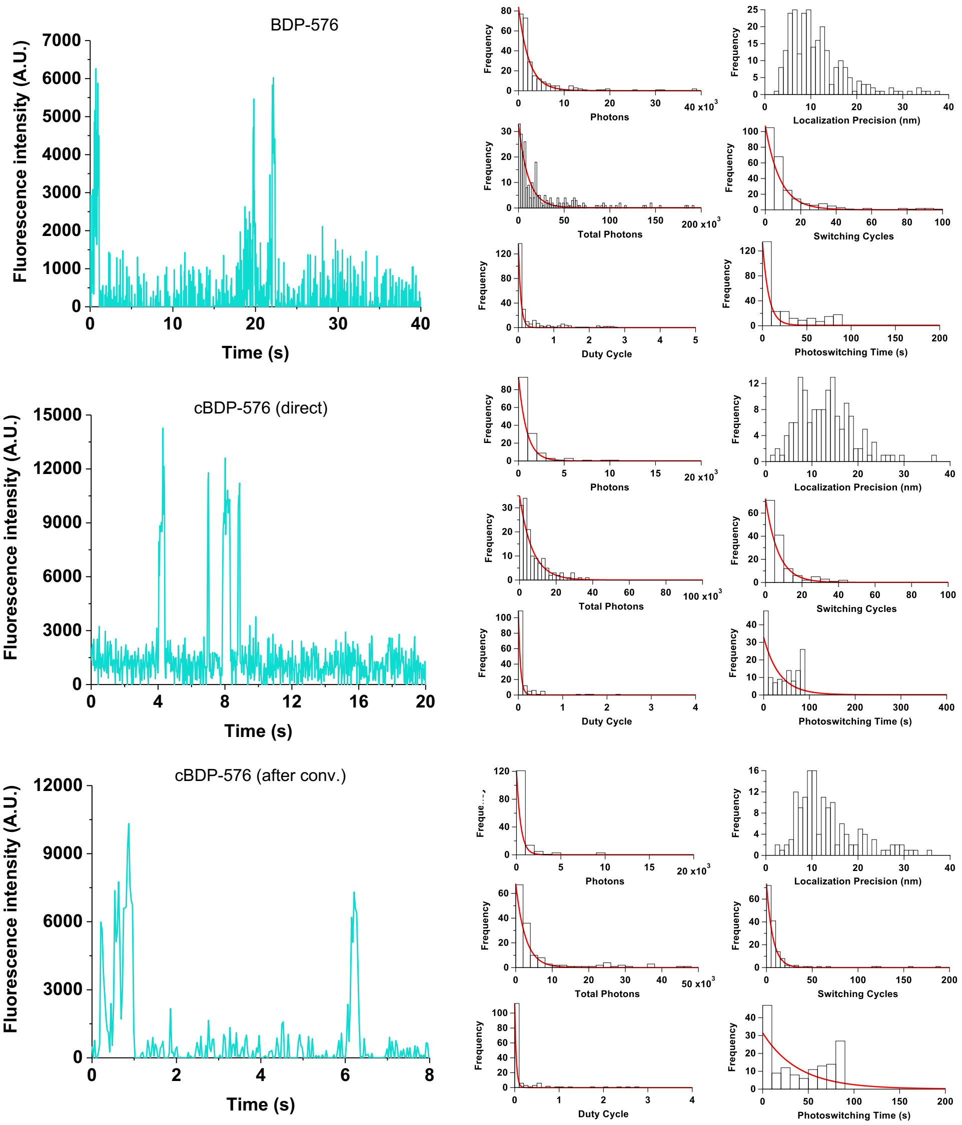


**Figure S18A. Single molecules and photoswitching properties analysis of BDP-576-Biotin.** "Photons" defines the number of photons detected by blinking event, it was calculated through the signal obtained knowing the setting of the camera (Gain, sensitivity). The "Total photons" defined the number of photons detected until the dye bleached. "Duty cycle" is the ratio between the time spent at the ON state and the time spent at the OFF state. "Localization precision" is the uncertainty of the localization, it is given by the thunderstorm plugin. "Switching cycle" is the number of ON/OFF cycles. The "photoswitching time" is the time during the dye switches before it bleaches.

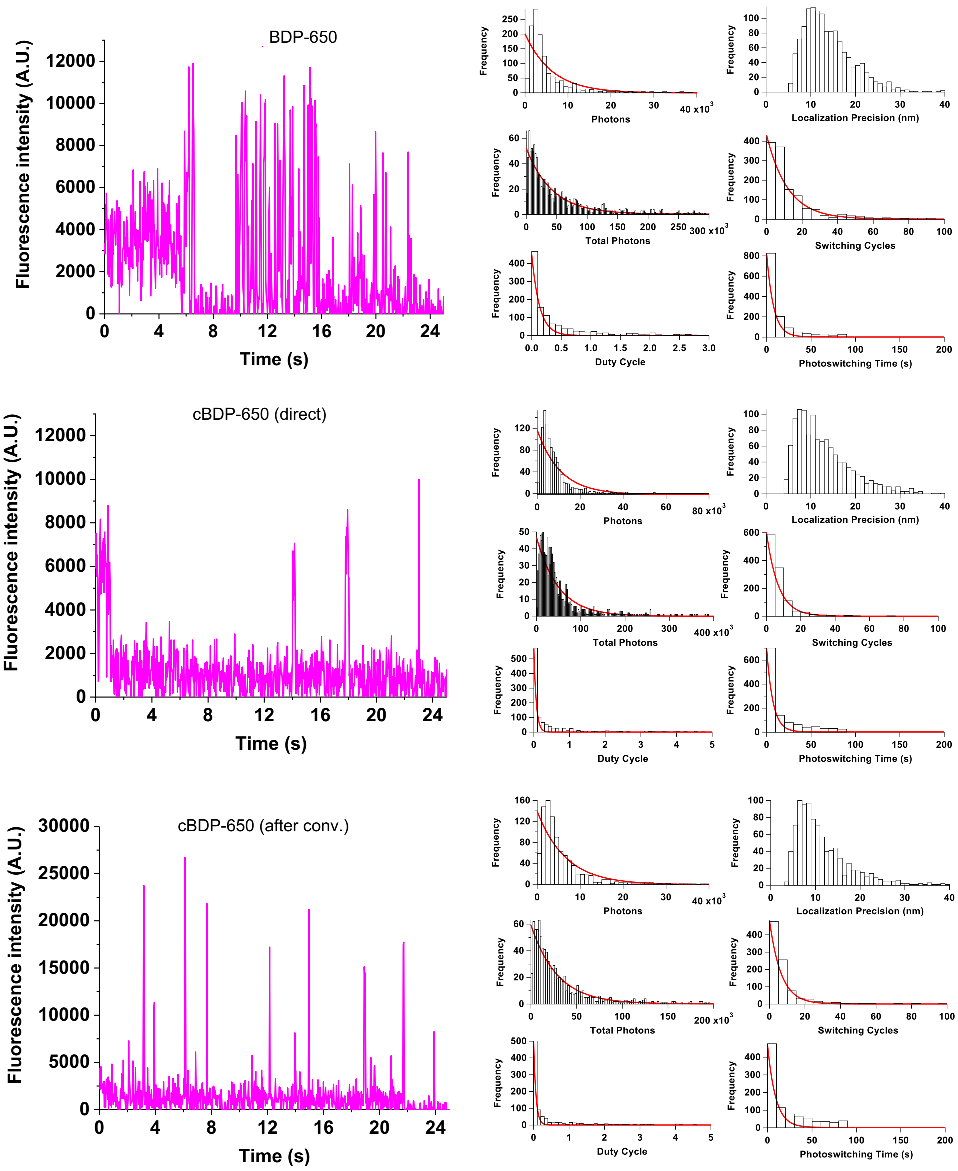


**Figure S18B. Single molecules and photoswitching properties analysis of BDP-650-Biotin.** “Photons” defines the number of photons detected by blinking event, it was calculated through the signal obtained knowing the setting of the camera (Gain, sensitivity). The “Total photons” defined the number of photons detected until the dye bleached. “Duty cycle” is the ratio between the time spent at the ON state and the time spent at the OFF state. “Localization precision” is the uncertainty of the localization, it is given by the thunderstorm plugin. “Switching cycle” is the number of ON/OFF cycles. The “photoswitching time” is the time during the dye switches before it bleaches.

**Algorithm used for the single molecule analysis on Igor Pro 8.**

```
function SMLM(wv,xx,yy,intensity,uncertainty,t_int)
wave wv,xx,yy,intensity,uncertainty
variable t_int
variable /G ON,OFF,bleach,pourcentage,kill,j,k,l,n,m,nbphotons
variable nb=numpnts(xx)
variable i,cx,cy,px,py
variable dd=dimSize(wv,2)
make /o/n=(dd) out
make /o/n=(nb)
Photons,Switching_Cycles,Total_photons,PhotoswitchingTime,DutyCycle
make /o/N=(1,6) Results
SetdimLabel 1,0,Photons,Results
SetdimLabel 1,1,Switching_Cycles,Results
SetdimLabel 1,2,Total_Photons,Results
SetdimLabel 1,3,localization_precision,Results
SetdimLabel 1,4,Photoswitching_Time,Results
SetdimLabel 1,5,Duty_Cycle,Results
OFF=0
ON=0
n=0
m=0
kill=0
wavestats /q wv
Mean_intensity=V_rms
for (i=0;i<nb;i+=1)
        string ii=num2str(i)
        string trace="bl_"+ii
        make /o/n=(dd) $trace
        wave out=$trace
        cx=xx[i]
        px=ScaleToIndex(wv,cx,0)
        cy=yy[i]
        py=ScaleToIndex(wv,cy,1)
        out[]=wv[px][py][p]
        wavestats /q out
        OFF=0
        ON=0
        n=0
        m=0
        bleach=0
        variable nc=numpnts(out)
        for(l=0;l<(nc);l+=1)
                if (out[l]< Mean_intensity*6 && n<1)
                        OFF=OFF+1
                        n=2
                        m=0
                        bleach=l
                elseif (out[l]> Mean_intensity*6 && m<1)
                        ON=ON+1
                        m=2
                        n=0
                endif
        endfor
        k=0
        j=0
        nbphotons=0
        for(l=0;l<bleach;l+=1)
                if (out[l]> Mean_intensity*6)
                        k=k+1
                        nbphotons=nbphotons+out[l]/V_max*intensity[l]
```

```
                Else
                        j=j+1
                endif
        endfor
        Switching_Cycles[i]=ON
        PhotoswitchingTime[i]=bleach*t_int*1e-3
        DutyCycle[i]=k/j
        Total_photons[i]=nbphotons
        Photons[i]=nbphotons/ON
        if (V_max<6*Mean_intensity || OFF<3)
                killWaves out
                Switching_Cycles[i]=nan
                PhotoswitchingTime[i]=nan
                DutyCycle[i]=nan
                Total_photons[i]=nan
                Photons[i]=nan
                uncertainty[i]=nan
                kill = kill + 1
        endif
        endfor
print "Number of molecules"
print nb-kill
print "Pourcent removed"
print kill/nb*100
```

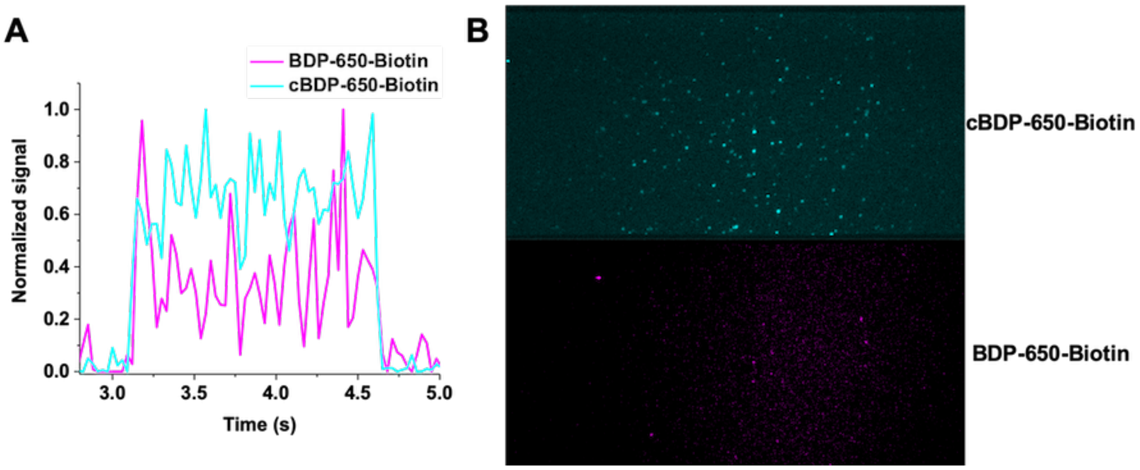


**Figure S19. Single molecules analysis of BDP-650-Biotin with dichroic mirror at 630 nm.** The dyes were irradiated with 561 nm laser (0.33 kW.cm$^{-2}$) and imaged with a 100× objective with integration time of 30 ms. The signal was split using a dichroic mirror at 630 nm to distinguish the emission from BDP-650-Biotin and cBDP-650-Biotin. (A) Example of a single molecule trace in both the initial and converted channels showing no obvious correlation. (B) Maximum projection of single molecule images over the time in both the initial (bottom one) and converted channels (top one).

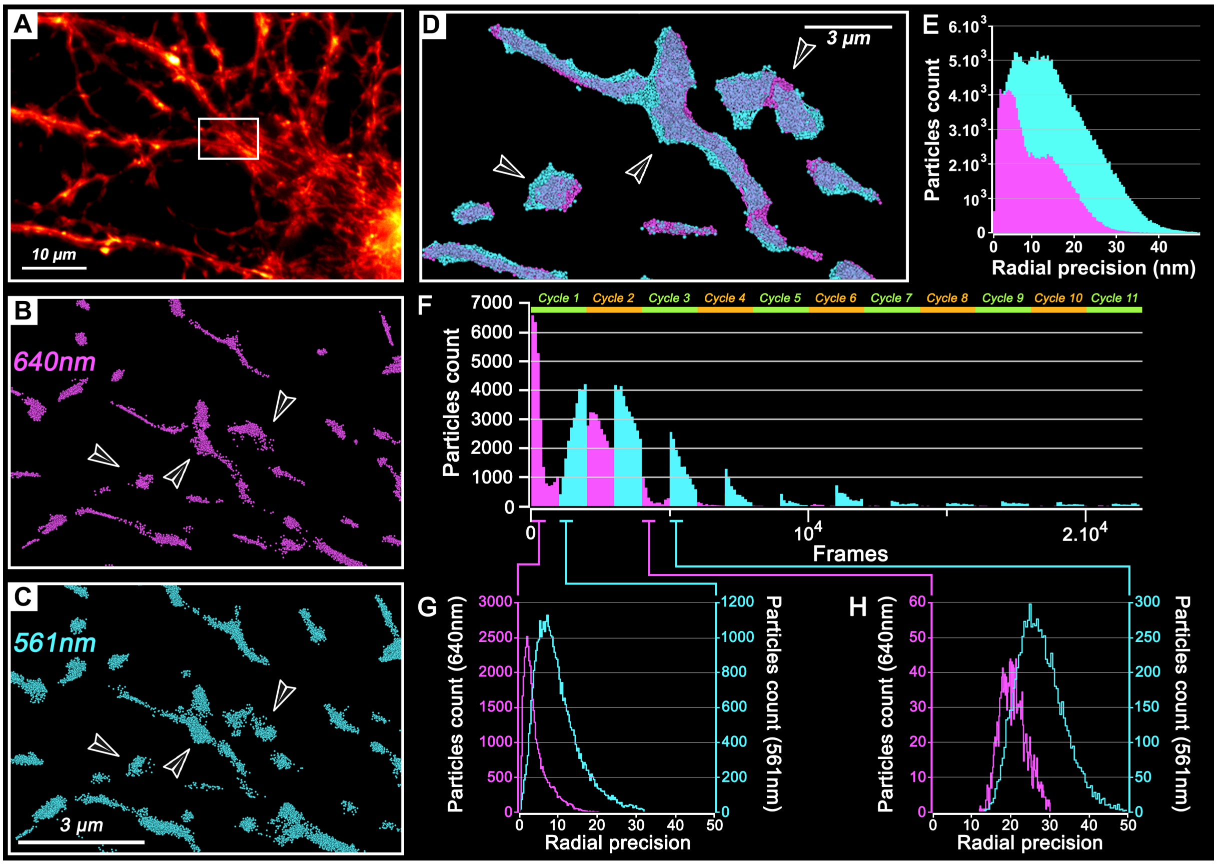


**Figure S20. Characteristics of both initial and converted form of BS-650-Mito in live neuron using interleaved 2 colors 3D-STORM imaging.** (A) Widefield fluorescence image of mitochondria in living 21 DiV-neuron labeled with BS-650-Mito over 11,000 frames. Fire LUT is color coding the intensity. White inset indicates magnification box in B-C. (B-C) Live-cell 3D-STORM imaging at 37°C in physiological buffer (Krebs-Ringer). The acquisition is composed of 11 times cycles of sequential illuminations with 640 nm (6.475 $kW.cm^{-2}$, 1,000 frames) followed by 561 nm (8.445 $kW.cm^{-2}$, 1,000 frames) at a frame rate of 10 ms. Single particles are represented as beads with a diameter of 50 nm. (B-D) Black arrows indicate mitochondrial structures which are shown in B, C and D. (B-G) Localizations retrieved within the 561 nm and 640 nm channels are shown in B for 561 nm converted form, and in C for 640 nm excitation of initial form. (D) DBScan segmentation of mitochondrial structures indicated with black arrows in B and C. Segmentation of non-converted and converted forms of BDP-650 labeling shows that they have an identical morphology. Both forms label the same cellular structures, indicating that the conversion does not affect its localization. (E) Radial precision of single particles localized during STORM imaging over 11,000 frames for non-converted (magenta) and converted (cyan) form of BDP-650. Mean radial precision is 11 nm (for 107,193 particles) of non-converted form and 20 nm (for 88,651 particles) of converted form. (F) Particles count over 22,000 frames; the 11 cycles are indicated with green and orange bars. Sequential illuminations are shown using a color code: 640 nm in magenta and 561 nm in cyan. After cycles 3, particles in 640 nm are not detected anymore whereas particles in 561 nm continued to blink. Graphic bars represent a bin of 100 frames. (G-H) Radial precision of single particles detected during the 500 first frames of cycle 1 (G) or cycle 3 (H). Mean radial precision is 12 nm for 8,003 particles of converted form in the first cycle, vs 5 nm for 22,544 particles of initial form. During the third cycle radial precision is decreased: 28 nm for 9,866 particles for the converted form, and 21 nm for 2,153 particles of non-converted form.

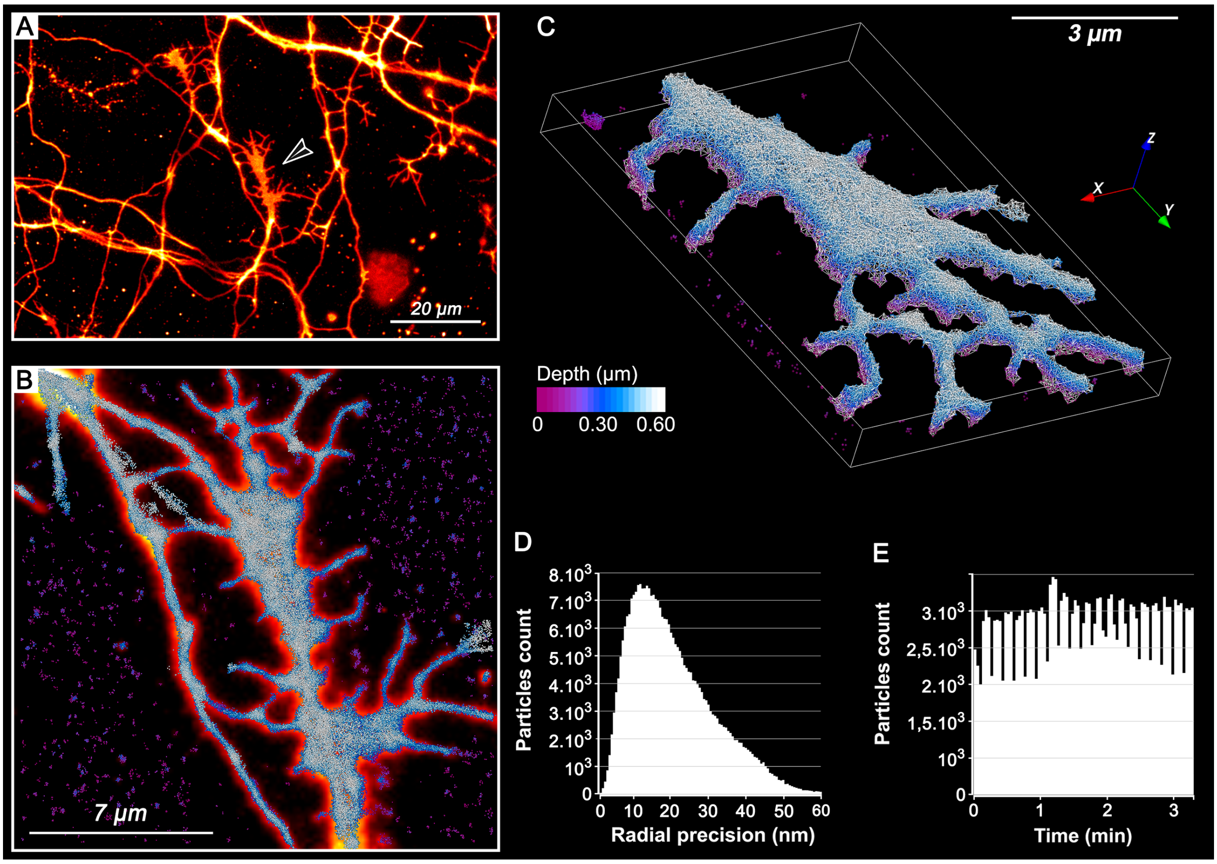


**Figure S21. 3D-STORM imaging of a neuronal growth cone in young live neuron using converted form cBDP-650-PM.** (A) Widefield fluorescence image of neuronal growth cone in living 7 DiV-neuron, labelled with BDP-650-PM. Colour code represents the intensity. Black arrow indicates the growth cone acquired in 3D-STORM in B. (B) Merge of widefield fluorescence image (fire LUT) and reconstructed 3D-STORM image (Ice LUT beads) corresponding to the growth cone indicated by the black arrow in A, at 37°C in physiological buffer (Krebs-Ringer). Single particles are represented as beads with a diameter of 50 nm. The colour code from purple to white represents the depth. 3D-STORM image using converted form cBDP-650 at 561 nm (10.185 kW.cm$^{-2}$) during 3 min without any illumination of 405 nm wavelength. (C) DBScan segmentation representing a part of the growth cone (white linear network) and single molecules detected using 3D-STORM (color gradient from purple to white is coding the depth). (D) Radial precision of single particles detected during 3D-STORM is 22 nm for 216,917 particles. (E) Particles count over 9,000 frames at a frame rate of 20 ms. Graphic bars represent a bin of 100 frames.

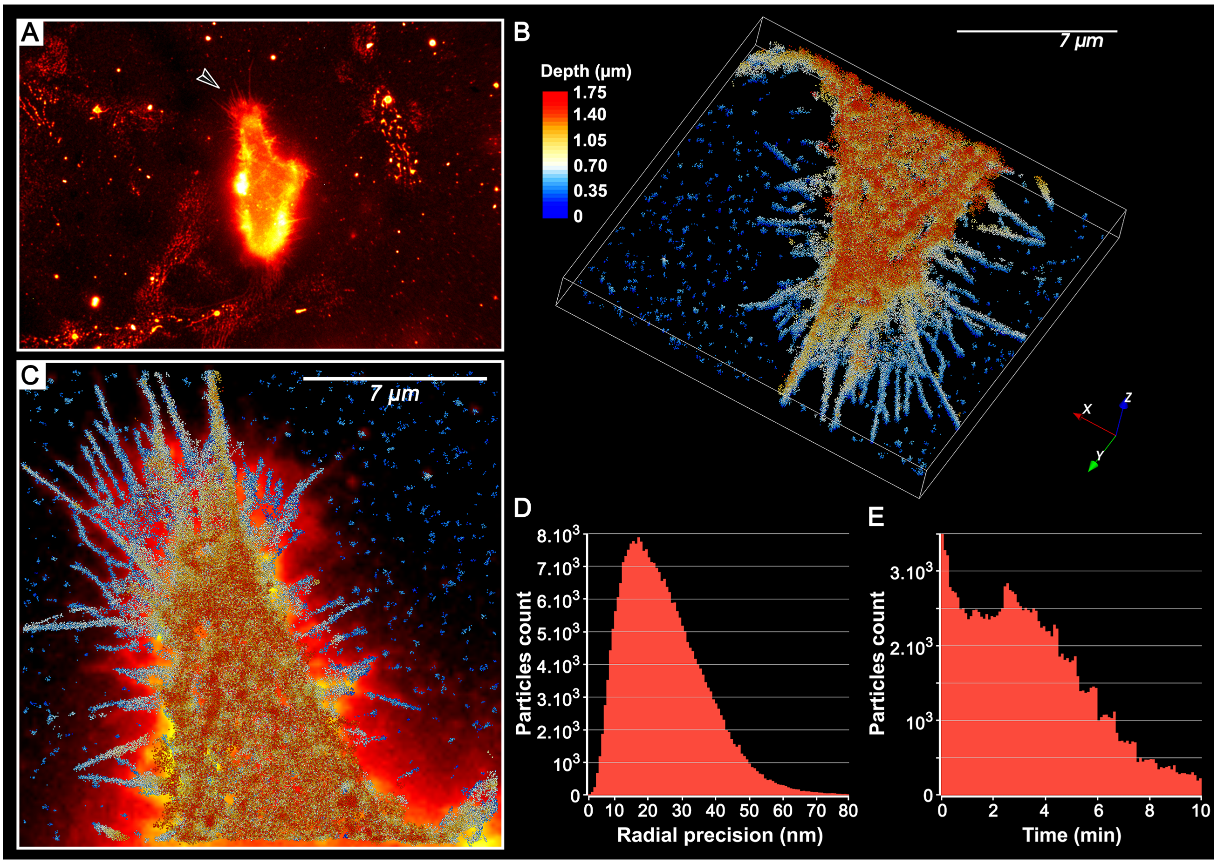


**Figure S22. 3D-STORM imaging of plasma membrane of an epithelial HeLa cell using converted form cBDP-650-PM.** (A) Widefield fluorescence image of Hela cell, labeled with BDP-650-PM. The intensity is color-coded with a fire LUT. Black arrow shows the filopodia acquired in 3D-STORM in B-C. (B-C) Live-cell 3D-STORM imaging, using cBDP-650-PM, at 37°C in physiological buffer (Krebs-Ringer) during 10 min 30 without any addition of 405 nm irradiation. The final 3D-STORM image is made up of 21Z containing 1,500 frames each. In totality, 3D-STORM reconstructions (B-C) are composed of 31,500 frames at a frame rate of 20 ms with 561 nm illumination (12.95 $kW.cm^{-2}$). Single particles are represented as 50 nm sphere. The color code from blue to red represents the depth. 3D reconstruction is presented in x,y axes (B) or in x,y,z view (C). (D) Radial precision of single particles, here mean precision is 25 nm for 164,751 particles. (E) Particles count over 31,500 frames at a frame rate of 20 ms. Graphic bars represent a bin of 100 frames.
